# A muscle hypertrophy-derived myokine reprograms the stromal vascular fractions differentiation towards thermogenic adipocytes in subcutaneous adipose tissue

**DOI:** 10.1101/2023.09.20.558627

**Authors:** Xiangping Yao, Xudong Mai, Ye Tian, Yifan Liu, Guanghui Jin, Ze Li, Shujie Chen, Xiaoshuang Dai, Xuejie Jin, Liujing Huang, Mulan Han, Zijing Fan, Lanling Xiao, Guihua Pan, Xiaohan Pan, Ziying Lin, Xiangmin Li, Jia Sun, Jingxing Ou, Hong Chen, Liwei Xie

**Affiliations:** Department of Endocrinology and Metabolism, Zhujiang Hospital, Southern Medical University, Guangzhou, 510280, China; State Key Laboratory of Applied Microbiology Southern China, Guangdong Provincial Key Laboratory of Microbial Culture Collection and Application, Guangdong Open Laboratory of Applied Microbiology, Institute of Microbiology, Guangdong Academy of Sciences, Guangzhou, 510070, China; School of Public Health, Xinxiang Medical University, Xinxiang, 453003, China; Department of Applied Biology and Chemical Technology, The Hong Kong Polytechnic University, Kowloon, Hong Kong SAR, China; BGI Institute of Applied Agriculture, BGI-Shenzhen, Shenzhen, China; Institute of Aging Research, Guangdong Provincial Key Laboratory of Medical Molecular Diagnostics, School of Medical Technology, Guangdong Medical University, Dongguan, China; Department of Hepatic Surgery and Liver transplantation Center of the Third Affiliated Hospital, Organ Transplantation Institute, Sun Yat-sen University; Organ Transplantation Research Center of Guangdong Province, Guangdong province engineering laboratory for transplantation medicine; Guangdong Provincial Key Laboratory of Liver Disease Research. Guangzhou 510630, China

**Author notes:** Corresponding authors Leading corresponding author: Liwei Xie. These authors contribute equally (Xiangping Yao, Xudong Mai, Ye Tian, Yifan Liu, Guanghui Jin).

**Keywords:** Bambi, Thbs4, muscle hypertrophy, stromal vascular fractions, beige adipocyte

## Abstract

Skeletal muscle plays a significant role in both local and systemic energy metabolism; however, the cross-talk between skeletal muscle and adipose tissue remains largely unexplored. In this study, we identify the HIF2α-Bambi-Thbs4 axis as a critical driver of muscle hypertrophy and metabolic improvement. Deletion of *Bambi* induces muscle hypertrophy and oxidative switching, thereby enhancing local and systemic metabolism under conditions of rodent chow and high-fat diets (HFD). Other than the metabolic regulation of skeletal muscle hypertrophy, we hypothesize that a portion of this improvement may be attributed to the metabolic reprogramming of the stromal vascular fraction (SVFs) of iWAT, promoting the development of beige adipocytes. Leveraging multi-omics approaches, we establish Thbs4 as a newly identified, muscle hypertrophy-derived myokine that serves as an essential regulator of this metabolic reprogramming. Thbs4 is upregulated in skeletal muscle during hypertrophy and aerobic exercise, localizes to the iWAT membrane, and facilitates the transformation of white adipocytes into beige adipocytes under conditions such as cold exposure, muscle hypertrophy, or aerobic exercise. Moreover, overexpression of Thbs4 elevates its plasma levels and leads to the transformation of white to beige adipocytes, therefore offering long-term protection against HFD-induced metabolic disorders. Conversely, *Thbs4* knockout (*Thbs4*-KO) disrupts the metabolic reprogramming of SVFs and exacerbates metabolic syndromes induced by an HFD, mirroring the observed decline in Thbs4 levels in aged mice. Therefore, our findings underscore Thbs4’s potential as both a long-term metabolic protective factor and a therapeutic target for the treatment of metabolic disorders.

**Graphical Abstract:** 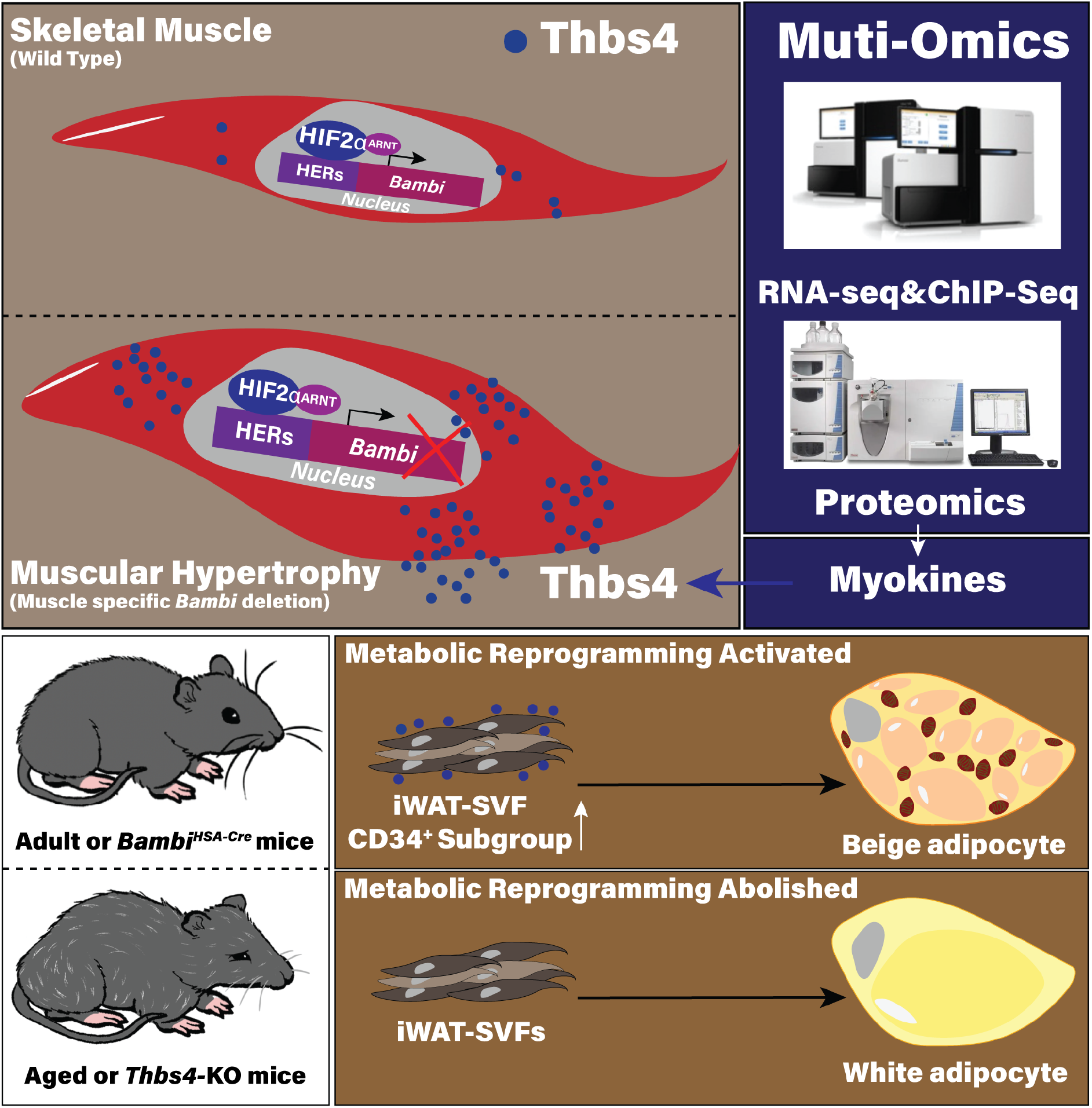

**Highlights:**

- *Bambi*, a direct target gene of HIF2α in skeletal muscle, when deleted, triggers muscle hypertrophy and the transformation of white into beige adipocytes, improving systemic energy metabolism.
- The metabolic reprogramming of stromal vascular fractions (SVFs) contributes to the observed improvements in energy metabolism and adipocyte hyperplasia.
- Thbs4, a novel myokine produced in skeletal muscle during hypertrophy and aerobic exercise, plays a key role in the transformation of white into beige adipocytes.
- Overexpression of Thbs4 provides enduring protection against high-fat diet (HFD)-induced metabolic disorders, demonstrating its potential as a therapeutic target.
- *Thbs4* deletion abrogates the beneficial metabolic reprogramming of SVFs and exacerbates HFD-induced metabolic syndromes, further underlining its crucial role in metabolic regulation.

## Introduction

Skeletal muscle, constituting approximately 40-50% of the total body weight in healthy individuals, and plays pivotal roles in daily physical activity^1,2^. Physical training delivers a wide range of benefits on host health through various mechanisms, such as weight loss, enhanced cardiovascular function, and preservation of muscle mass, strength and oxidative fiber type percentage^3–6^. These alterations in skeletal muscle due to exercise training, via inter-organ crosstalk, are associated with systemic benefits to distal organ physiology and metabolism. Recent studies have established that physical training-induced and skeletal muscle-derived metabolites or secretory factors, such as β-aminoisobutyric acid (BAIBA)^7^, GABA (γ-aminobutyric acid)^7^, IL6^8–10^, and irisin^11^, exert positive impacts both locally and systemically on glucose, lipid and energy metabolism^12^. Conversely, under pathophysiological conditions like sarcopenia and cachexia, an imbalance between protein degradation and synthesis is closely linked with muscle atrophy, which often coincides with chronic metabolic disorders^13^. Therefore, maintaining an appropriate amount of muscle mass during development and aging is indispensable for systemic metabolic homeostasis.

Postnatal skeletal muscle growth, regeneration, and hypertrophy are derived from satellite cells (SCs)^14^. Quiescent satellite cells (QSCs), positioned beneath basal lamina and outside the plasma membrane of myofibers, express the transcription factor paired box protein 7 (Pax7)^15^. QSCs are activated upon external stimulation, such as injury or physical training, followed by the expression of a family of myogenic regulatory factors (MRFs) that include myogenic differentiation 1 (MyoD), myogenin (MyoG) and myogenic factor 5 (Myf5)^14^. In addition to the regulation by MRFs, SC physiology and homeostasis are also precisely regulated by niche environments, such as oxygen^16^ and nutrients^17^. Recent studies have uncovered a group of transcription factors involved in regulating SC physiology, which in turn affect skeletal muscle growth and hypertrophy. These include hypoxia inducible factors (HIFs)^18,19^, and Yin Yang 1(YY1)^20^. HIFs are an oxygen-sensitive transcriptional factor complex, composed of α- and β-subunit^21,22^. HIF1α and HIF2α interact with a spectrum of factors, involved in regulating the cell cycle and metabolism, such as Sp1^23^ and YY1^24^. Among these factors, transient deletion of *Hif2α* or inhibition via small molecule inhibitor promotes SC’s proliferation, muscle regeneration and hypertrophy, while long-term deletion of HIF2α results in a reduction in the number of SCs and eventually a defect in regeneration ^18^. Thus, identifying HIF2α downstream targets to phenocopy the muscle hypertrophy remains poorly understood and require further exploration to identify novel target that could avoid the side-effect of long-term deletion.

Skeletal muscle not only locally regulates glucose and lipid metabolism but also communicates with distal organs or tissues, such as the liver, brain and adipose tissue, through cytokines, myokines and metabolites during exercise or due to muscle hypertrophy^6,25–27^. Recent findings have demonstrated that myokines exert their regulatory effects on different subtypes of adipose tissues to induce metabolic benefits in activating the thermogenesis of brown adipocytes or the transformation of inducible beige adipocytes^28^. The best-known function of brown and beige adipocytes is the thermogenic capacity, characterized by multilocular lipid droplets, a large number of mitochondria in the cytoplasm and the selective expression of peroxisome proliferator-activated receptor gamma coactivator 1α (PGC1α) and uncoupling protein 1 (Ucp1) ^28^. These phenotypes of thermogenic adipocytes couple with energy expenditure and glucose homeostasis in healthy individuals. In recent decades, several essential transcriptional regulators, cascades, and epigenetic regulators have been identified, such as PR domain zinc-finger protein 16 (Prdm16) ^29,30^, Prdm16-binding partner CCAAT/enhancer-binding protein-β (C/EBPβ) ^31^, EBF Transcription Factor 2 (Ebf2)^32^, Peroxisome proliferator-activated receptor ψ (Pparψ) ^33,34^, and Euchromatic Histone Lysine Methyltransferase 1 (EHMT1) ^35^, which participate directly in the regulation of the development and thermogenic function of brown and beige adipocytes. Besides the internal regulators, several exercise-induced myokines with beigeing effects have been identified, such as IL6^36^, irisin^11^, meteorin-like (Metrnl)^37^, GDF15^38^ and BAIBA^7^. Exercise has a spectrum of benefits for healthy individuals but it is impractical for disabled or aged population. Thus, it is urgent to identify new biomarkers involved in muscle hypertrophy or muscle-derived myokines that can recapitulate the beneficial effects of exercise on systemic glucose and lipid metabolism. This study focused on elucidating a gene, *Bambi* linked to muscle hypertrophy and identifying a corresponding secretory factor that influences systemic metabolism. The skeletal muscle-specific deletion of the *Bambi* gene resulted in muscle hypertrophy and an enhancement of metabolic performance, even under high-fat diet conditions. This change was attributed to the metabolic reprogramming of stromal vascular fractions (SVFs) into thermogenic adipocytes. The research pinpointed Thbs4, a myokine, which was significantly upregulated and secreted in mice undergoing *Bambi* gene deletion or aerobic exercise. Elevated Thbs4 levels corresponded to its concentration predominantly in specific adipose tissues but not in other organs. In *Thbs4*-knockout mice, the severity of metabolic syndrome exacerbated. A thorough examination of muscle samples revealed a chromatin-opening event on particular factors, such as MEFs, MyoD and MyoG. This finding correlates with the upregulation of Thbs4 expression during SC activation, myoblast proliferation, differentiation and muscle growth/hypertrophy, with secretion through a canonical endoplasmic reticulum pathway. The study, therefore, illuminates the role of a hypertrophy-induced myokine that bolsters adipocyte thermogenesis, suggesting potential therapeutic applications for enhancing energy metabolism in combating age-related or metabolic syndromes.

## Results

### *Bambi* is a direct target gene of HIF2α in skeletal muscle

HIF2α is a dominant oxygen-sensing transcription factor expressed in both SCs and skeletal muscles^18,19,39^. Previously, we demonstrated that transient inhibition of HIF2α through a small molecule inhibitor accelerated injury regeneration and promoted muscle hypertrophy in adult *C57BL/6J* mice without causing a significant change in fiber type^18^. However, downstream target genes and molecular mechanisms mediated by HIF2α in regulating skeletal muscle hypertrophy remain obscure. Further investigation revealed that other than Pax7^+^ satellite cells, HIF2α is also expressed and localized in myonuclei both in tibialis anterior (TA) muscle cryosections (Figure S1A) and in single myofibers isolated from extensor digitorum longus (EDL) muscle as shown before^18^. We thus hypothesized that HIF2α in myonuclei may also play a role in skeletal muscle growth following injury and regeneration, as intramuscular injection of HIF2α small molecule inhibitor may affect HIF2α protein in entirety of muscle cells, including SCs and their surrounding cells^18^. Leveraging the INTACT-nucleus isolation approach established and optimized previously^18^, PFA-fixed GFP^+^ myonuclei from *HSA-Cre;INTACT* mice were precipitated, purified and sonicated to generate 300-400 bp DNA fragments (Figure S1B) for HIF2α chromatin immunoprecipitation and high-throughput sequencing (ChIP-seq). The bioinformatic analysis identified 24571 peaks, which were annotated to be associated with 10329 genes (Figure S1C). Among these DNA fragments, ∼5.7% were enriched in the promoter region (Figure S1C-D). Gene Ontology (GO: biological process, cellular component, and molecular function) and Kyoto Encyclopedia of Genes and Genomes (KEGG) pathway analyses revealed these genes were associated with HIF2α-mediated hypoxic signaling pathway (Figure S1E). These genes are involved in macronutrient metabolism (*e.g.*, pyruvate, and amino acids), divalent metal ion transport, the HIF signaling pathway and muscle tissue development (Figure S1E). Motif analysis identified a group of HIFs-related co-activator and myogenic differentiation binding sites, such as HIF and associated co-factors like Sp1^23^, CTCF, and MyoD/MyoG (Figure S1F). Among them, *Bambi* could be an intriguing HIF2α target that is involved in muscle growth and development^40,41^. Direct binding of HIF2α to the *Bambi* promoter was further confirmed by HIF2α ChIP-qPCR, presenting ∼6- and 4-fold enrichment in the HIF2α pull-down fragment for *Bambi* and Cav1^18,42^, respectively, while there was no enrichment on the negative control gene *Tubulin* (Figure S1G). The *Bambi* promoter flanking the 1 kb region with five putative hypoxia response elements (HREs) in the luciferase construct was transfected into 293T cells. CoCl_2_ treatment (to mimic hypoxia) was known to stabilize HIFs^43,44^, and led to a 1.5-fold induction of luciferase activity (Figure S1H). Co-transfection of either wild-type or mutant HIF1α and HIF2α expression vectors with the *Bambi* promoter construct was performed in 293T cells, and only HIF2αTM (carrying P405A, P530V and N851A mutations of murine HIF2α) activated *Bambi* promoter activity but not HIF1α or wild type HIF2α (Figure S1I-J). The truncated constructs (0.68, 0.47, 0.33 and 0.3 kb) co-transfected with the HIF2αTM vector preserved luciferase activity similar to that of the 1 kb promoter construct, indicating that HRE at the proximal region of 0.33kb may be essential to HIF2α-mediated induction (Figure S1K). Thus, HRE located at the 5’ UTR on the 0.3 kb construct was mutated, which completely abolished HIF2α-mediated induction of luciferase activity (Figure S1L). These findings reveal that *Bambi* is a novel HIF2α directly regulated gene.

From this point, we also examined the association of HIF2α-Bambi axis in *in vivo* and i*n vitro* model. The muscle-specific overexpression of *Hif2α* (*Hif2α^LSL/HSA-Cre^* mice) was established, and the data showed that in skeletal muscles (including TA, Sol, EDL and Gas), HIF2α overexpression upregulated *Bambi* and *Vegfa* mRNA expression (Figure S2A-D). HIF2α overexpression also increased Bambi protein levels (Figure S2E). In both C2C12 myoblasts and primary myoblasts, HIF2α overexpression, achieved via adenovirus infection, significantly upregulated *Bambi* expression and downregulated myogenesis-related gene expression, like MyoG, myosin heave chains (MHCs, Myh7, 2, 1, 4), followed by the inhibition of myogenic differentiation (Figure S2 F-K). These *in vivo* and *in vitro* findings echo previous observations that transient transfection and overexpression of HIF2α inhibits the differentiation of C2C12 and primary myoblasts. Therefore, it is necessary to assess whether *Bambi*-deletion could recapitulate HIF2α function on skeletal muscle growth and hypertrophy.

### Muscle-specific deletion of *Bambi* promotes muscle growth and induces muscle hypertrophy

We crossed *Bambi^fl/fl^* mice with *HSA-Cre* mice, generating skeletal muscle-specific *Bambi* knockout mice (*Bambi^HSA-Cre^*, Figure S3A and S3H). Between *Bambi^HSA-Cre^* and *Bambi^fl/fl^* mice, the weights of TA, EDL and iWAT from *Bambi^HSA-Cre^* mice were significantly heavier than those from *Bambi^fl/fl^* mice, with correspondent heavier body weight but no change on food intake (Figure S3B-D, G). These data were further supported by DEXA scanning result of lean and fat mass (Figure S3E-F). The increased muscle weight in *Bambi^HSA-Cre^* mice was likely due to the activation and myogenic differentiation of SCs that express MyoD, which was confirmed by single myofiber isolation from EDL muscle and immunostaining of Pax7 and MyoD. *Bambi* deletion dramatically increased MyoD^+^ satellite cells on single myofibers (Figure 1A-C). *Pax7* and *Myod1* mRNA and protein expression were also induced in the TA muscle of *Bambi^HSA-Cre^*mice (Figure 1D and Figure S3I). Meanwhile, *Bambi* deletion also led to a shift in fiber type to slow-twitch, with elevated expression of *Myh7* and *Myh2* (Figure 1D). A similar expression pattern on *Pax7*, *Myod1*, *Myog* and *Myh7* were observed in Sol muscle (Figure S3J). This was followed by a significantly increased percentage of MyHC I^+^ and MyHC IIA^+^ myofibers on TA cryosections (Figure 1E-F). This is consistent with our previous finding that *Bambi* expression is negatively associated with oxidative fiber type in the soleus^40^. Expression of the fast-twitch fiber-related genes *Myh1* and *Myh4* was not affected, which corresponded to the unchanged percentages of MyHC IIB^+^ and MyHC IIX^+^ myofibers in the TA muscle (Figure 1D and Figure S3K).

**Figure 1.**
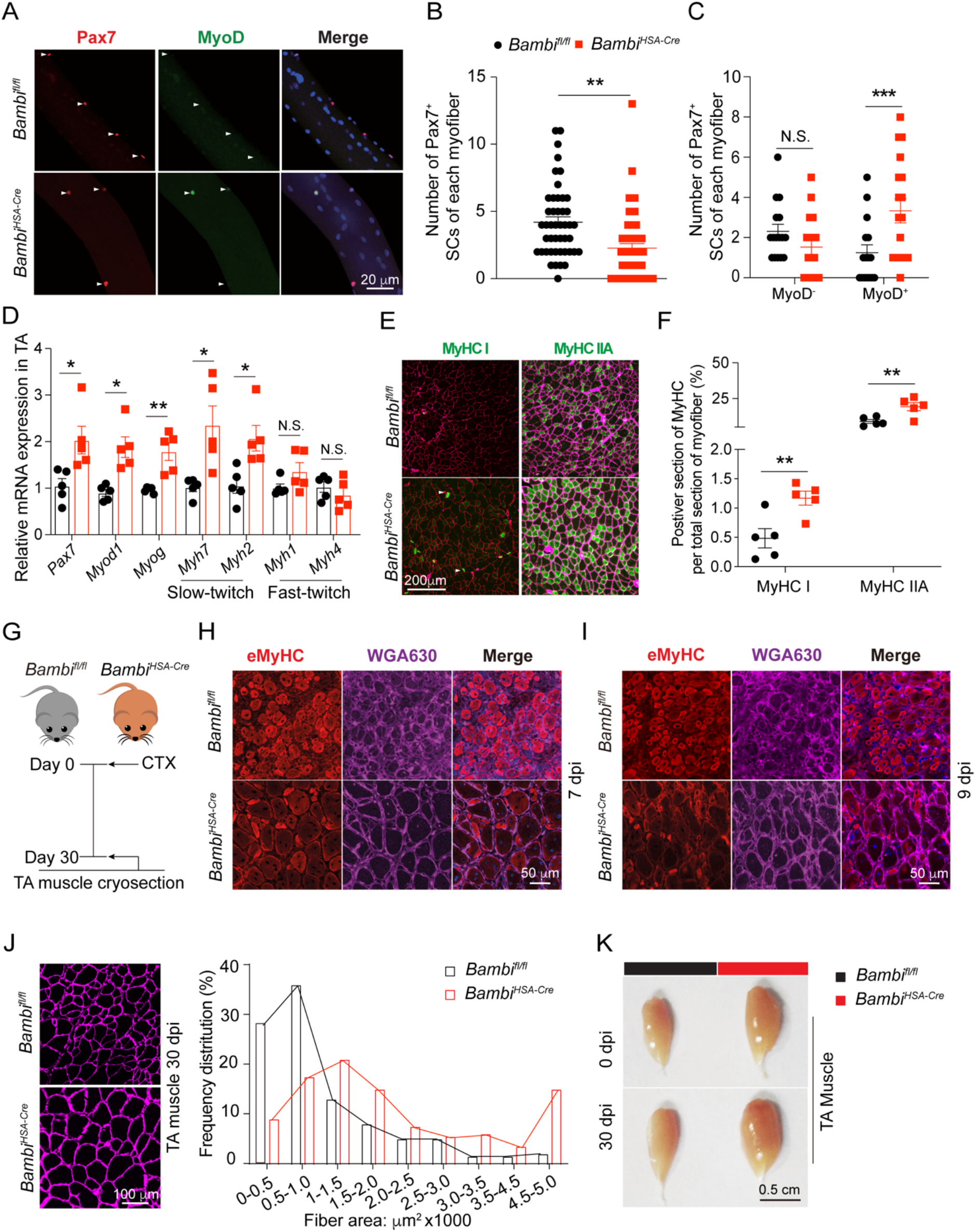
Muscle-specific deletion of *Bambi* induces muscle hypertrophy. (A) Representative images of immunostaining of Pax7 (Red), MyoD (Green) and DAPI (Blue) of satellite cells in single myofiber isolated from EDL muscle of *Bambi^fl/fl^* and *Bambi^HSA-Cre^*mice (n >100 myofibers from 6 mice/group). (B) Number of Pax7^+^ SCs of each myofiber from the *Bambi^fl/fl^* and *Bambi^HSA-Cre^* mice. (C) Number of Pax7^+^/MyoD^-^ (left) and Pax7^+^/MyoD^+^ (right) cells in each myofiber of *Bambi^fl/fl^* and *Bambi^HSA-Cre^* mice. (D) qPCR analysis of muscle-related gene expression in TA muscle samples from *Bambi^fl/fl^* and *Bambi^HSA-Cre^*mice (n=5 mice/group). (E) Representative images of the immunostaining of MyHC I (Green), MyHC IIA (Green) and laminin B2 (Purple) in TA cryosection of *Bambi^fl/fl^* and *Bambi^HSA-Cre^* mice (n=5 mice/group). (F) The percentage of MyHC I^+^ and MyHC IIA^+^ myofibers per section. (G) Timeline characterizing skeletal muscle regeneration experiment between *Bambi^fl/fl^*and *Bambi^HSA-Cre^* mice. (H-I) Representative images of eMyHC (Red) and WGA630 (Purple) immunofluorescence in TA muscles 7 dpi and 9 dpi (n=6 mice/group). (J) Representative images of WGA630 (Purple) (left) immunostaining in TA muscle sections 30 days post injury, and the frequency distribution of fiber cross-section area (right) between *Bambi^fl/fl^* and *Bambi^HSA-Cre^*mice. (K) Representative images of TA muscles from *Bambi^fl/fl^*and *Bambi^HSA-Cre^* mice before and 30 days post injury. N.S., not significant, \**p* < 0.05, \*\**p* < 0.01, and \*\*\**p* < 0.001, by two-sided Student’s t test. Data represent the mean ± standard error of the mean.

To further delve into the physiological function of *Bambi* during muscle regeneration, TA muscle injury was induced using intramuscular injection of cardiotoxin (CTX) (Figure 1G). *Bambi* deletion in skeletal muscle couples with more activated SCs and accelerated regeneration (Figure 1A and 1D). At 7- and 9-day post injury (dpi), the TA of *Bambi^HSA-Cre^* mice had fewer eMyHC^+^ myofibers and more myofibers at larger sizes (Figure 1H-J, S3L). *Bambi^HSA-Cre^* mice gained extra body weight, along with larger and heavier TA muscle at 30 dpi (Figure 1K, S3M-N), but exerted no obvious difference in food intake (Figure S3O). The mass and ratio of tissue (TA muscle and iWAT)/BW were increased, except for the soleus, eWAT and iBAT (Figure S3P). Muscle hypertrophy continuously occurred in *Bambi^HSA-Cre^* mice at 30 dpi, with an enlarged fiber size and upregulated myogenic and oxidative fiber type-related gene expression (Figure 1J-K and Figure S3Q). These results strongly suggest that the deletion of *Bambi* gene promotes skeletal muscle growth and fiber type alteration, likely through *Bambi*-mediated modulation of the skeletal muscle physiology.

### *Bambi* deletion-mediated muscle hypertrophy promotes systemic energy metabolism

Deletion of *Bambi* resulted in SC activation, TA muscle hypertrophy, and a fiber-type shift favoring oxidative phosphorylation. To determine the molecular mechanisms that underlie these phenotypes, we performed a transcriptomic analysis of TA muscle from *Bambi^fl/fl^*and *Bambi^HSA-Cre^* mice before and after injury/regeneration. A distinct subset of genes was differentially expressed in TA muscle between *Bambi^HSA-Cre^* and *Bambi^fl/fl^* mice before (Figure S4A-B) and 30 days post injury (Figure S4C-D). We particularly focus on the differentially expressed genes (DEGs) that were associated with muscle growth and hypertrophy in TA muscle 30 days post injury (Figure S4E). In particular, the DEGs for TA muscle post-regeneration presented even dramatically changes than before injury between *Bambi^HSA-Cre^* and *Bambi^fl/fl^* mice, as it was analyzed by gene clustering and principal component analysis (PCA) (Figure 2A-B). For a subset of DEGs in TA muscle post injury, gene ontology analysis (biological process) revealed that upregulated genes were associated with ATP and ADP metabolism (Figure S4E) and functional enrichment analyses against KEGG pathways prioritized the significantly changed pathways, *e.g.*, thermogenesis, insulin, and the AMPK signaling pathway (Figure 2C). This was further confirmed by gene set enrichment analysis (GSEA) and gene expression associated with the ATP metabolism process and skeletal muscle growth by heatmap (Figure 2D-F). At the systemic level, the indirect calorimetry data indicated that *Bambi^HSA-Cre^* mice exhibited higher VO_2_ (Figure S4F), VCO_2_ (Figure S4G) and energy expenditure (Figure 2G-I) than those of *Bambi^fl/fl^* mice before and 30 days post injury (Figure 2J). However, these indices were not significantly different between non-injured and post-injured *Bambi^HSA-Cre^* mice (Figure 2I, S4F, S4G). Differentially expressed energy metabolism-related genes and increased systemic metabolism corresponded to the increased number of mitochondria in transmission electron microscopy imaging for *Bambi^HSA-Cre^* mice (Figure 2K), and an induction of PGC-1α protein levels, a transcriptional coactivator of mitochondrial biogenesis in cells, was also markedly enhanced in the TA muscle of *Bambi^HSA-Cre^* mice after 30 dpi (Figure S3H). Glucose and insulin tolerance assays (GTT and ITT) showed significant improvement in glucose tolerance and insulin sensitivity of *Bambi^HSA-Cre^* mice (Figure 2L-M), especially significant improvement in insulin sensitivity of *Bambi^HSA-Cre^* mice 30 days post injury (Figure 2M), suggesting that unexplored mechanisms may exist. Together, these findings demonstrate that *Bambi* ablation in skeletal muscle leads to significant metabolic alterations in skeletal muscle and whole-body metabolism.

**Figure 2.**
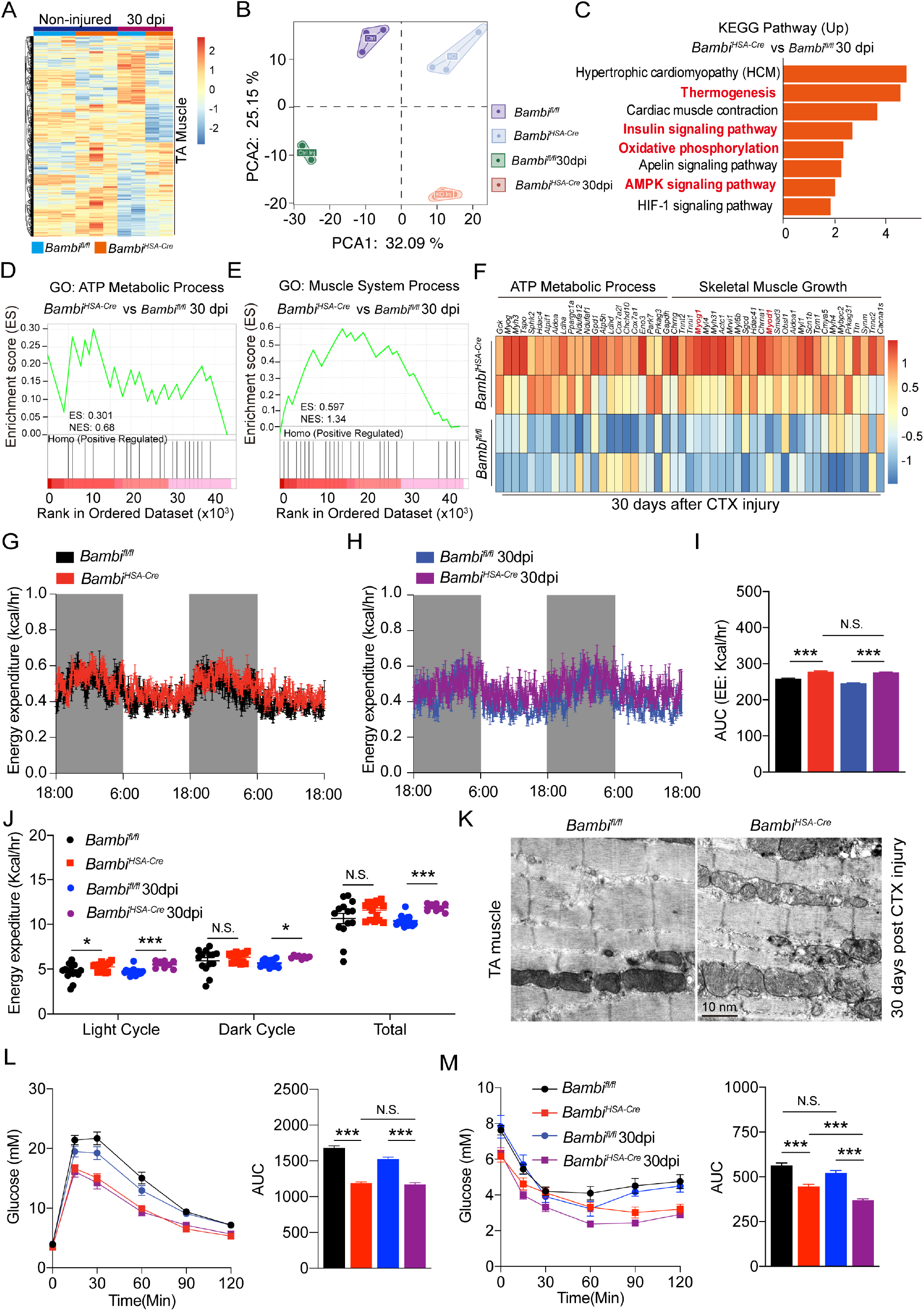
*Bambi* deletion-induced muscle hypertrophy promotes systemic energy metabolism. (A) Heatmap showing the mRNA expression profile in TA muscle between *Bambi^fl/fl^*and *Bambi^HSA-Cre^* mice before and 30 days post injury. (B) Principal component analysis (PCA) of the top 100 DEGs. (C) KEGG pathway enrichment analysis of upregulated genes. (D-E) GSEA of the ATP metabolic process (GO) and muscle system process (GO). (F) Heatmap of ATP metabolic process and skeletal muscle growth-related gene expression in TA muscle 30 days post injury. (G-H) Average energy expenditure (E.E.) was monitored over a 48-hr period for *Bambi^fl/fl^* and *Bambi^HSA-Cre^*mice before (G) and 30 days post injury (H). (I) AUC of E.E. (J) Average energy expenditure in light, dark and total cycles between *Bambi^fl/fl^* and *Bambi^HSA-Cre^* mice (n=4 mice/group for two independent repeat). (K) Representative images of transmission electron microscopy of TA muscle from *Bambi^fl/fl^* and *Bambi^HSA-Cre^* mice 30 days post-CTX injury (n=3 mice/group). (L-M) The glucose tolerance test (GTT, L) and insulin tolerance test (ITT, M) of *Bambi^fl/fl^* and *Bambi^HSA-Cre^* mice (n= 6 mice/group). N.S., not significant, \**p* < 0.05, \*\**p* < 0.01, and \*\*\**p* < 0.001, by two-sided Student’s t test. Data represent the mean ± standard error of the mean.

### *Bambi* deficiency in skeletal muscle activates the thermogenic capacity of iWAT

In addition to skeletal muscle, the thermogenesis of brown and beige adipocytes is coupled with enhanced systemic metabolism and increased energy expenditure^28^. To determine whether the effect of *Bambi* deficiency on energy expenditure is attributable to induced thermogenesis of subcutaneous adipose tissue and increased heat production, leading to further improvement on insulin sensitivity 30 days post injury, we next analyzed the gene expression profile in iWAT at two stages between *Bambi^fl/fl^*and *Bambi^HSA-Cre^* mice. Consistent with increased mitochondrial biogenesis and energy metabolism in skeletal muscle, a distinct subset of genes was differentially also expressed in iWAT between *Bambi^HSA-Cre^* and *Bambi^fl/fl^* mice before and 30 days post injury (Figure S5A-D). The KEGG pathway, GO (biological process) and GSEA analyses revealed that thermogenesis, oxidative phosphorylation and mitochondria-related biological functions were enriched (Figure 3A, S5E-G). qPCR confirmed the induction of thermogenesis (*Prdm16*, *Ppargc1α*, *Ucp1*, *Cox7a1*, *Cox8b*, and *Cidea*) and beige-related genes (*Gatm*, and *Ebf2*) in iWAT of *Bambi^HSA-Cre^* mice (Figure 3B). Meanwhile, adipogenesis-related genes, such as *Srebp1* and *Pparγ*, were also increased without change on expression of *Fas* and *Adipoq* (Figure S5H). Meanwhile, protein expression levels of Ucp1, p-HSL, p-AMPK, ATGL and Glut4 were also markedly increased in iWAT of *Bambi^HSA-Cre^*mice (Figure 3C). Glut4 protein expression was increased and enriched on the membrane of iWAT in *Bambi^HSA-Cre^* mice (Figure 3D). Consistent with thermogenic gene expression, immunohistochemistry (IHC) revealed an increased Ucp1-positive area around multilocular lipid droplets in the iWAT of *Bambi^HSA-Cre^*mice (Figure 3D). Through transmission electron microscopy imaging, we observed an increased number of mitochondria (M) and smaller lipid droplets in the iWAT of *Bambi^HSA-Cre^* mice (Figure 3E). We further assessed the lipid profile of *Bambi^fl/fl^* and *Bambi^HSA-Cre^* mice, exhibiting a high level of HDL-c in the circulating plasma of *Bambi^HSA-Cre^* mice, but no changes in TG, TC, FFA or LDL-c were observed (Figure S5I). ALT and AST levels and the AST/ALT ratio, both of which are the indicators of liver function, demonstrated that AST levels were clearly reduced and that the ratio was lower in *Bambi^HSA-Cre^* mice, indicating improved on systemic energy metabolism and liver function in *Bambi^HSA-Cre^* mice (Figure S5J-K). To further delve into the contribution of the beigeing of iWAT to the improvement of systemic metabolism, the metabolic indices, including VO_2_ (Figure 3F-H) and heat production (Figure 3I-K) were recorded at room temperature (R.T.) and under cold challenge for both *Bambi^fl/fl^* and *Bambi^HSA-Cre^* mice. At R.T., VO_2_ and heat production were only significantly higher in light cycle, while with acute cold challenge, the VO_2_ and heat product were dramatically induced at both light and dark circle, suggesting cold-induced non-shivering thermogenesis from adipose tissue contributed to the improved basal metabolism. H.&E. and IHC of PGC1α, Ucp1 and Glut4 results demonstrated that in the area of multilocular lipid droplets, there were more beigeing area with higher expression of PGC1α, Ucp1 and Glut4 under both R.T. and cold challenge (Figure 3L). These data establish a link between muscle hypertrophy and beigeing of subcutaneous adipose tissue, both of which contribute to the boosted basal metabolism and improved systemic metabolic indices. However, the exact regulatory mechanisms that contribute to the induced thermogenesis and beigeing in *Bambi^HSA-Cre^* mice remains unclear.

**Figure 3.**
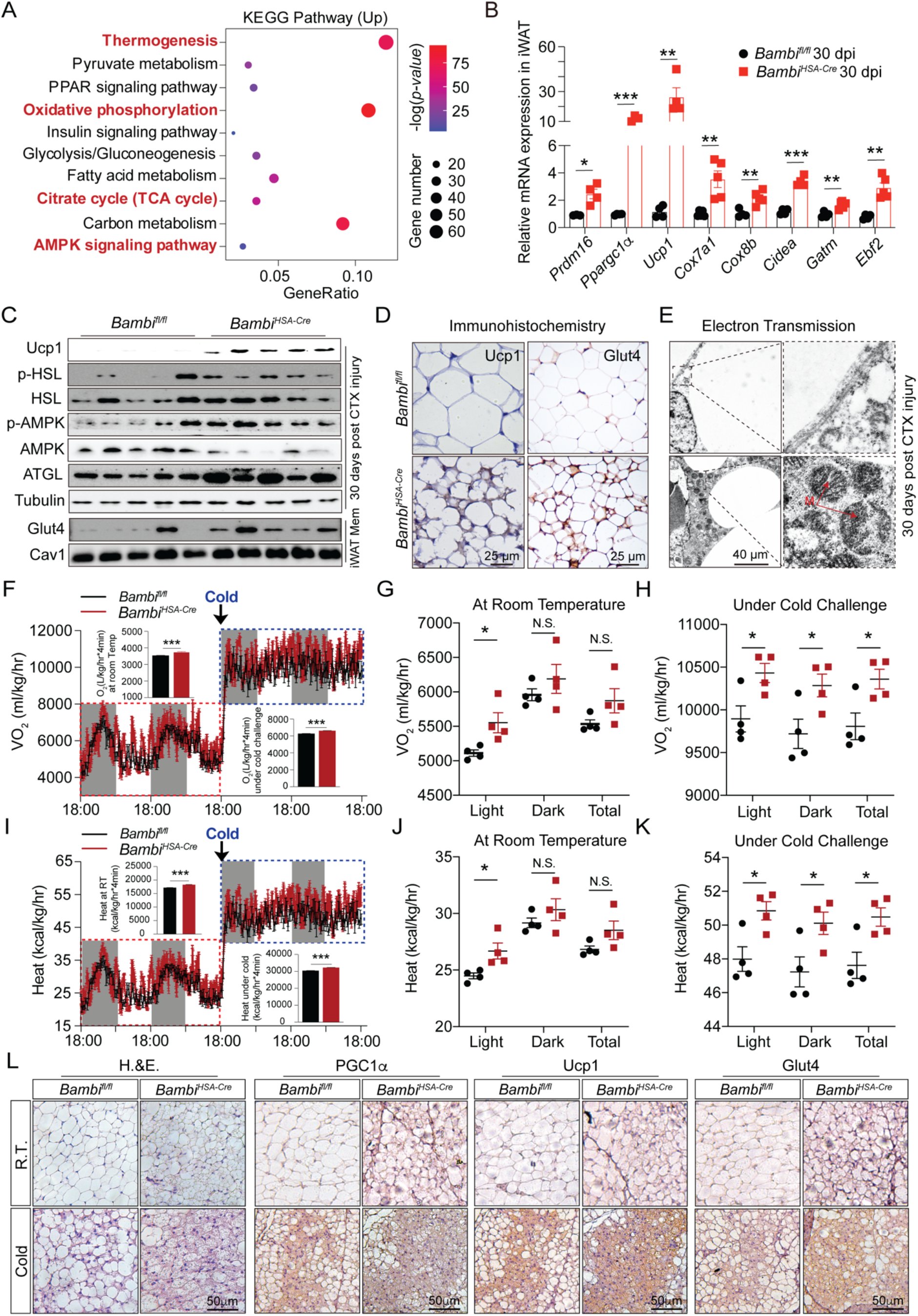
*Bambi* deletion in skeletal muscle activates the thermogenic capacity of iWAT. (A) KEGG pathway analysis of DEGs in iWAT of *Bambi^fl/fl^* and *Bambi^HSA-Cre^* mice 30 dpi of TA muscle. (B) qPCR analysis of thermogenesis- and beige adipocyte-related biomarkers in iWAT between *Bambi^fl/fl^* and *Bambi^HSA-Cre^* mice 30 dpi of TA muscle (n=5 mice/group). (C) Representative immunoblots of Ucp1, p-HSL, t-HSL, p-AMPK, t-AMPK, and ATGL protein levels in the iWAT of between *Bambi^fl/fl^* and *Bambi^HSA-Cre^*mice 30 dpi of TA muscle (n=5 mice/group, top). Glut4 and Cav1 protein levels in the membrane fraction of iWAT (n=5 mice/group, bottom) (D) Representative images of immunohistochemistry of Glut4 and Ucp1 in the iWAT of *Bambi^fl/fl^* and *Bambi^HSA-Cre^* mice 30 dpi of TA muscle. (E) Transmission electron microscopy images of iWAT sections from *Bambi^fl/fl^* and *Bambi^HSA-Cre^* mice 30 dpi of TA muscle. (F-H) Average oxygen consumption rate (VO_2_) was monitored over a 48-hr period for *Bambi^fl/fl^* and *Bambi^HSA-Cre^*mice at R.T. and under cold challenge (n=4 mice/group for two independent repeat). (I-K) Average heat generation was monitored over a 48-hr period for *Bambi^fl/fl^*and *Bambi^HSA-Cre^* mice at R.T. and under cold challenge (n=4 mice/group for two independent repeat). (L) Representative images of H.&E. and immunohistochemistry (PGC1α, Ucp1 and Glut4) of iWAT from *Bambi^fl/fl^* and *Bambi^HSA-Cre^* mice at R.T. and under cold challenge (4-6°C). \**p* < 0.05, \*\**p* < 0.01, and \*\*\**p* < 0.001, by two-sided Student’s t test. Data represent the mean ± standard error of the mean.

### Metabolic reprogramming of SVFs in subcutaneous adipose tissue results in the generation of beige adipocyte in mice with *Bambi* deletion in skeletal muscle

We speculated that in *Bambi^HSA-Cre^* mice, the SVFs were reprogrammed towards the precursors with high thermogenic potentials and differentiated into adipocytes with multilocular lipid droplets and high mitochondrial contents. To test this hypothesis, we explored an *ex vivo* culture model to evaluate the reprogramming of SVFs of iWAT in *Bambi^HSA-Cre^* mice (Figure S6A). The SVFs were isolated from both *Bambi^fl/fl^* and *Bambi^HSA-Cre^* mice and directly seeded into the Seahorse plate until fully confluency. Seahorse flux analysis showed that the oxygen consumption rate (OCR) was higher at all stages for SVFs from *Bambi^HSA-Cre^*mice than that of *Bambi^fl/fl^* mice, including the basal respiration, maximal respiration, proton leaky, ATP production and spare respiration capacity (Figure S6B-C). This suggested that the SVFs from *Bambi^HSA-Cre^* mice were reprogrammed to have higher metabolic potentials. Indeed, the epithelial mesenchymal transition (EMT)-related biomarkers, Ly6a, Cd34 and Kit were significantly upregulated in SVFs of *Bambi^HSA-Cre^* mice (Figure S6D). The metabolic reprogramming of SVFs was further confirmed by flow cytometry (increased number and intensity of CD34^+^ cells) and Mito-Tracker staining (Figure S6E-F). BODIPY and Mito-Tracker staining of differentiate primary adipocytes also proved that *Bambi^HSA-Cre^*mice-derived SVFs could differentiated into adipocyte with multilocular lipid droplet and a higher number of mitochondria (Figure S6G). Meanwhile, primary adipocytes from *Bambi^HSA-Cre^* mice also had a higher maximal respiration rate, higher thermogenic gene expression (*e.g.*, *Ucp1*, *Dio2*, *Cidea* and *Ebf2*) and corresponding thermogenesis-related proteins, such as Ucp1, PGC1α and Mito-complex (Figure S6H-K). Taken together, these data indicate that under physiological conditions, *Bambi*-deletion in skeletal muscle couples with the metabolic reprogramming of the SVFs, which contributes to the differentiation into multilocular adipocyte with higher thermogenic potentials.

### *Bambi*-deficient mice are resistant to high-fat diet (HFD)-induced metabolic disorders through enhanced systemic energy metabolism

As *Bambi* deletion in skeletal muscle enhances systemic energy metabolism, it is worthwhile to investigate whether *Bambi^HSA-Cre^*mice could resistant to high-fat diet (HFD)-induced metabolic disorder. We subjected *Bambi^HSA-Cre^* and *Bambi^fl/fl^* mice to a HFD to provoke metabolic disorders (Figure 4A). Consistent with the observation on a rodent chow, *Bambi^HSA-Cre^* mice were highly susceptible to HFD-induced body weight gain (Figure 4B-C). The net weight of tissue, such as TA, iWAT and liver, was greater in *Bambi^HSA-Cre^* mice but not for iBAT or eWAT (Figure 4D). However, contrary to body weight gain, symptoms of HFD-induced metabolic disorder, such as glucose tolerance and insulin sensitivity, were significantly mitigated (Figure 4E-H). Interestingly, after HFD feeding, even with a larger iWAT fat pad, the expression of thermogenesis and mitochondria-related genes like *Ucp1*, *Cidea* and *Cox7a1* remained elevated (Figure 4I). Notably, the activation of thermogenetic gene expression correlated with increased multilocular lipid droplets, as evidenced by *Ucp1* expression in iWAT of *Bambi^HSA-Cre^* mice (Figure 4J-K). The activation of iWAT browning was paired with enhanced expression of Glut4 and p-AMPK (Figure 4L). Concurrently, skeletal muscle also significantly contributed to the overall improvement of systemic energy metabolism via myogenic differentiation, muscular hypertrophy, and shift to slow-twitch fiber type (Figure 4M-N). Collectively, these results strongly underscore a protective role of *Bambi* deletion on systemic energy metabolism improvement by promoting adipogenesis *via* the hyperplasia but not the hypertrophy of subcutaneous adipose tissue, probably through the reprogrammed SVFs. Thus, we speculate that unknown factors or signaling pathways from skeletal muscle may contribute to the metabolic reprogramming of SVFs, leading to the beigeing of subcutaneous adipocyte and resistance to the HFD-induced metabolic syndromes.

**Figure 4.**
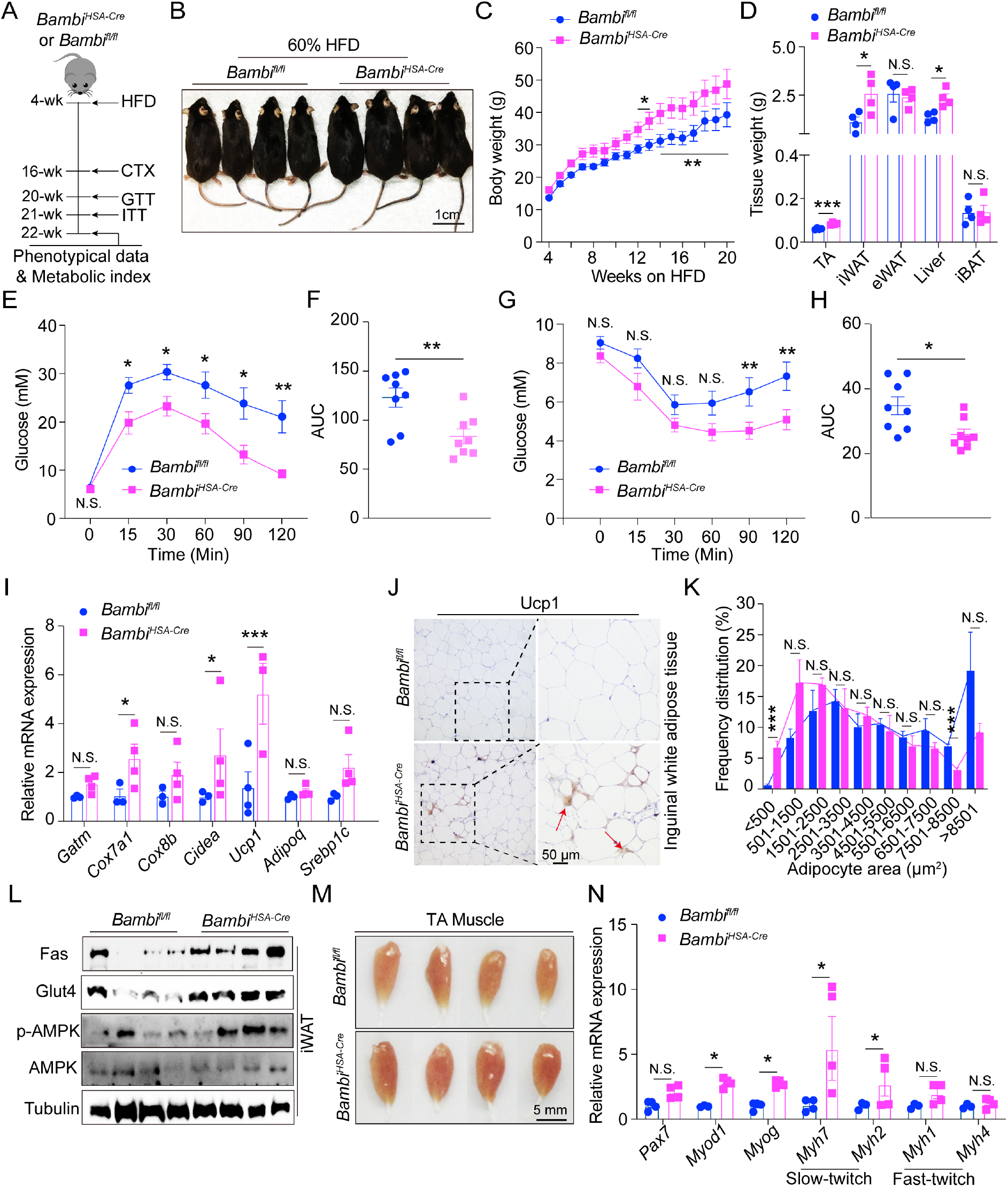
Mice with *Bambi* deletion in skeletal muscle resistant to HFD-induced metabolic disorders. (A) Diagram depicting the timeline of the HFD feeding plan for *Bambi^fl/fl^* and *Bambi^HSA-Cre^* mice (n=8 mice/group). (B) Representative images of *Bambi^fl/fl^* and *Bambi^HSA-Cre^* mice fed a HFD. (C) Body weight were measured over HFD feeding. (D) Tissue weights of TA, iWAT, eWAT, liver and iBAT. (E-F) GTT test of *Bambi^fl/fl^* and *Bambi^HSA-Cre^* mice. (G-H) ITT test of *Bambi^fl/fl^* and *Bambi^HSA-Cre^* mice. (I) qPCR analysis of the mRNA expression in iWAT. (J) IHC of Ucp1 in iWAT sections indicated that mice with *Bambi* deletion in skeletal muscle have a larger Ucp1^+^ area with multilocular lipid droplets. (K) Distribution of lipid droplet areas from iWAT. (L) Representative immunoblots of Fas, Glut4, p-AMPK, AMPK and Tubulin in the iWAT. (M) Representative images of TA muscle 30 dpi of regeneration in *Bambi^fl/fl^*and *Bambi^HSA-Cre^* mice. (N) qPCR analysis of *Pax7*, *MyoD1*, *Myog*, *Myh7*, *Myh2*, *Myh1*, and *Myh4* expression in TA muscle. N.S., not significant, \**p* < 0.05, \*\**p* < 0.01, and \*\*\**p* < 0.001, by two-sided Student’s t test. Data represent the mean ± standard error of the mean.

### Thbs4 expression and secretion is induced in *Bambi^HSA-Cre^* mice with muscle hypertrophy

To identify potential factors involved in the crosstalk between skeletal muscle and subcutaneous white adipocytes, we conducted serum proteomics for *Bambi^fl/fl^*and *Bambi^HSA-Cre^* mice. In the interest of screening secretory proteins expressed at lower levels, the hemoglobin proteins were first removed. We identified 95 differentially expressed protein biomarkers in serum, with 57 increased and 38 decreased proteins (Figure S7A-B). Gene ontology analysis (biological process) suggested that the increased secretory proteins in serum were associated with tyrosine phosphorylation, muscle cell migration and the insulin-like growth factor receptor signaling pathway (Figure S7C). To screen for potential secretory proteins released from skeletal muscle, we further analyzed the serum proteomics data, alongside the transcriptomic data of TA muscle from the matched mice against the secretory protein database^45^. Through this process, we identified two differentially regulated biomarkers in both TA muscle and serum at both mRNA and protein levels, Thbs4 and Jchain (Figure 5A-B). *Thbs4* mRNA expression level corresponded with the induced protein levels in serum (Figure 5B). Thbs4 is a member of the thrombospondins family, a group of stress-inducible secreted glycosylated proteins primary expressed in cardiac and skeletal muscle^46,47^. In *C57BL/6J* mice, *Thbs4* transcripts were presented at high levels in various skeletal muscles (*e.g.*, TA, EDL, Gas and Sol muscle) but is expressed minimally or not expressed in the other tissues like the liver and fats (Figure S7D). *Bambi* deletion led to an upregulation of *Thbs4* expression in skeletal muscle, which was then followed by an elevated Thbs4 protein level in serum and enrichment in iWAT of *Bambi^HSA-Cre^* mice (Figure 5C-D and Figure S7E). Secretion of Thbs4 was confirmed in *ex vivo* cultures of muscle tissue, suggesting that *Bambi*-deletion upregulates *Thbs4* expression and secretion from skeletal muscle (Figure S7F). The induced secretory Thbs4 protein level in the serum of *Bambi^HSA-Cre^* mice was accompanied by membrane enrichment on iWAT in the area of the multilocular lipid droplet (Figure 5E). In line with these evidence, we next investigated whether upregulated *Thbs4* expression, secretion and iWAT membrane enrichment were positively correlated with the beigeing of iWAT under cold challenge. *C57BL/6J* mice were subjected to a cold challenge sequentially for 1-, 3-, 5- and 7-days (Figure 5F). Compared to mice at R.T., plasma Thbs4 protein levels gradually decreased, while Thbs4 protein levels in iWAT slightly increased (Figure 5G). The cold challenge did not increase, but rather decreased, Thbs4 expression levels in the TA muscle (Figure 5H). Most importantly, along with cold-induced thermogenesis, it is worth noting that Thbs4 protein is highly enriched in beige adipocytes transdifferentiated from iWAT and eWAT, but not significantly in iBAT compared to R.T. condition, as demonstrated by Ucp1, PGC1α and Thbs4 IHC (Figure 5I and Figure S7G-H). Here, these findings pinpoint the newly identified skeletal muscle-derived myokine, Thbs4, being positively associated with the iWAT beigeing.

**Figure 5.**
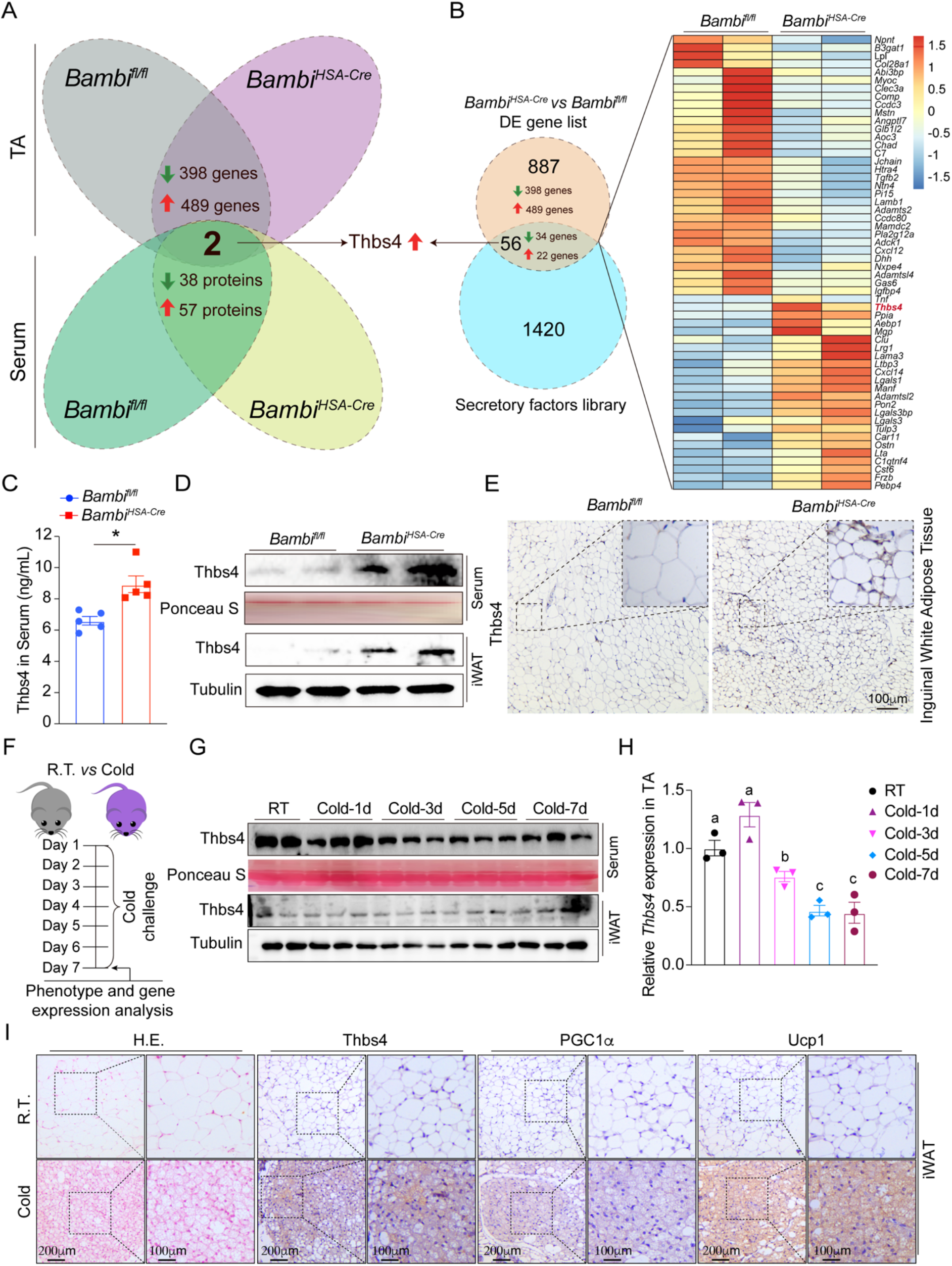
*Bambi* deletion-induced hypertrophy enhances Thbs4 expression and secretion. (A) Venn diagram indicating overlapping genes between the TA muscle transcriptome and matched serum proteomics between *Bambi^fl/fl^* and *Bambi^HSA-Cre^* mice. (B) Venn diagram and heatmap showing overlapping genes in TA muscle of *Bambi^fl/fl^* and *Bambi^HSA-Cre^* mice against the secretory protein library. (C) Thbs4 serum levels of *Bambi^fl/fl^*and *Bambi^HSA-Cre^* mice (n=5 mice/group). (D) Representative immunoblots of Thbs4 expression in iWAT and serum. (E) IHC staining of Thbs4 in the iWAT between *Bambi^fl/fl^* and *Bambi^HSA-Cre^*mice. (F) Diagram depicting the timeline of cold challenge for *C57BL/6J* mice. (G) Representative immunoblots showing the levels of Thbs4 in serum and iWAT from *C57BL/6J* mice with cold challenge. (H) qPCR analysis of Thbs4 expression levels in the TA muscle of *C57BL/6J* mice. Different letters indicate significant differences between two groups. (I) Representative images of H&E, Thbs4, PGC1α, and Ucp1 IHC in iWAT sections from *C57BL/6J* mice at the condition of R.T. or cold challenge for 7 days. \**p* < 0.05, by two-sided Student’s t test. Data represent the mean ± standard error of the mean.

### Thbs4 plays a significant role in metabolic enhancements triggered by aerobic exercise

Thbs4 expression and secretion are induced in skeletal muscle with hypertrophy in *Bambi^HSA-Cre^* mice. However, it remains unknown whether the enhanced Thbs4 expression and secretion can be recapitulated under physiological conditions. To test this, we measured *Thbs4* mRNA expression in skeletal muscle and protein in serum and adipocytes for mice subjected to aerobic exercise through treadmill running, compared to those under sedentary conditions. Thirty days post aerobic training resulted in attenuated weight gain and improved insulin sensitivity (ITT), but no significant difference in glucose tolerance test (GTT) (Figure S8A-D). Treadmill running also increased Thbs4 levels in serum, followed by membrane enrichment on beige adipocytes, which were also stained with Ucp1 and PGC1α (Figure S8E). The aerobic exercise-induced beigeing of iWAT was further confirmed by upregulation of genes associated with thermogenesis (*e.g.*, *Prdm16*, *Ucp1*, *Ppargc1α*, *Cidea*, *Cox7a1* and *Cox8b*), accompanied by elevated Ucp1 protein levels in iWAT and Glut4 protein enrichment on the membrane of iWAT (Figure S8F-H). Long-term aerobic exercise induced the expression of Thbs4 protein and its secretion into the serum (Figure S8G, I-J), also accompanied by an increase in the percentage of lean mass, but without a significant change in fat mass (Figure S8K). In line with the aerobic exercise data, we speculate that Thbs4 could be utilized as a skeletal muscle-specific secretory factor to mimic metabolic benefits from exercise or muscle hypertrophy.

### Activation of satellite cells and muscle hypertrophy contributes to upregulation of *Thbs4* expression

*Thbs4* expression patterns are dominantly and highly expressed in cardiac and skeletal muscle, whereas its expression is induced during injury^48^. Therefore, to further explore the regulatory mechanism of *Thbs4* upregulation in the skeletal muscle of *Bambi^HSA-Cre^* mice, we utilized an assay for transposase-accessible chromatin (ATAC) followed by high-throughput sequencing (HTS) to assess chromatin opening and identify potential transcriptional activators of the *Thbs4* promoter. Chromatin accessibility between *Bambi^HSA-Cre^* and *Bambi^fl/fl^*mice was dramatically altered with approximately 40% of peaks primarily located in the promoter region, others located in distal intergenic regions (∼23%) and the first intron (∼11%) (Figure S9A). To identify potential transactivator binding in the promoter region, we performed motif analysis and identified a group of highly enriched myogenic transcription factor binding sites in the proximal promoter region, including Mef2s, Myf5, MyoD, MyoG, and the E-box (Figure S9B). From ATAC-seq, 19304 peaks with opened chromatin accessibility were predicted to match 12357 genes. Integrative analysis against the DEGs between *Bambi^HSA-Cre^* and *Bambi^fl/fl^*mice resulted in 887 DEGs with 372 upregulated and 309 downregulated genes (Figure S9C-D). KEGG pathway analysis further demonstrated that these upregulated genes were related to skeletal muscle fiber development, regulation of muscle cell differentiation, and myoblast/myotube differentiation, suggesting both the increased muscle mass and activation of SCs contributed to the increased expression and secretion of Thbs4 (Figure S9E). Echoing the increased muscle mass, satellite cells activation and upregulation of myogenic regulatory factors (MRFs), such as MyoD and MyoG result in myogenic differentiation and skeletal muscle development ^14^. Upon aligning the peaks on chromatin, obvious chromatin openness was observed in the promoter regions of *Myod1* and *Myog* but not for *Pax7* (Figure S9F). In the proximal region of the *Thbs4* promoter region, we also identified a peak enrichment with a highly conserved MyoD/MyoG binding site, suggesting that MyoD and MyoG are directly involved in *Thbs4* upregulation (Figure 6F). To test this hypothesis, we measured *Thbs4* mRNA and protein expression in activated satellite cells (ASCs) in single myofibers and in C2C12 myoblasts during the proliferation and differentiation stages. Thbs4 protein was undetected in quiescent satellite cells (QSCs, Pax7^+^), while its level was highly induced in ASCs (Pax7^+^/MyoD^+^) after 3 days of *ex vivo* culture (Figure S9G). Moreover, during skeletal muscle regeneration and C2C12 differentiation, *Thbs4* mRNA and protein levels were positively correlated with MyoD and MyoG expression levels (Figure S9H-I). In C2C12 myoblasts with external overexpression of Thbs4 via adenovirus infection, the secretion of overexpressed Thbs4 was also identified in the medium (Figure S9J). Brefeldin A (BFA) indirectly inhibits protein migration from the endoplasmic reticulum to the Golgi apparatus ^49^. BFA inhibited Thbs4 protein secretion from C2C12 myoblasts in a concentration-dependent pattern in response to Ad-Thbs4-mediated overexpression (Figure S9J). These results suggest that during regeneration or hypertrophy, Thbs4 expression is induced in ASCs, myoblasts or myotubes and secreted through the ER-mediated secretory pathway.

**Figure 6.**
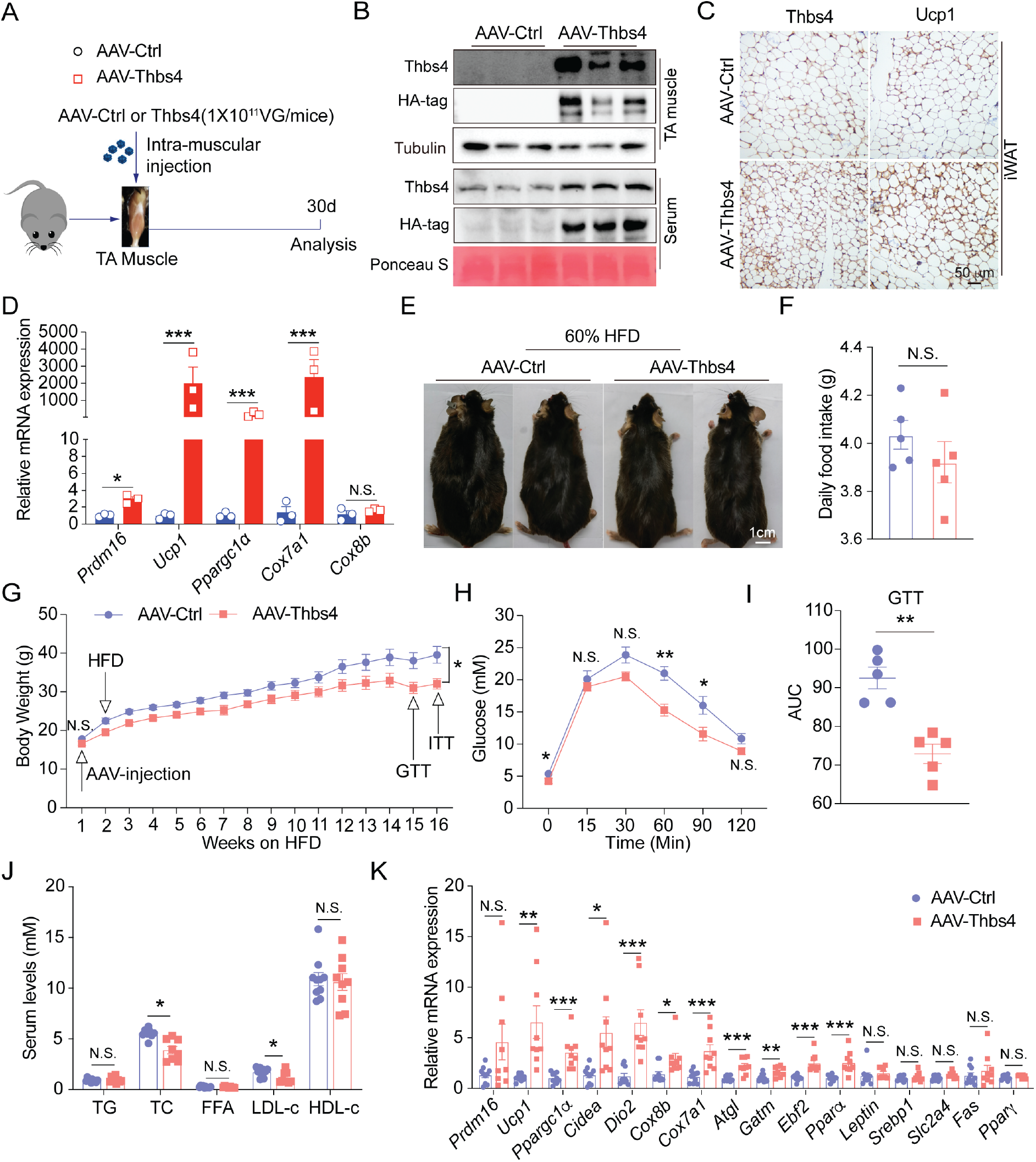
Thbs4 plays a long-term role in protecting against metabolic disorders. (A) Experimental scheme of intramuscular injection of AAV-Ctrl and AAV-Thbs4 groups (n=3 mice/group). (B) Representative immunoblots of Thbs4 protein in TA muscle and serum in the AAV-Ctrl and AAV-Thbs4 groups. (C) Representative images of IHC staining of Thbs4 and Ucp1 in the iWAT of mice with either AAV-Ctrl or AAV-Thbs4 intramuscular injection. (D) qPCR analysis of *Prdm16*, *Ucp1*, *Ppargc1α*, *Cox7a1*, *Cox8b* mRNA expression levels in iWAT in the AAV-Ctrl and AAV-Thbs4 groups. (E) Representative images of mice on HFD with intramuscular injection of either AAV-Ctrl or AAV-Thbs4 (n=5 mice/group). (F) Daily food intake for mice in the AAV-Ctrl and AAV-Thbs4 groups. (G) Body weight was measured weekly during the HFD feeding. (H-I) Glucose tolerance test (GTT) and AUC. (J) Serum levels of TG, TC, FFA, LDL-c and HDL-c between the AAV-Ctrl and AAV-Thbs4 groups. (K) qPCR analysis of thermogenesis- and beige fat-related gene expression in iWAT in the AAV-Ctrl and AAV-Thbs4 groups. N.S., not significant, \**p* < 0.05, \*\**p* < 0.01, and \*\*\**p* < 0.001, by two-sided Student’s t test. Data represent the mean ± standard error of the mean.

### External expression of Thbs4 promotes beigeing of white adipocytes

To explore the beneficial effect of Thbs4 on systemic metabolic homeostasis, we performed an intramuscular injection of AAV-Thbs4 to overexpress Thbs4 in skeletal muscle and thereby induce elevated serum levels of Thbs4 (Figure 6A). Thirty days post AAV injection, body weight, rectal temperature and TA muscle weight were not significantly different between the AAV-Ctrl and AAV-Thbs4 groups, except for a slight increase in the weight of iWAT (Figure S10A-C). An intramuscular injection of AAV-Thbs4 led to the overexpression of Thbs4 protein (detected by both HA and Thbs4 antibodies) in TA muscle and induction of externally expressed HA-tagged Thbs4 in serum (Figure 6B). AAV-Thbs4 also increased the membrane enrichment of Thbs4 in iWAT, along with the increased expression of Ucp1 in the region of multilocular lipid droplets and upregulation of thermogenesis-related genes (Figure 6C-D). To assess whether externally expressed Thbs4 localized to tissues other than iWAT, we performed immunohistochemistry for iBAT, heart, liver and pancreas against Thbs4, PGC1α and Ucp1. This indicated that externally expressed Thbs4 through skeletal muscle was not enriched in these tissues or possible virus leaky from skeletal muscle (Figure S10D). These data further demonstrate that external expression of Thbs4 activates beigeing of iWAT through the enrichment solely on the cellular membrane of iWAT.

### Thbs4 demonstrates a long-term protective role against high-fat diet-induced metabolic disorders by promoting beige fat activation in mice

To further evaluate the prolonged protective effect of Thbs4, *C57BL/6J* mice, having received intramuscular injection of AAV-Ctrl or AAV-Thbs4, were fed with a HFD for 16 weeks. High serum Thbs4 levels exhibited a markedly protective role against HFD, demonstrated by decreased body weight, fat mass (iWAT and eWAT) and liver weight (Figure S11A-C). These variables were significantly different, though there was no significant difference in food intake (Figure 6E-G). The AAV-Thbs4 injected group also showed improved glucose tolerance (GTT) and insulin sensitivity (ITT) (Figure 6H-I, Figure S11D-E). Serum lipid profiles indicated that higher serum levels of Thbs4 resulted in lower serum TC and LDL-c (Figure 6J). Both groups had similar serum TG, FFA, HDL-c, ALT and AST levels (Figure 6J, Figure S11F). Notably, the AAV-Thbs4 groups exhibited enhanced levels of thermogenic gene expression such as *Prdm16*, *Ppargc1a*, *Ucp1 Cidea*, *Dio2*, *Cox7a1*, *Cox8b*, *Gatm*, *Pparα* and *Ebf2*, as well as lipolysis-related genes, *e.g.*, *Atgl* (Figure 6K). H&E staining indicated that both groups displayed a similar phenotype in iBAT (Figure S11G), but the AAV-Thbs4 groups had long-term and systemic protective effects on inducing beigeing of iWAT and eWAT and, lipid clearance in the liver (Figure S11H-K). The phenomena of subcutaneous white adipocyte were in line with high serum levels of Thbs4 and increased Ucp1 protein levels (Figure S11L). Collectively, these results demonstrate a protective role for Thbs4 against HFD-induced metabolic disorders by promoting beige fat activation in mice, which highlights its potential therapeutic application in the treatment of metabolic disorders.

### *Thbs4*-KO exacerbates HFD-induced metabolic syndromes

In order to further evaluate Thbs4 as a critical myokine that participates in systemic metabolic regulation, we took advantage of the whole-body *Thbs4* knockout mice as a model to delve into the metabolic regulatory function of Thbs4. The metabolic cage measurement results indicated that *Thbs4*-KO mice showed lower VO_2_ and heat production than those of WT mice (Figure S12A-B). Thbs4 is predominantly expressed in skeletal muscle and we showed that its expression was completely ablated in TA and Sol of *Thbs4*-KO mice (Figure S12C). When fed with rodent chow, the thermogenic and lipid metabolism-related gene expressions were not significantly affected in both iBAT and iWAT, except *Prdm16*, *Cd36* and *Srebp1c* expression slightly reduced in iBAT (Figure S12D-E). For both WT and *Thbs4*-KO mice, the OCR of undifferentiated SVFs from iWAT did not significantly differ at any stages (Figure S12F). However, for the *ex vivo* differentiated white adipocytes, the OCR was significantly reduced, at the stage of the maximal respiration, and spare respiration capacity (Figure S12G-H). The *in vitro* cultured and differentiated SVFs from iWAT also showed larger lipid droplets, lower expression level of thermogenesis and lipolysis-related gene expression, at both mRNA and protein levels, such as Ucp1, PGC1α, Mito-complex and p-HSL (Figure 12I-K). Compared with the SVFs from aged mice, SVFs from *Thbs4*-KO mice presented similar expression pattern of Ucp1 of aged mice (Figure S12L). These data suggest that *Thbs4* ablation may abolished the metabolic reprogramming of SVFs in iWAT by Thbs4 towards state with low metabolic potential.

To further test this hypothesis, WT and *Thbs4*-KO mice were fed with 60% HFD for 25 weeks (Figure 7A). Compared to the WT mice, *Thbs4*-KO mice at the initial stage showed slightly better insulin sensitivity (ITT, Figure S13B), but no differences in glucose tolerance (GTT, Figure S13A). The *Thbs4*-KO mice had lower rectal temperature after 16-hr fasting (Figure S13C), suggesting that the *Thbs4*-KO may be closely associated with lower basal metabolism rate. After a 25-week HFD challenge, the body weight gain and food intake were lower for *Thbs4*-KO mice (Figure 7B-C), but the metabolic indices were further exacerbated for *Thbs4*-KO mice, such as TC, LDL-c, glucose tolerance and insulin sensitivity, except a slightly higher HLD-c (Figure 7D-F). After HFD feeding, the *Thbs4*-KO mice displayed a greater accumulation of lipids in the abdominal adipose tissue (eWAT) and a higher eWAT weight (Figure S13D-E). H.&E. staining revealed larger lipid droplets in iBAT, iWAT, eWAT, with more F4/80^+^ cells (inflammatory areas) observed in the section of iWAT and eWAT (Figure 7G, I and J). Furthermore, qPCR analysis demonstrated an increase in lipid absorption and *de novo* lipogenesis, especially the long chain fatty acid biosynthesis-related gene expression was significantly upregulated (Figure S13F), followed by the lipid accumulation in the liver of *Thbs4*-KO mice (Figure 7H). Our extended data showing that the *Thhs4* mRNA expression in skeletal muscles and protein levels in serum declined gradually with aging (Figure S14). These data demonstrate Thbs4’s role as a key myokine in systemic metabolic regulation, and also suggest that *Thbs4* ablation may alter the metabolic programming of SVFs, which could have implications for individuals with aging or metabolic syndromes.

**Figure 7.**
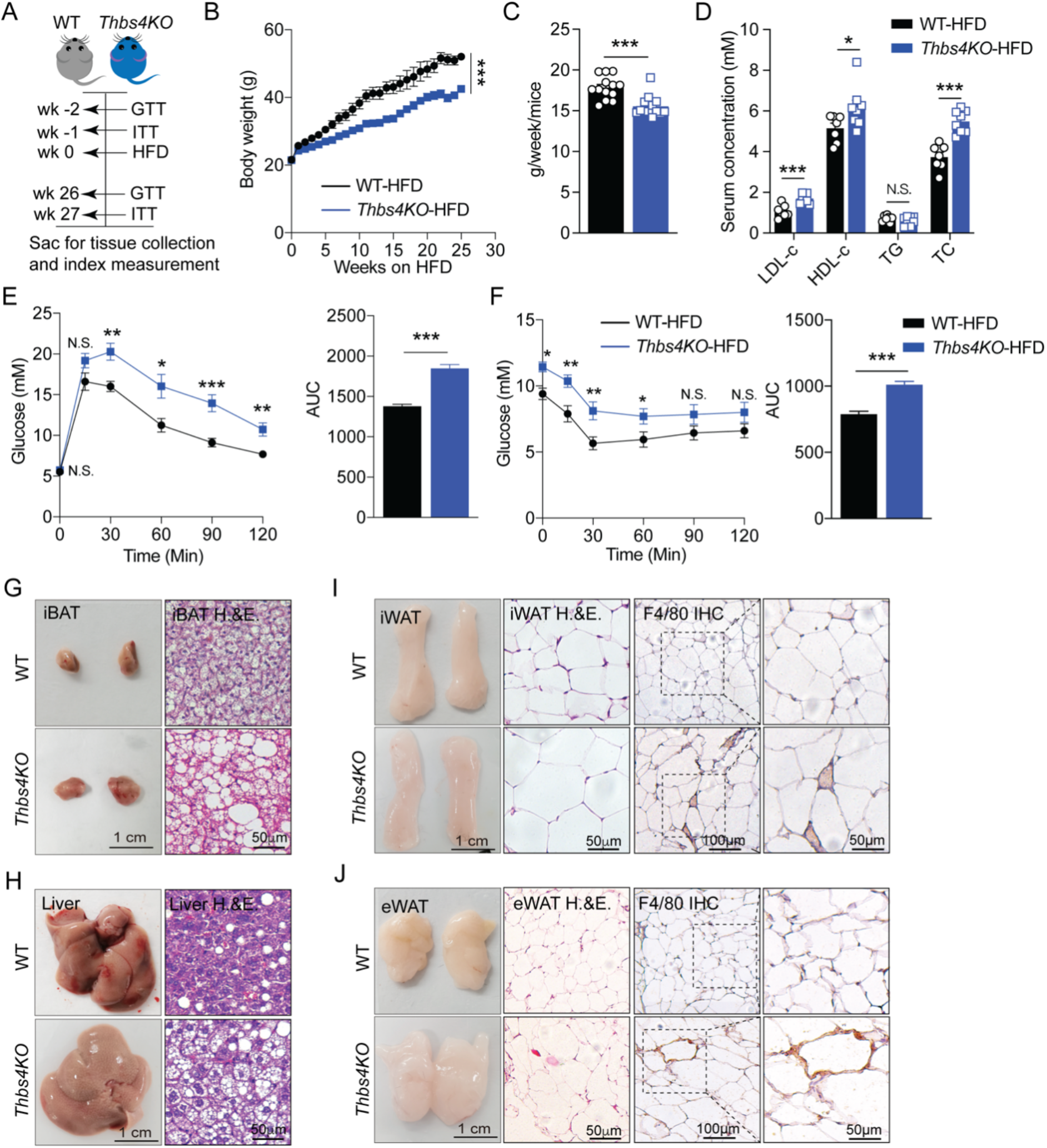
*Thbs4*-KO exacerbates HFD-induced metabolic syndromes. (A) Diagram depicting the timeline of the HFD feeding plan for WT and *Thbs4-*KO mice (n=8 mice/group). (B) Body weight was measured weekly during HFD feeding. (C) Daily food intake for mice on HFD feeding over 25 weeks. (D) Serum levels of TG, TC, FFA, LDL-c and HDL-c (n=8 mice/group). (E-F) Glucose tolerance test (GTT) and insulin tolerance test (ITT). (G) Representative images of iBAT and H.&E. staining. (H) Representative images of liver and H.&E. staining. (I) Representative images of iWAT, H.&E. and F4/80 IHC staining. (J) Representative images of eWAT, H.&E. and F4/80 IHC staining. N.S., not significant, \**p* < 0.05, \*\**p* < 0.01, and \*\*\**p* < 0.001, by two-sided Student’s t test. Data represent the mean ± standard error of the mean.

**Figure 8.**
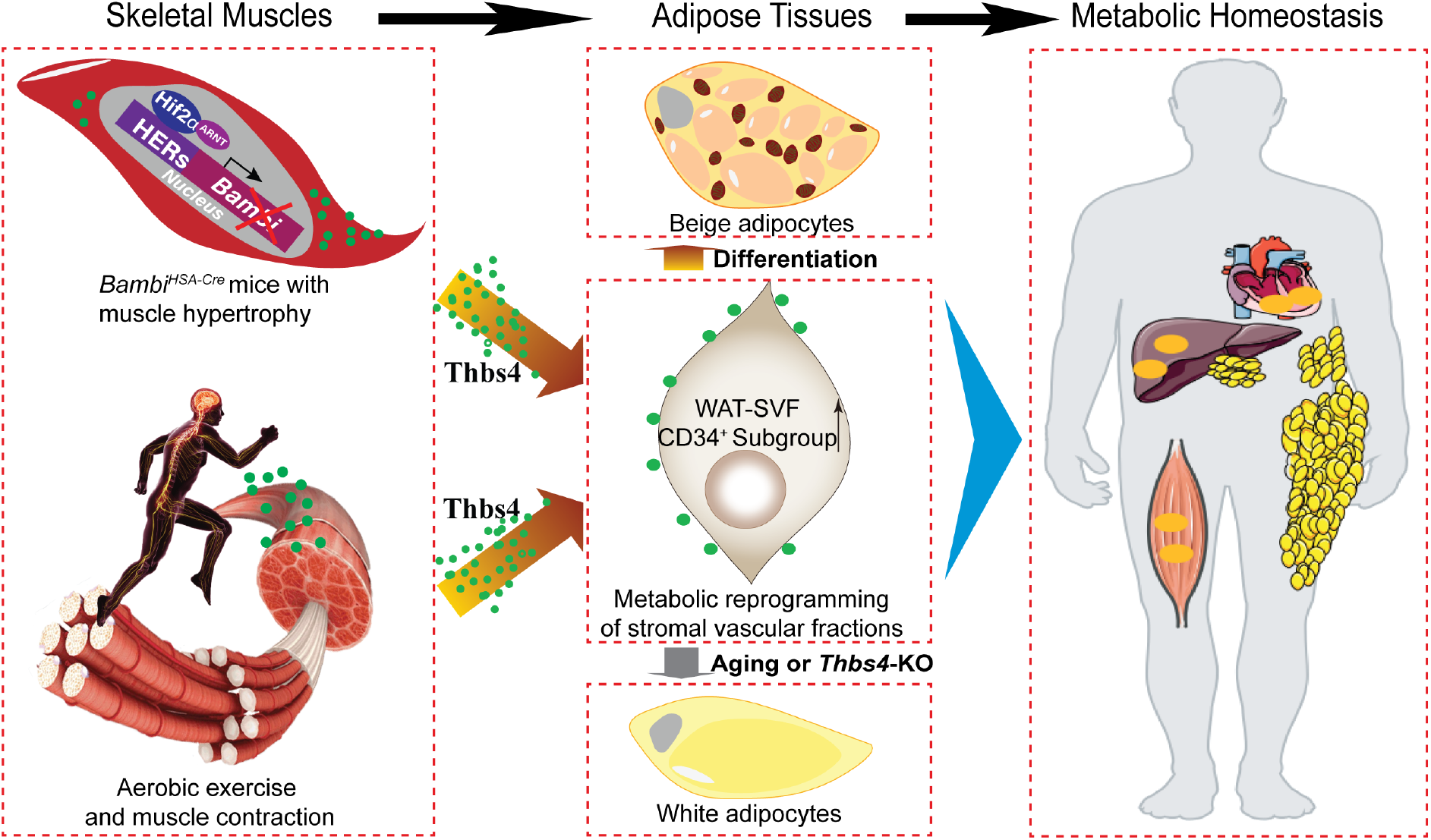
Working model of the skeletal muscle and adipose tissue crosstalk. This working model depicts how HIF2α-Bambi-Thbs4 axis in skeletal muscle drives muscle hypertrophy and improves systemic metabolic. Deletion of *Bambi* promotes the metabolic reprogramming of the stromal vascular fraction of white adipose tissue, enhancing beige adipocyte development. The myokine Thbs4, upregulated in subjective of muscle hypertrophy or aerobic exercise, is identified as a critical factor in this transformation, supporting the browning of white adipocytes and offering long-term metabolic protection against metabolic disorders. This model underscores the potential of Thbs4 as a therapeutic target in the treatment of metabolic disorders.

## Discussion

Exercise has a broad spectrum of benefits on systemic metabolic homeostasis, including weight loss, resistance to metabolic syndromes and prevention of cardiovascular diseases^6,25,50^. However, how physical training improves systemic metabolic homeostasis and through which approaches or signaling pathways the internal organs communicate with each other regarding the metabolic benefits are less investigated. In particular, understanding how to manipulate or recapitulate the beneficial effect of exercise for inactive or aged individuals with accompanying metabolic syndromes requires in-depth exploration. In this study, leveraging multi-omics, bioinformatics analysis and different experimental models, we identified a muscle hypertrophy or exercise-induced myokine, Thbs4 that is involved in the beigeing of subcutaneous adipose tissue and systemic metabolic improvement. Moreover, external overexpression of Thbs4 is protective against HFD-mediated metabolic disorders, while *Thbs4*-KO exacerbates HFD-induced metabolic syndromes, shedding light on its potential applications to treat metabolic syndromes.

One of the significant findings is that *Bambi*-deletion induces muscle hypertrophy. As a HIF2α downstream target gene, *Bambi* deletion is consistent with our previous study showing that transient pharmaceutical inhibition of HIF2α accelerates muscle regeneration and increases myofiber diameters^18^. However, long-term ablation of *Hif2α* both in SCs or skeletal muscle results in disruption of SC homeostasis, regeneration defects and skeletal muscle atrophy^18,51^. *Hif2α* ablation *via* the *Myog*- or *Ebf2-Cre* recombinase system results in an increase in fast-twitch fibers, with a phenotype of an increased percentage of Type IIB fibers and a reduced percentage of Type I fibers^39^. Although the number of SCs was not precisely quantified, skeletal muscle with switching from Type I to IIB fiber type will be more glycolytic, have less endurance and display decreased resistance to muscular atrophy. Skeletal muscle atrophy during aging and denervation is due to the reduced number of SCs, as well as defects in SC activation, proliferation and differentiation^52^. *Hif1/2α* double deletion is driven by the *MyoD-Cre* system in embryonic myoblasts, resulting in normal muscle development and growth^19^. This discrepancy suggests that HIF2α may exert distinct and stage-specific roles during embryogenesis, postnatal development and injury/regeneration^53^. HIF2α also targets a spectrum of genes involved in the regulation of cellular inflammation, cell cycle arrest and energy metabolism^54^. Therefore, understanding HIF2α-mediated transient and long-term mechanisms is practically important. Our ChIP-seq data indicate that HIF2α directly targets multiple biological functions, such as the TGF-β/Wnt signaling pathway, tissue development and nutrient metabolism, suggesting the necessity of identifying downstream targets to induce beneficial phenotypes. Further analysis demonstrated that HIF2α regulates *Bambi* expression *via* direct binding to HREs in the *Bambi* promoter, in line with the data of HIF2α overexpression on *Bambi* expression in skeletal muscle and myoblasts. *Bambi* deletion recapitulates the phenotype of muscle hypertrophy in HIF2α small molecular inhibitor injected TA muscle, suggesting that *Bambi* may be an alternative HIF2α downstream target for constant fiber enlargement and muscle growth. In agreement with our findings, *Bambi* is also reported to be a novel HIF1α-regulated gene that contributes to hypoxia-induced cell polarity in malignant carcinomas^55^. Under physiological and aerobic training, skeletal muscle is subjected to low oxygen conditions^56^, suggesting that *Bambi* deletion may avoid additional influence on other HIF2α-regulated genes or pathways required for cell proliferation, tissue development and metabolism, also with bona fide observation on oxidative fiber switching other than muscle hypertrophy.

Fiber type switching is a significant medical concern, as slow-twitch muscle with type I or IIA fiber (oxidative fibers) is more resistant to muscle loss and atrophy due to aging. Physical training typically promotes the switch from type IIb/X (more glycolytic) to type I/IIA (more oxidative). This is also practically important for individuals against metabolic syndromes. HIF2α deletion reported in two different studies either did not alter fiber type in TA muscle^18^ or resulted in switching from Type I to IIB^39^. However, another major finding of the current study is that *Bambi* deletion activates mitochondrial biogenesis with a preference for fiber type switching to oxidative phosphorylation. These suggest that *Bambi* may mediate an alteration in the skeletal muscle microenvironment that may cause alternative manipulation of the cellular response to energy metabolism. Previously, we demonstrated that Bambi protein is dominantly detected in fast twitch fibers of IIB^+^ and IIX^+^ in TA, EDL and Gas muscles but not in the soleus muscle^57^. Therefore, it is reasonable to speculate that downregulation of the *Bambi* gene may be associated with an alternative switch toward oxidative metabolism. Here, we showed that *Bambi* deletion in TA muscle increases Type I fibers, activates mitochondrial biogenesis and boosts the ATP metabolism. Beyond local metabolic alterations in response to fiber type switching in skeletal muscle, systemic benefits, especially the improvement of metabolic syndromes, are linked with metabolic reprogramming in skeletal muscles. Several recent studies also report the link between alteration on skeletal muscle and systemic metabolic improvement. For instance, clenbuterol administration causes significant systemic improvement in glucose metabolism *via* selective β2-adrenergic receptor activation in skeletal muscle^58^. Systemic or muscle-specific manipulation of ubiquitin-specific proteases (USPs), USP21, leads to changes in myofiber conversion to type I, muscle mass and energy expenditure, benefiting whole-body energy metabolism^59^. In addition to fiber type conversion, *Bambi* deletion in skeletal muscle also couples with increased overall mitochondrial contents observed by transmission electron microscopy. Mitochondria are critical organelles for the regulation of glucose, lipid and systemic energy metabolism^60^. This idea was supported by the outcome of indirect calorimetry cages in which energy expenditure and oxygen consumption were higher in *Bambi^HSA-Cre^* mice either post injury or cold challenge. Consequently, increased mitochondrial activity and systemic energy metabolism coordinately contribute to the metabolic response of skeletal muscle, leading to improved glucose tolerance and insulin resistance. Interestingly, under the cold challenge, the non-shivering thermogenesis derived from the beigeing of white adipose tissue further contributes to the improved basal metabolism on energy expenditure. This could be further supported by the DEGs, including thermogenesis, oxidative phosphorylation, TCA cycle, and the AMPK signaling pathway in adipose tissues of *Bambi^HSA-Cre^*mice. The activation of beigeing and thermogenesis is well established and recognized as a promising approach for combating metabolic syndromes, such as obesity, type 2 diabetes and fatty liver disease^61,62^. However, how the skeletal muscle communicates with adipose tissue with respect to the activation of beigeing and the induction of lipolysis remains unknown.

To uncover the potential mechanism by which the thermogenic pathway is activated in iWAT, we performed serum proteomics to identify novel muscle contraction- and hypertrophy-related myokines. An integrative analysis of proteomic data and RNA-seq of TA muscle for matched mice against the secretory protein database was performed. Among the co-regulated and muscle-derived secretory proteins, Thbs4 was the only one upregulated at both the mRNA and protein levels. Thbs4 is a member of the thrombospondins (Thbs) family, a group of secreted matricellular proteins, involved in diverse biological processes such as cell proliferation, adhesion and migration^46,63^. The Thbs family consists of five genes in mammals, with Thbs4 predominantly expressed in cardiac and skeletal muscles^64^. However, our data demonstrated that in healthy adult *C57BL/6J* mice, Thbs4 is highly expressed in skeletal system compared to other tissues like liver, adipose and lung. Within the cells, Thbs4 has a critical cardioprotective^65^ and myofiber protective function during muscle injury, dystrophy or wasting^66^. On the other hand, recent studies also highlighted that Thbs4 expression is markedly increased under pathological conditions such as heart failure and the development of atherosclerotic lesions, typically seen in heart remodeling and ApoE(-/-) mice^67,68^. The deletion of Thbs4 in these models has been shown to alleviate local inflammation in both the atherosclerotic lesions and failing hearts, primarily through the downregulation of endothelial adhesion proteins and chemokines^67,68^. Consequently, in the cardiovascular system, Thbs4’s role appears paradoxical, as its expression can exert both protective and detrimental effects depending on the context. Future research is required to elucidate this complexity and identify the specific regulatory mechanisms governing the differential effects of Thbs4. In muscular system, overexpression of Thbs4 in myofibers has been found to rescue muscle dystrophy in both mouse and Drosophila models^66^. Its protective effect is mediated through activating and interacting with transcription factor 6 (ATF6α), enhancing the adaptive aspect of ER stress response^69–71^. Interestingly, Thbs4, traditionally characterized as a secreted extracellular matrix (ECM) and matricellular protein, is continually released outside the cells depending on intracellular levels of Ca^2+^ during physical exercise, muscle contraction and injury/regeneration^64^. A study investigating muscle intrinsic and secretory regulators identified a group of exercise- and contraction-responsive secretome during concentric or eccentric exercise^72^. Among them, Thbs4 is recognized as an autocrine or paracrine modulator over muscle cells^72^. Moreover, after extended cycling training, circulating Thbs4 protein acts as an endocrine regulator to improve cardiorespiratory fitness (VO_2_)^73^. Furthermore, high-intensity aerobic exercise (with a higher maximum oxygen consumption rate) for six-week significantly boosts *Thbs4* expression level (∼10-fold) in muscle biopsies from volunteers^73^. A bout of exercise is positively associated with increased serum Thbs4 protein levels in multiple studies^72,74^. In agreement with these studies, aerobic exercise induces *Thbs4* mRNA and protein expression in skeletal muscle, followed with up to a 20% increment of Thbs4 protein in serum, which is a key biological step for Thbs4-mediated metabolic benefit. Therefore, it seems likely that muscle-derived Thbs4 could not only protect myofibers from injury but also extend its benefit globally to other organs. However, these studies did not consider the age of participants or animal models. As our and other studies have shown, Thbs4 levels in skeletal muscles and serum decline gradually with aging. Therefore, whether Thbs4 is physiologically important from adulthood to middle-aged and aged periods remain unknown.

A recent study using a *Thbs4*-KO model examined the connection between Thbs4 and muscle dystrophy (MD)-related membrane stability, cellular attachment and skeletal muscle integrity^66^. In MD models (*e.g.*, LGMD2F with δ-sarcoglycan deletion, or mdx background) with Thbs4 deletion, the MD phenotype is robust and predisposed. However, in wild type mice, MD phenotype is marginal until they reach advanced age. Our data show that Thbs4 protein in serum and skeletal muscle start to decrease at 50 weeks of age. This observation aligns with previous findings that in *Thbs4*-KO mice, the dystrophic pathology does not worsen until 6-month or even 1-year old^66^, suggesting Thbs4’s age-dependent metabolic regulatory mechanism may be partially compensated by other factors. Overexpression of Thbs4 can protect *mdx* mice from the MD throughout their lifetime ^66^. Interestingly, beyond cardiac and skeletal muscle, Thbs4 protein levels are higher in the serum of young mice and are inversely associated with brain function and synapse numbers^75^. Mass spectrometry revealed that in the serum of young mice (but not aged mice), Thbs4 and SPARC-like protein 1 (SPARCL1) are enriched, and the recombinant proteins increase dendritic arborization, and double synapse number in an *in vitro* system^75^. This study suggests a positive correlation between serum levels of Thbs4 and age. Consistent with this, our data indicate *Thbs4* mRNA expression in skeletal muscle and protein level in serum kept decreasing in rodents (Figure S14). However, it remains challenging to confirm whether Thbs4 physiologically acts as a rejuvenation factor that crosses the blood-brain barrier or functions in other organs to combat the age-driven progressive decline in multiple functions.

Aging is indeed closely associated with metabolic disorders, such as hyperglycemia and hyperlipidemia, primarily due to the impairment of insulin sensitivity and mitochondrial function in liver, skeletal muscle and adipose tissues^76^. It is not yet fully understood whether muscle-derived myokines sever as critical regulatory factors to improve whole-body glucose tolerance and insulin sensitivity. Interestingly, similar to the observation for IL-6 and irisin, Thbs4 was found to be highly enriched on the membrane of beige adipocytes under cold challenge. External overexpression of Thbs4 through intramuscular injection of AAV-Thbs4 led to elevated serum levels of Thbs4 under both physiological and pathophysiological conditions. This is associated with the activation of mitochondrial biogenesis, increasing respiratory capacity, and improving glucose tolerance, even under HFD conditions. This was accomplished by increasing the phosphorylation of AMPK and HSL as well as dynamically regulating mitochondrial function. Mitochondrial content in adipose tissue and the activation of lipolysis are critical steps in metabolic homeostasis for individuals with metabolic syndrome^60^. Despite the known involvement of secreted Thbs4 in angiogenesis, heart, nervous system, myocardium and vasculature remodeling, our study is the first to report a link between Thbs4 and long-term protection and improvement of metabolic homeostasis. This long-term protection could be attributed to Thbs4’s regulatory effect on adipogenesis from SVFs and further extend to the adipose expansion.

White adipose tissue (WAT) expansion and remodel have direct implications for metabolic syndromes risk, such as obesity and Type 2 Diabetes (T2D). It is been shown that the preferential expansion of subcutaneous rather than visceral WAT is key to resistance against glucose tolerance and insulin resistance. In current study, another interesting finding was that *Bambi^HSA-Cre^* mice displayed heavier body weight with an expansion of subcutaneous adipose tissue mass, in addition to moderate increase of lean mass. The biological process of adipose expansion is mediated via the adipocyte differentiation, also known as adipogenesis. This can occur by enlarging existing adipocytes (adipocyte hypertrophy), which is associated with maladaptive and pathological remodeling or by forming new adipocyte (adipocyte hyperplasia), which is an adaptive and beneficial state for metabolic health^77,78^. The biological process of adipogenesis is mediated by a subpopulation of SVFs, or adipocyte precursors committed to an adipogenic lineage (or called preadipocytes)^77,78^. Recent studies have indicated that lipid redistribution and a higher adipogenesis rate result in adipose expansion and systemic metabolic healthy during caloric excess in the setting of HFD but positive energy balance. Some transgenic obese mouse models (*e.g.*, *Adiponectin*, *mitoNEET*, and *Tnmd*) demonstrate better glucose tolerance, insulin sensitivity, lipid profile, and less inflammation^79–81^. These mice have large adipose tissue but smaller, more multilocular adipocytes with increased mitochondrial content, similar to the observation made in our *Bambi^HSA-Cre^*mice. The regulatory mechanisms leading to hyperplasia expansion with small but more mature adipocyte remain unknown. Our study provides data supporting the involvement of Thbs4 in the metabolic reprogramming of SVFs and adipose expansion. As shown in *Bambi^HSA-Cre^*and *Thbs4*-KO mice, differentiated SVFs display completely opposite directions of energy metabolism, as reflected by the oxygen consumption rate (OCR), thermogenesis-related gene expression and the size of adipocyte in *in vitro* culture system. Furthermore, *Thbs4*-KO mice display no difference at GTT and even slightly better insulin sensitivity, but are predisposed to metabolic disorders upon HFD challenge. These metabolic phenotypes are completely opposite to that of *Bambi^HSA-Cre^* mice and the serum levels of Thbs4 is a determinant regulator. These *in vivo* and *in vitro* phenotypes are similar to what is observed in aged mice with metabolic disorders and less sensitivity to adipocyte beigeing^82^.

Alongside investigation into the function of Thbs4 cross various tissues, recent studies have demonstrated that the upregulation of Thbs4 expression is mediated by the TGF-β1-SMAD3 signaling pathway in angiogenesis^83^. TGF-β is a state-specific and process-dependent cytokine, that is associated with many cell types and involved in tissue remodeling. Bambi is reported to negatively regulate TGF-β family signaling via its interaction with TGF-β receptors and inhibiting the formation of functional receptor complexes without interfering with ligand binding^84^. As a result, silencing or decreasing *Bambi* expression releases the TGF-β receptor to initiate signaling. Thus, deletion of *Bambi* to activate TGF-β signaling could be one potential regulatory mechanism to upregulate Thbs4 expression in *Bambi^HSA-Cre^*mice. Hence, to precisely understand the regulatory mechanisms, we performed ATAC-seq, along with the integrative analysis of RNA-seq from the matched model. Chromatin openness in the promoter region of Thbs4 has conserved MyoD and MyoG binding sites, followed by observation of SC activation and upregulation of MRFs in *Bambi^HSA-Cre^* mice. Our data demonstrated that expression of MyoD, MyoG and Thbs4 is upregulated in the TA muscle of both *Bambi^HSA-Cre^*mice, in SCs of *ex vivo* cultured myofibers and in C2C12 myoblasts during proliferation and differentiation. These data suggest that SC activation in response to external stimulation, such as injury/regeneration, physical training and genetic manipulation, is closely associated with upregulation and secretion of Thbs4.

Overall, our study characterizes a novel HIF2α downstream target gene, *Bambi*, as an alternative target for transient inhibition of HIF2α in skeletal muscle for the phenotype of muscle hypertrophy. *Bambi*-deletion not only induces muscle hypertrophy and oxidative switching of fiber type but also systemically extends metabolic benefits by activating adipocyte beigeing. This is mediated by a skeletal muscle-derived myokine, Thbs4, which exerts long-term protective effects on HFD-induced metabolic disorders and sheds light on potential applications for the development of novel strategies for the treatment of metabolic syndromes.

### Limitations of the study

In this study, we have found that the upregulation of Thbs4 expression in skeletal muscle and secretion into serum can be coupled with its enrichment on the membrane of iWAT and the beigeing of iWAT via activation of the p-AMPK and p-HSL signaling pathways. However, the receptor for Thbs4 on iWAT, particular beige adipocytes, remains unidentified. Although previous studies have reported that Thbs4 may interact with multiple integrins in skeletal muscle, vascular, endothelial and smooth muscle cells^48^, we need further research to confirm the identity of Thbs4’s receptor in iWAT. This will validate the interaction and possibly help us understand the downstream signaling required to activate adipocyte beigeing. Furthermore, Thbs4 level are found to decline gradually with aging. Whether an increase in Thbs4 protein in serum could improves white adipocyte beigeing and ameliorates aging-mediated metabolic disorders is a question that needs to be addressed. Moreover, exploring the correlation between Thbs4 serum levels and various body/metabolic indices, such as age, Body Mass Index (BMI), or the impact of aerobic training, in human populations could yield significant insights. Furthermore, assessing the therapeutic potential of Thbs4 in treating metabolic disorders warrants additional exploration to understand the specific pathological conditions that might benefit from Thbs4 application. Moreover, polymorphisms of Thbs4 have been associated with a recurrent risk of cancers^85^ and vascular inflammation^86^. It would be worthwhile to investigate whether there is a relationship between Thbs4 polymorphisms and the risk of metabolic syndromes. Such studies could pave the way for new therapeutic targets and potential strategies to prevent or treat metabolic disorders and possibly extend healthy aging.

## STAR★METHODS

### KEY RESOURCES TABLE

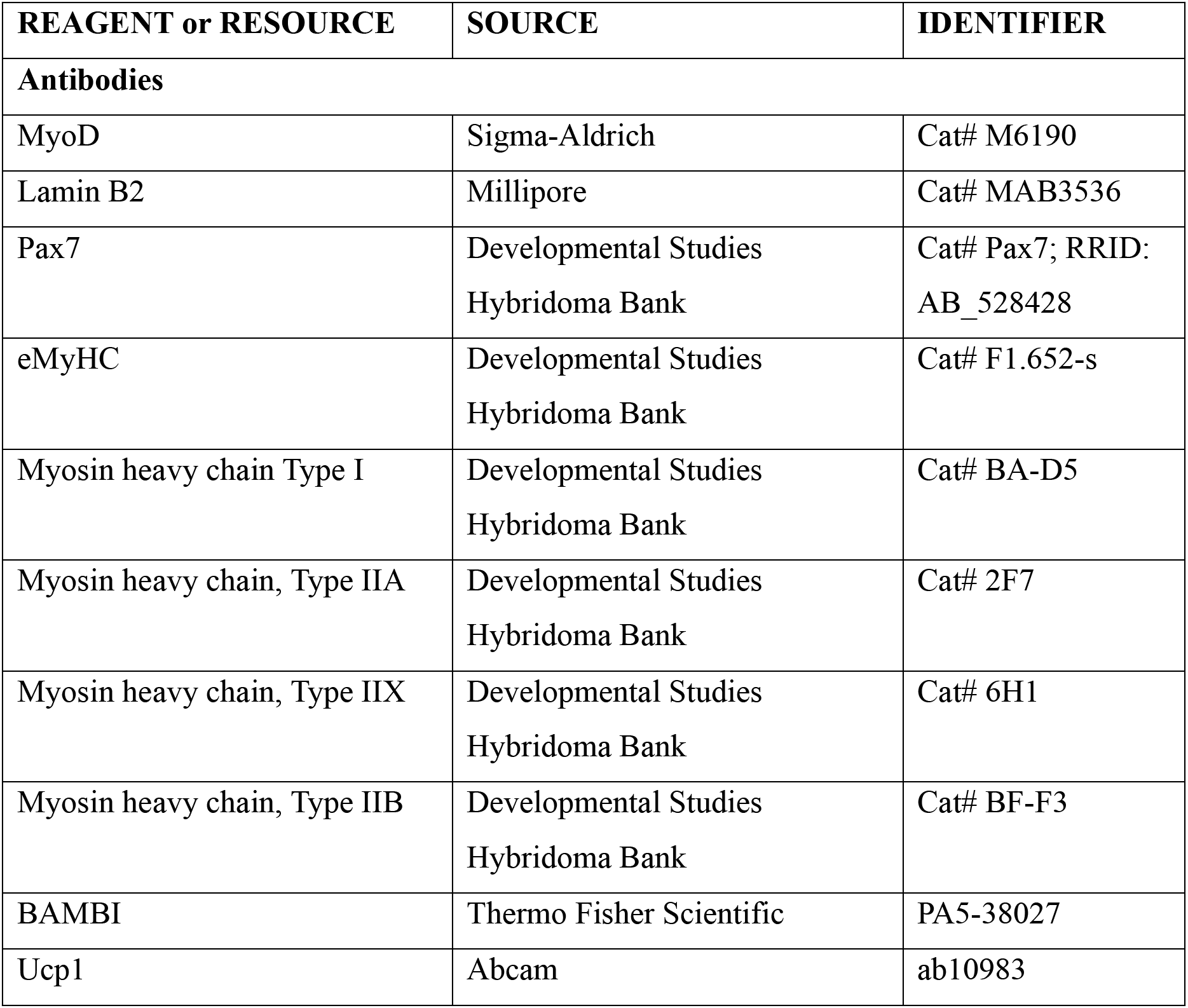
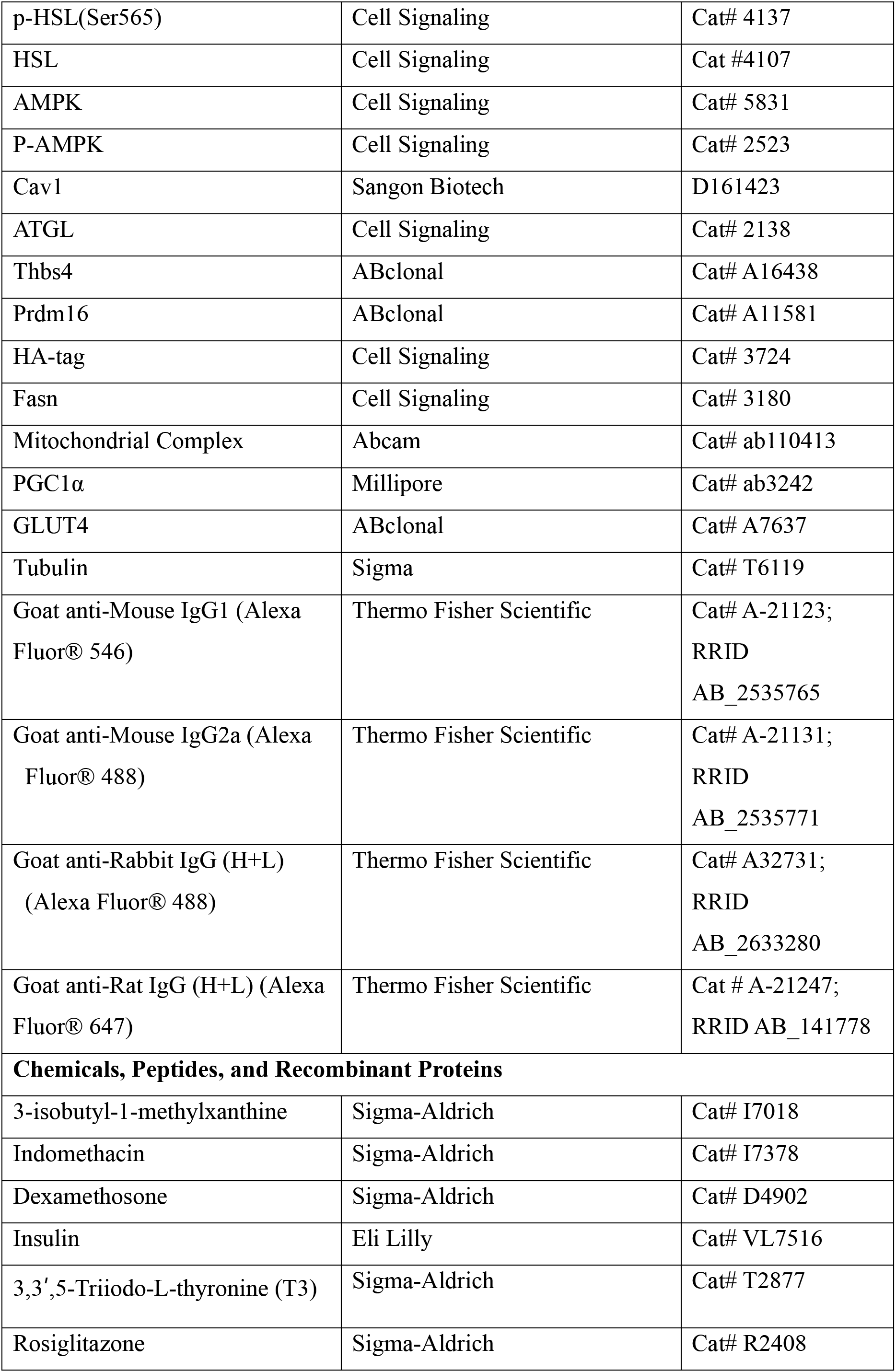
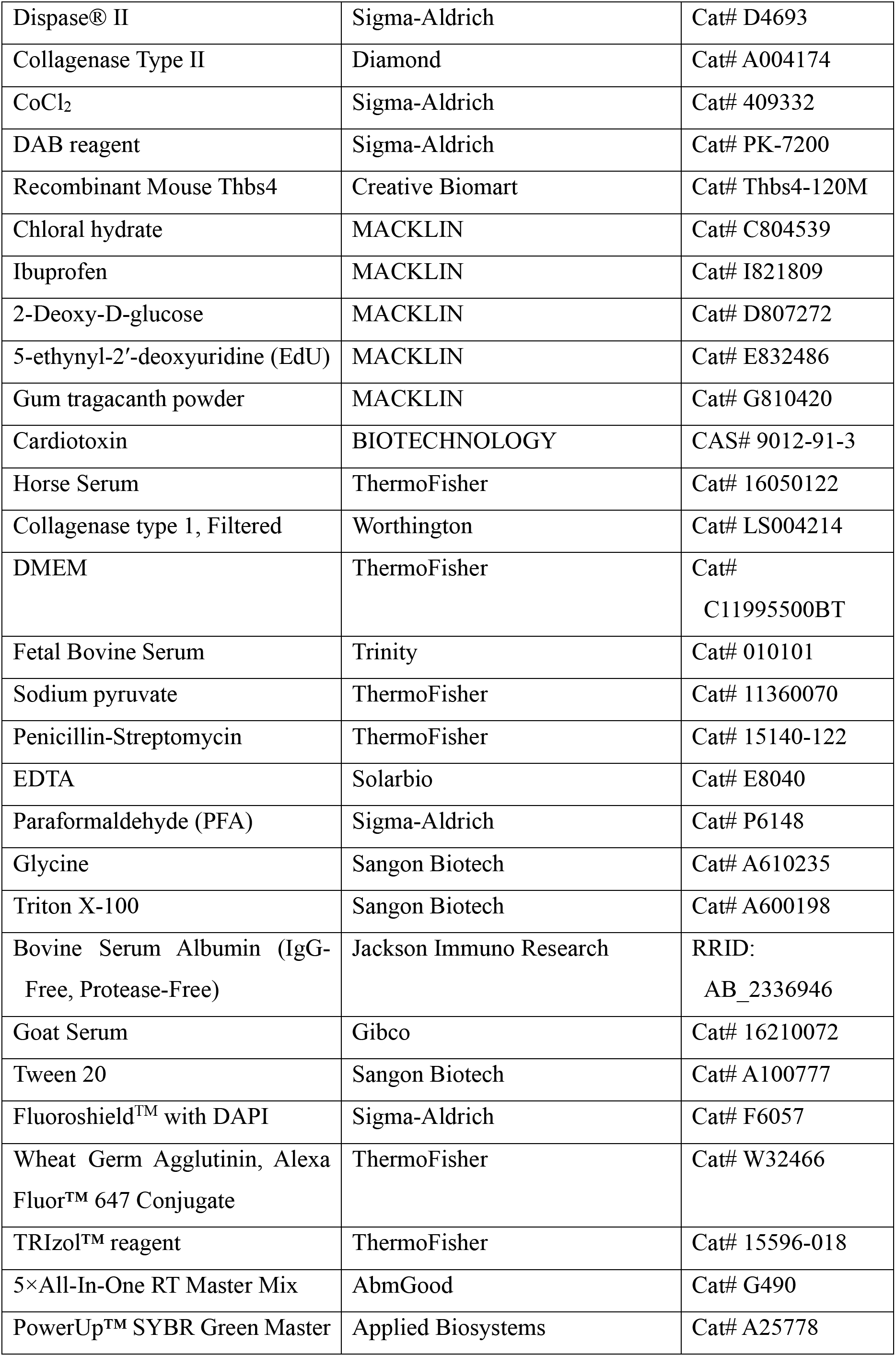
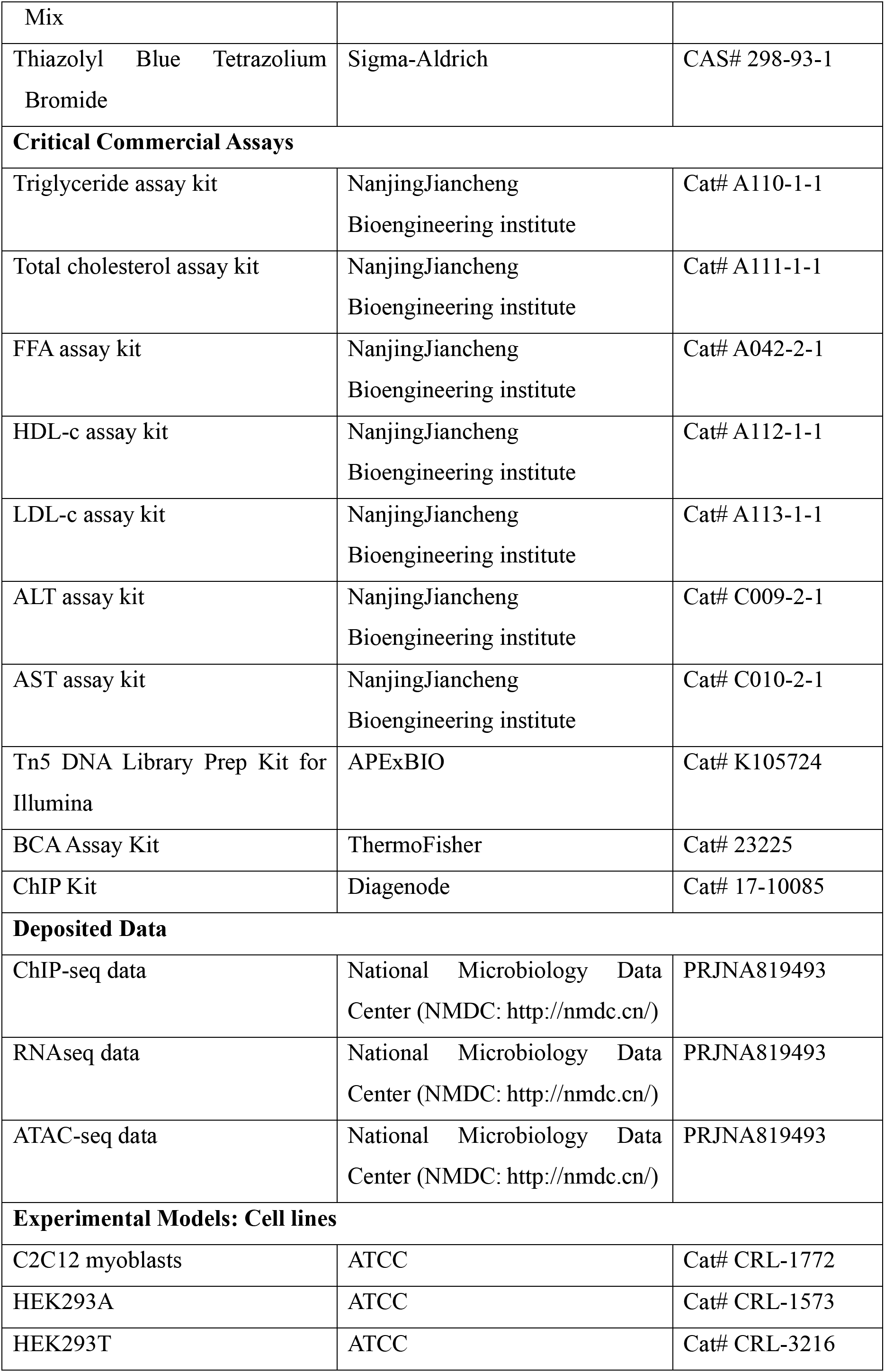
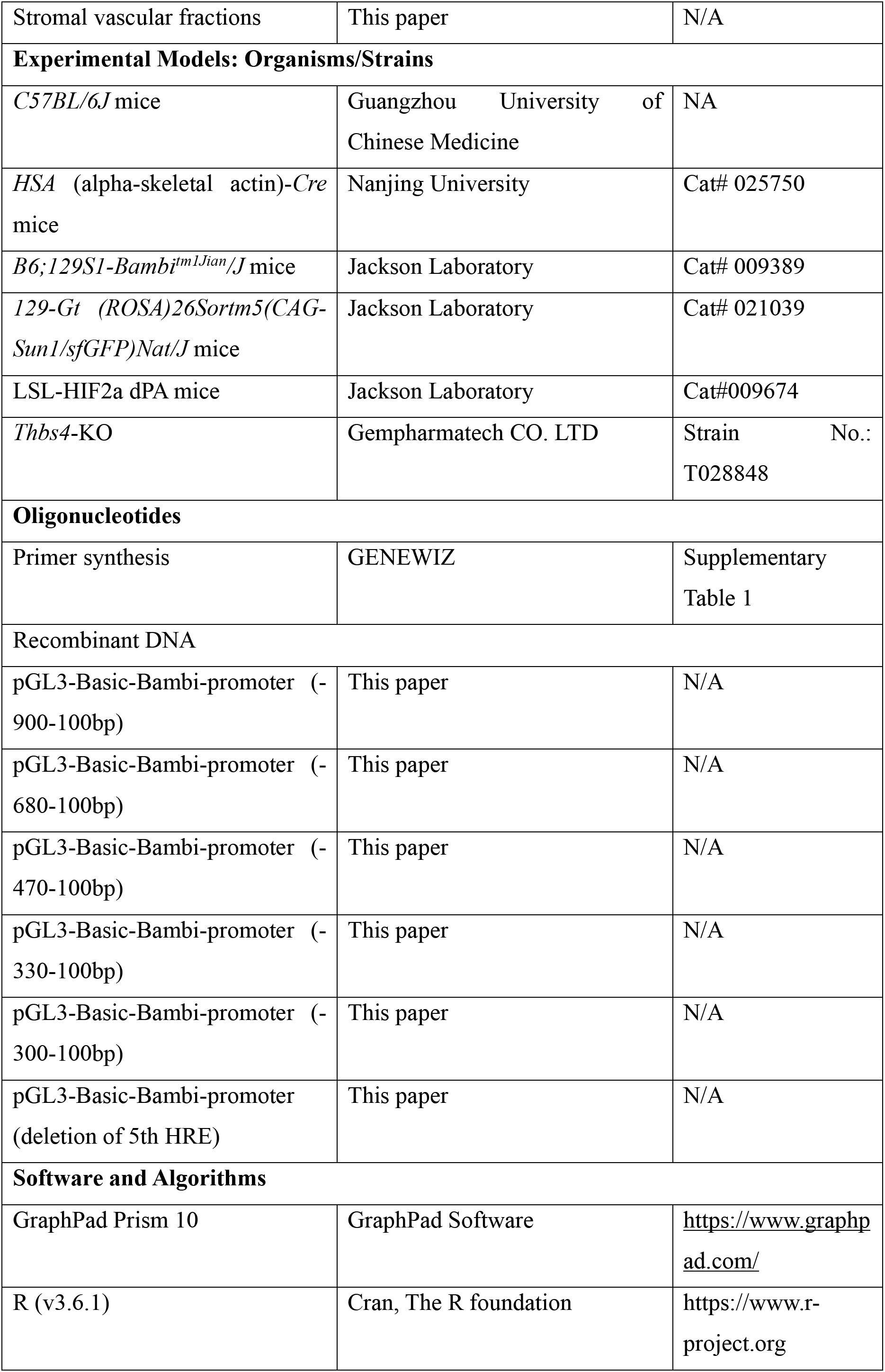
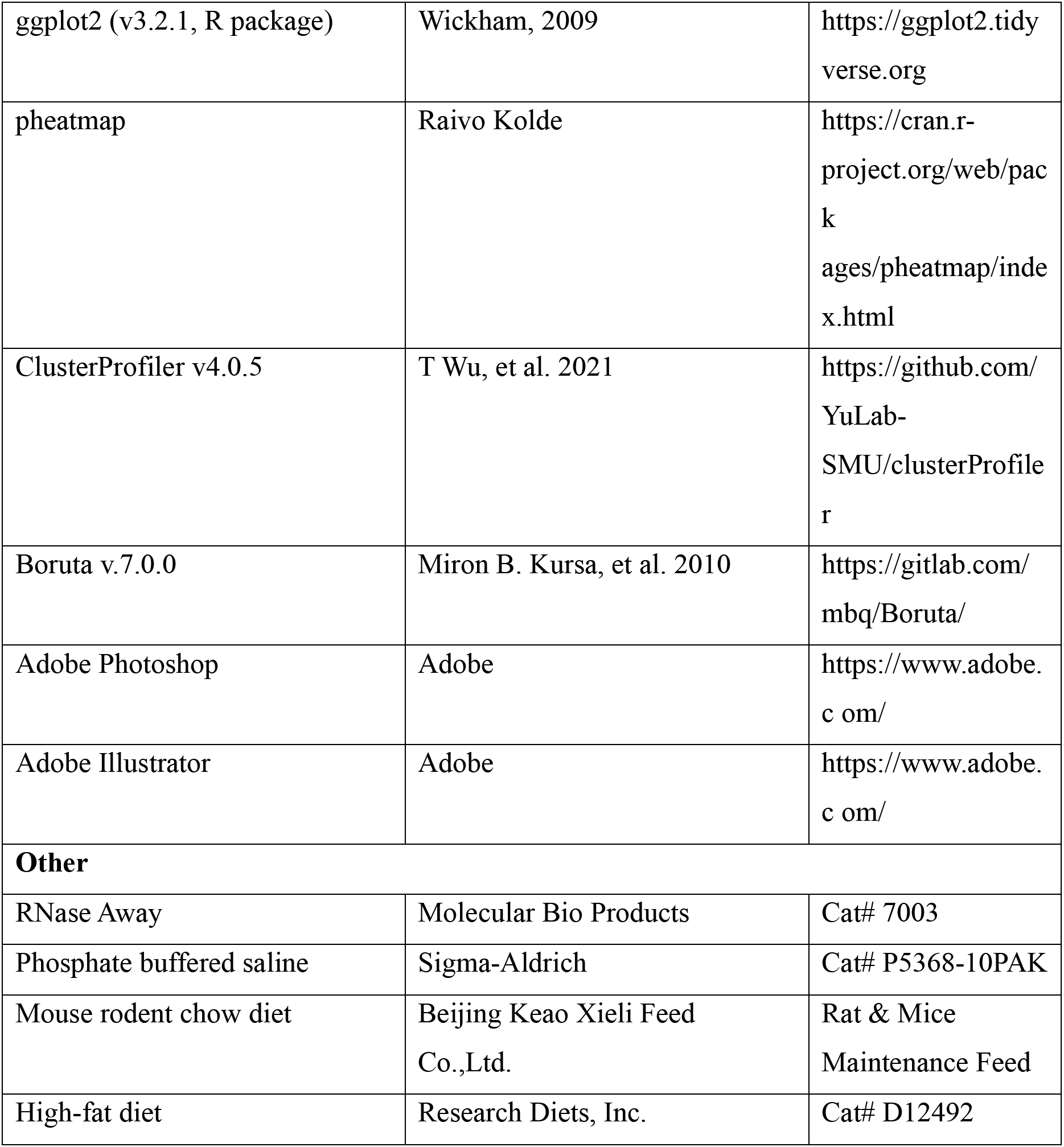

### EXPERIMENTAL MODEL AND SUBJECT DETAILS

#### Animals

*C57BL/6J* mice were purchased from the Guangzhou University of Chinese Medicine Animal Center or Center of Guangdong Experimental Animal Laboratory. *HSA (ACTA, alpha-skeletal actin)-Cre* mice (JAX Stork No.: 025750) were kindly shared by Dr. Zhenji Gan from Nanjing University, *Bambi^fl/fl^* mice (JAX Stork No.: 009389) were shared by Prof. Gongshe Yang from Northwestern Agriculture and Forest University, LSL-HIF2α dPA mice (JAX Stork No.:009674) were kindly shared by Dr. Yatrik Shah from the University of Michigan ^42,87^, the 129-Gt (ROSA)26Sortm5(CAG-Sun1/sfGFP)Nat/J (INTACT, JAX Stork No.: 021039) mice were directly purchased from the Jackson Laboratory, and *Thbs4*-KO (Strain No.: T028848) mice purchased directly from the Gempharmatech CO. LTD. with deletion of exon 3-10 on genome. Heterozygous mice were crossed to generate the homozygous mice, with the wild type mice used as the control littermate. *HSA-Cre* transgenic mice were crossed with *Bambi^fl/fl^*mice carrying flox on *Bambi* gene flanking exon 1 to generate mice with skeletal muscle-specific *Bambi* deletion (annotated as *Bambi^HSA-Cre^*) and LSL-HIF2α dPA (*Hif2α^LSL/LSL^*) mice to excise of a floxed stop cassette enables expression of *Hif2α* cDNA driven by endogenous mouse Gt (ROSA)26Sor promoter. *Bambi^fl/fl^*and *Hif2α^LSL/LSL^* mice were used as the control littermate. To generate mice with skeletal muscle nucleus labeled with GFP, *HSA-Cre* mice were cross with mice carrying INTACT-transgene. To externally induce serum Thbs4 protein level, AAV9-CMV-Thbs4-HA virus were intra-muscular injected into TA muscle to induce Thbs4 overexpression in skeletal muscle. To evaluate serum Thbs4 levels before the downstream experiment, blood was collected from tail vein and confirmed by western blot with HA antibody. Serum Thbs4 levels were measured by using the Thbs4 ELISA kit (EL6512, Cusabio) by following the manufacture’s instruction. All experimental animals were housed in a temperature and humidity controlled and ventilated specific pathogen free (SPF) cages at the animal facility of Institute of Microbiology at Guangdong Academy of Sciences, under a 12-hr/12-hr light and dark cycle (7 a.m.-7 p.m.) at 24 ±2°C and fed a standard irradiated rodent chow diet with free access to food and water *ad libitum*. For cold challenge, the experimental mice were placed in temperature-controlled fridge at 4 ±2°C. For diet induced obesity (DIO)-related experiments, 6-8-week-old mice (body weight around ∼15-20 g) were fed with HFD (60% fat) for at least 20 weeks. All animal handling and procedures were approved by the Animal Care and Use Committee at Institute of Microbiology, Guangdong Academy of Sciences [Permission #: GT-IACUC201704071].

### Method Details

#### Rectal temperature measurement

The rectal temperature was measure by Thermalert (TH-5, Serial No.7329, USA) as described before^88^.

#### Treadmill running

Treadmill running was performed according to established protocol as described before^17^.

#### Cell Culture

HEK293A (CRL-1573™), HEK293T (CRL-3216™), or C2C12 myoblasts (CRL-172™) or were purchased from ATCC. The C2C12 were cultured with DMEM basic medium (C11995500BT, Thermo Fisher) supplemented with 10% fetal bovine serum (FBS, Trinity), and 1% Gibco™ Penicillin-Streptomycin (Thermo Fisher).The primary myoblasts were isolated from the *C57BL6/J* mice as described before^17,88^ and cultured with DMEM/F12 medium (C11330500BT, Thermo Fisher), supplemented with 20% FBS (FBS, Trinity), 1% Gibco™ Penicillin-Streptomycin and 1% recombinant human FGF-basic (154 a.a., 100-18B, PeproTech). To determine the effect of HIF2α on myogenic differentiation, 95% confluent C2C12 or primary myoblasts were infected with adenovirus with expression of *Hif2α* (100 MOI) and 48-hr later, the myoblasts were washed twice with pre-warmed PBS, followed with switching to differentiation medium (DMEM containing 2% horse serum) to induce myogenic differentiation for 5-7 days. For Thbs4 secretion experiment, C2C12 myoblasts were infected with either control or Thbs4 overexpression adenovirus (50 MOI), followed with BFA (an endoplasmic reticulum inhibitor) treatment at concentration of 0, 1,2 μg/ml for 24-hr. The cell culture medium and cell lysates were collected and Thbs4 protein levels were confirmed by western blot.

#### Primary white adipocyte isolation, culture and differentiation

Inguinal white adipose tissue was dissected, minced and digested for 30 min at 37 °C in PBS containing Collagenase II (1.5 mg/mL, A004174, Diamond), Dispase II (2.4 U/mL, D4693, Sigma-Aldrich), and CaCl_2_ (10 mM, 409332, Sigma-Aldrich). The tissue suspension was filtered through a 40 µm cell strainer and centrifuged at 1000 g for 10 min to pellet the stromal vascular fractions (SVFs) and resuspended in growth medium (DMEM containing 10% fetal bovine serum (FBS), and 1% Gibco™ Penicillin-Streptomycin). The cells were plated onto collagen-coated plates overnight before culturing. Preadipocytes were grown to 100% confluence. Primary cells were grown in induction medium (DMEM containing 10% FBS, 1% Gibco™ Penicillin-Streptomycin, 0.5 mM isobutylmethylxanthine (IBMX, I7018, Sigma-Aldrich), 125 μm indomethacin (I7378, Sigma-Alrich), 1 µM dexamethasone (D4902, Sigma-Aldrich), and 0.5 µM rosiglitazone (R2408, Sigma-Aldrich) for 3 days, followed by differentiation medium (DMEM containing 10% FBS, and 1% Gibco™ Penicillin-Streptomycin, 1 μM insulin, VL7516, Eli Lilly and 1 nM triiodothyronine, T2877, Sigma-Aldrich) for 7 days.

#### Seahorse analysis

For metabolic analysis with undifferentiated cells, SVFs isolated from inguinal white adipose tissue were plated on a Seahorse XFe24 cell culture microplate (V7-PS, 100777-004, Agilent) under normal culture condition as described above until 100% confluency. For differentiated adipocyte, SVFs isolated from inguinal white adipose tissue were plated on a 24-well plate under normal culture condition as described above until 100% confluency, followed with switching to the induction medium for 3 days. The induced pre-adipocytes were dissociated with 0.2% Trypsin and counted. ∼20000 pre-adipocytes were plated on a Seahorse XFe24 cell culture microplate (V28, 100882-004, Agilent), followed by 6 days adipocyte differentiation and then mitochondrial stress test. Prior to the mitochondrial stress assay, the Seahorse XF Cartridges was normalized with 1 ml XF base medium (102353-100, Agilent) and incubated overnight in a non-CO_2_ incubator at 37°C. One hour prior the assay, the cells were washed three times with complete XF assay medium, which was supplemented with sodium pyruvate (1 mM), Glutamine (2 mM) and Glucose (10 mM). The cells were then incubated in a non-CO_2_ incubator at 37°C for 50 mins prior the mitochondrial stress test. Oxygen consumption rate (OCR) were measured by the normalized concentration of drugs used in Seahorse XF Cell Mito Stress kit (103015-100, Agilent). Oligomycin (1.5μM, Sigma-Aldrich), FCCP (4 μM, Sigma-Aldrich), Antimyocin/Rotenone (1 μM, Sigma-Aldrich) were pre-loaded into the cartridge and delivered to each well at indicated time points. After the mitochondrial stress test, the protein concentration of each well was measured by BCA Protein Assay Kit (A53225, Thermo Fisher) and the normalized results were calculated with the Seahorse Bioscience Wave Desktop software (Agilent).

#### iTRAQ Proteomics Analysis

Isobaric tags for relative and absolute quantification (iTRAQ) was used to quantitatively screen and determine the secretory proteins expressed at lower levels in serum. The hemoglobin and albumin in the serum of first removed, resulting in enrichment of proteins at lower levels, followed with the standard protocol, following the proteomic enzyme digestion, labeling with iTRAQ reagent (AB Scies, USA), and mass spectrometry analysis with LC-MS/MS (Thermo UltiMate 3000 UHPLC) as described before^88,89^.

#### Body composition

The body composition of *Bambi^HSA-Cre^* and *Bambi^fl/fl^* mice was determined by quantitative NMR (EchoMRI-100H) performed at the College of Animal Science of South China Agricultural University.

#### Glucose tolerance test (GTT) and insulin tolerance test (ITT)

GTT or ITT tests were performed by measuring blood glucose levels with a Yuyue 306 using blood collected from the tail vein. Thus, before each experiment, mice were fasted for 16-hr (GTT: 6 p.m. to 10 a.m.) or 6 hrs. (ITT: 8 a.m. to 2 p.m.). Fasting glucose levels were measured and designated initial glucose levels. For GTT, experimental mice were intraperitoneally (i.p.) injected with 20% D-glucose (2 g/kg fasted body weight) or 10% D-glucose (1g/kg fast body weight for mice fed with HFD), and blood glucose levels were subsequently quantified 15, 30-, 60-, 90- and 120-min post-injection. For ITT, mice were administered an intraperitoneal injection of insulin (VL7516, Eli Lilly) (0.75 U/kg–body weight). Tail blood glucose levels were measured at different time points (15, 30, 60, 90 and 120 min) post *i.p*. administration of insulin.

#### Total RNA isolation and real-time quantitative PCR (qPCR)

Total RNA was extracted from tissues or cells including skeletal muscles, iWAT, iBAT, liver, myoblasts, SVFs or primary adipocytes with TRIzol^TM^ reagent (15596018, Thermo Fisher) according to the manufacturer’s instruction. RNA concentration was measured by NanoDrop absorbance spectroscopy (Thermo Fisher) and reverse transcribed by utilizing 5x All-In-One Master Mix (G490, AbmGood) according to the manufacturer’s instruction. cDNA was used to analyze gene expression by Power SYBR Green Master Mix (A25778, Applied Biosystems) on a QuantStudio 6 Flex Real-Time PCR System (Thermo Fisher). The expression of individual genes was normalized to the expression of 18S ribosomal RNA, a house-keeping gene. The primer sequences for real time qRT-PCR were listed in Supplementary Table 1-qPCR Primer List.

#### INTACT nuclei isolation

INTACT nuclei isolation was performed following procedures developed for SC nuclei isolation described in a previous publication^18^. In short, *HSA-INTACT* mice with GFP labeling on skeletal muscle nuclei were used. Experimental mice were perfused with 4% paraformaldehyde (PFA, P6148, Sigma) via the vascular system to crosslink proteins to chromatin in nuclei, followed by 125 mM glycine to quench the fixation. Skeletal muscles from the hind limbs and trunk were dissected. Skeletal muscle tissues were homogenized (15 ml Wheaton Dounce, loose pestle) in hypotonic buffer (25 mM Tris-base and 25 mM NaCl, pH 7.4). The crude nuclei were collected by centrifuging at 1000 *g* at 4 °C, followed by removing cellular debris by passing through a 40 μm filter. The nuclei were immunoprecipitated with prebound GFP (AE012, ABclonal) and Myc (AE010, ABclonal) antibodies coated with Pierce® Protein A Magnetic Beads (88845, Thermo Fisher) after overnight rotation in a 4 °C freezer. Nuclei bound with beads were isolated on a magnetic stand and washed with PBS 6-10 times to remove the unbound GFP^-^ nuclei. After each round of washing, the purity of nuclei was assessed under a fluorescence microscope (Thermo Fisher).

#### Chromatin immunoprecipitation (ChIP) and sequencing library construction

INTACT-isolated nuclei were sonicated using a Bioruptor^®^ Pico sonication device (B01060010, Diagenode) following a well-established protocol with 30 s on/off for 15 cycles as described in our previous publication^18,88^. DNA fragments of ∼400 bp were generated for subsequent chromatin immunoprecipitation utilizing a TRUE MicroChIP Kit (C01010132, Diagenode) with either IgG or HIF2α antibody, according to the manufacturer’s instructions. The precipitated DNA fragments were purified using the McroChIP DiaPure Columns (C03040001, Diagenode) and used for qPCR analysis or library construction for high-throughput sequencing. The ChIP-seq library was constructed utilizing the MicroPlex Library Preparation Kit (C05010012, Diagenode).

#### Assay for transposase accessible chromatin (ATAC)

Nuclei isolation from skeletal muscle was performed as previously described. Briefly, TA muscle was homogenized using in-house made hypotonic buffer (Tris 25 mM and NaCl 25 mM, pH 7.4) followed by centrifugation and passing through a 40 μm filter to remove cellular debris. A total of 50000 nuclei were counted and used for ATAC experiments with the Tn5 DNA Library Prep Kit for Illumina (K105724, APExBIO). The DNA fragments were purified using MiniElute® PCR Purification Kit (28004, QIAGEN) followed by library preparation using an Index Kit for Illumina (K1058, APExBIO), followed by high-throughput sequencing and bioinformatics.

#### High-throughput sequencing (HTS) and bioinformatic analysis

Total RNA samples were sequenced using a BGI-SEQ2500 platform (Beijing Genomics Institute). ATAC and ChIP samples were sequenced on a HiSeq 4000 platform (Illumina).

RNAseq, ATAC-seq and ChIP-seq raw data in FASTQ format were processed with a quality check using the FASQC algorithm, after which low-quality reads were trimmed using the FASTX-Toolkit. The high-quality RNA-seq reads were further aligned to the mouse genome (GRCm38/mm10) using HISAT2 and assembled against mouse mRNA annotation using HTSeq on a high-performance computational system. Differentially expressed genes (DEGs) were analyzed using the DESeq2 package in R. Genes were considered significantly upregulated or downregulated at *padj*<0.05. For ChIP-seq and ATAC-seq, the high-quality reads were further aligned to the mouse genome (GRCm38/mm10) using Bowtie2.0 with one unique read. The ChIP-seq and ATAC-seq enrichment peaks were called using MACS1.4.2 and annotated using Great 2.0.2. Heatmaps were generated using the heatmap package in R based on the raw count of DEGs. Gene ontology (GO) analysis was performed using the R package ClusterProfiler for DEGs (either up- or downregulated). DEGs (*padj*<0.05) were further analyzed using Gene Set Enrichment Analysis (GSEA). Both upregulated and downregulated genes were functionally categorized using GO and KEGG pathway enrichment analyses. The primers used for ChIP-qPCR were listed in Supplementary Table 2-ChIP-qPCR Primer List.

#### Immunohistochemistry

Adipose tissues were embedded in paraffin, sectioned at 4 μm and mounted onto poly-l-lysine-coated slides. Paraffin slides were deparaffinized in xylene and sequentially hydrated in 100%, 90%, 80%, 70%, and 50% ethanol and then rinsed in ddH_2_O. Antigen retrieval was performed by boiling the sections with Tris-EDTA buffer (Tris 1.21 g and EDTA 0.37 g dissolved in 1 L ddH2O, pH 9.0) for 1 hr. After cooling to R.T. for 1 h, the slides were washed in PBS twice for 5 min. The sections were permeabilized in PBS containing 0.2% Triton X-100 for 10 min. The sections were incubated in 3% H_2_O_2_ in PBS for 10 mins. After washing with PBS, the sections were blocked in 5% normal goat serum in 3% BSA at R.T. for 1 hour and then incubated with primary antibodies against Glut4 (1:200, A7637, ABclonal), Ucp1 (1:500, ab10983, Abcam), PGC1α (1:100, ab3242, Millipore) and Thbs4 (1:200, A16438, ABclonal) overnight at 4 °C in 3% BSA containing 5% normal goat serum. After overnight incubation, the sections were washed 3 times with PBS and subsequently incubated with 1:500 goat anti-rabbit secondary antibody in 3% BSA containing 5% normal goat serum at R.T. for 1 h. The sections were then incubated with DAB reagent (D5905, Sigma-Aldrich) for 3-5 mins. After counterstaining with hematoxylin solution for 2 sec, the sections were dehydrated and mounted using Mounting Medium (Sigma-Aldrich).

### Transmission electron microscopy

TA muscle or iWAT (at dimension of 1 mm^3^) were dissected and immediately fixed in 4% phosphate-glutaraldehyde. Each sample was dehydrated, permeabilized, embedded, sectioned at 60-80 nm and mounted. For each sample, five fields of view were randomly selected and the images were captured with transmission electron microscope (Hitachi), as described before^17^.

#### Immunoblotting

Tissues (*e.g.,* skeletal muscle, iWAT, or iBAT) were homogenized in RIPA buffer containing 150 mm NaCl, 1% NP-40, 0.1% SDS, 25 mm Tris-HCl (pH 7.4), 0.5% sodium deoxycholate, and 100x in DMSO Protease Inhibitor Cocktail EDTA-Free (GK10014, GLPBIO). The protein concentration was measured using a Pierce™ BCA Protein Assay Kit (A53225, Thermo Fisher). Proteins were separated using SDS-polyacrylamide gel electrophoresis (30% Acr-Bis, RM00006, ABclonal) and transferred to PVDF membranes (Merck Millipore). Proteins on the membrane were blocked in 5% (w/v) nonfat milk at R.T. for 1 h, followed by incubation with primary antibodies (Bambi: PA5-38027, Thermo Fisher; β-Tubulin: BBI Life Sciences; CAV1: D161423, Sangon Biotech; PGC1α: AB3242, Millipore; Ucp1: ab10983, Abcam; Glut4: A7637, ABclonal; Thbs4: A16438, ABclonal; MyoD: 39991, DSHB; phospho-AKT, total AKT, phospho-AMPK, total AMPK, ATGL: Cell Signaling). Images were visualized using Ultra High Sensitivity ECL Kit (GK10008, GLPBIO) and were captured using the ChemiDoc^TM^ Imaging System (Bio-Rad).

#### Myofiber isolation and immunofluorescence staining

Single myofibers were isolated from the extensor digitorum longus (EDL) as previously described^17,18,90^. Briefly, dissected EDL muscle was digested in DMEM supplemented with 0.2% Type I Collagenase (LS004214, Worthington) and incubated at 37 °C for 75 min. Single myofibers were isolated in a 60-mm culture dish precoated with horse serum by triturating the digested EDL muscle with polished Pasteur pipettes in washing medium (DMEM supplemented with 10% fetal bovine serum (FBS) and 1% Gibco™ Penicillin-Streptomycin). Freshly isolated myofibers were subsequently fixed in 4% paraformaldehyde (PFA, P6148, Sigma-Aldrich) for 10 mins, quenched with 125 mM glycine, and permeabilized with 0.5% Triton-100, followed by incubation with blocking medium containing 3% BSA and 5% normal goat serum in PBS. Immunostaining of SCs was performed using the following primary antibodies: anti-Pax7 (1:50, Pax7, DSHB), anti-MyoD (1:250; M6190, Sigma-Aldrich) and Thbs4 (1:300, A16438, ABclonal) overnight at 4 °C, followed by secondary antibodies at RT for 1 h in the dark. Slides were mounted with DAPI (F6057, Sigma-Aldrich) and imaged on a confocal microscope (Zeiss 710).

#### Skeletal muscle injury and regeneration

To induce skeletal muscle injury and regeneration, the TA muscle was intramuscularly administrated with cardiotoxin (CTX: BIOTECHNOLOGY, 9012-91-3). TA muscle was harvested either at 30 days post injury (dpi) or at a series of time-points (*e.g.*, 5-, 7- and 9-dpi), followed with the cryosection and immunostaining, as described before^17,18^.

#### Cryosection and immunofluorescence staining

TA muscle was dissected, mounted, frozen, and sectioned at 10 μm as previously described^17,18,40^. Briefly, TA sections on the slide were washed with PBS and incubated in blocking buffer (3% bovine serum albumin, 5% goat serum, 0.2% Triton X-100 and 0.1% sodium azide in PBS) for 30 min. Primary antibodies were incubated overnight at 4 °C (Pax7: Pax7, Lamin B2: 05-206, Millipore; type I: #BA-D5, type IIA: SC-71 dilution with blocking buffer), followed by incubation with secondary antibodies (1:500, diluted in the blocking buffer) at RT for 1 hr in the dark. The sections were washed 5 times with PBS containing 0.5% Tween-20. Nuclei were counterstained with DAPI-containing mounting medium (Life Technologies). Images were visualized and captured using the EVOS Cell Imaging System (Thermo Fisher, Waltham, MA).

#### Plasmid construction

The *Bambi* promoter region was predicted using an online database (The Eukaryotic Promoter Database: https://epd.epfl.ch//index.php). The promoter region flanking 1 kb from −900 to 100 bp upstream of the transcriptional start site (TSS) was PCR-amplified from genomic DNA isolated from mouse liver. The promoter fragment was subsequently cloned into the multiple cloning site at Kpn I (FD0524, Thermo Fisher) and Hind III (FD0504, Thermo Fisher) of the pGL3-basic luciferase constructs. For additional bioinformatics analysis on transcription factor-binding sites, five putative hypoxia response elements (HREs) were predicted. To identify the functional HRE through which HIF2α directly binds to and activates *Bambi* expression, an HRE deletion construct with a shorter promoter fragment was PCR-amplified and subcloned into the pGL-3 basic construct. Constructs expressing HIFs, including W.T. (pcDNA-HA-mHIF1α: #18949, pcDNA-HA-mHIF2α: #18950)^91^ or mutated HIF1α (pcDNA-Myc-HIF1αTM: #44028) and HIF2α (pcDNA-HA-HIF2αTM: #44027)^92^, were directly purchased from Addgene. *Thbs4* expression vectors were constructed by biosynthesizing mouse *Thbs4* coding sequencing (NM_011582.3), which was further inserted into adenovirus or AAV9 constructs. The primers used for cloning were listed in Supplementary Table 3-Cloning Primer List.

#### Luciferase assay

293T cells were transiently co-transfected with the pGL3-*Bambi* promoter and β-gal expression vector as an internal control. To induce mimic hypoxia, after transfection, 293T cells were incubated with CoCl_2_ (200 µM, 409332, Sigma-Aldrich) for 24 hrs ^43,44^. To identify the HIF isoform involved in *Bambi* promoter activation, both W.T. and mutated HIF1/2α (HIF1/2αTM) expression vectors were co-transfected with *Bambi* promoter constructs into 293T cells as described in previous study^42^. A luciferase assay was performed following an internally developed protocol as described before^18,42^. β-Gal levels were measured and used as an internal control. Both luciferase activity and the color matrix of β-gal levels were determined by BioTek CYTATION 5 instrument.

#### Indirect calorimetry

For metabolic studies of *Bambi^HSA-Cre^* and *Bambi^fl/fl^* mice at R.T., mice were housed individually in the Promethion Metabolic Screening Systems (Sable Systems International, North Las Vegas, NV, United States). After adaptation for 24-hr, data of VO_2_ (Oxygen consumption rate), VCO_2_ (Carbon dioxide exhalation rate), and energy expenditure (EE) were monitored and collected for 48 hr. For metabolic studies with the change of temperature, mice in metabolic cage were placed in a temperature-controlled cabinet of Oxymax/CLAMS system (Columbus). After 24-hr adaptation to the system, the data collection of VO_2_, VCO_2_, RER (respiratory exchange ratio) and Heat production were initiated at 6 PM for 48-hr at the R.T., followed by dropping the temperature to 6℃ within 1 hr. The metabolic data were collected for another 48-hr at the cold. These mice were with free access to food and water *ad libitum* and under an environment similar to the animal facility with environment temperature (23-25 °C) and a 12-h light/dark cycle.

## Statistical analysis

Experimental data were presented as the mean ± SEM. Data were analyzed using the unpaired Student^’^s t test or one-way ANOVA, and a *p-value* < 0.05 was considered statistically significant. Bar plots and statistical analysis were generated using GraphPad Prism 10. Representation of *p-*values is shown as \**p* < 0.05, \*\**p* < 0.01, \*\*\**p* < 0.001 and NS: not significant (*p* > 0.05).

## Acknowledgments

The journey of our six-year research project has been enriched by the relentless dedication and invaluable contributions from our authors, participants, and collaborators, forming the backbone of this significant scientific endeavor. Dr. Xiangping Yao, who initiated and steered this complex project during her Ph.D. and nurtured it through her postdoctoral period, deserves our profound admiration and appreciation for her vision and her pioneering efforts. The spotlight also shines brightly on Dr. Xudong Mai, whose profound contributions during the revision stage and his exceptional aptitude for devising advanced experiments and troubleshooting complex issues, have been the lifeblood of our research. His relentless pursuit of excellence and his ceaseless innovation have significantly enhanced our study’s standing. Our esteemed colleagues, Ms. Ye Tian, Ms. Yifan Liu, and Ms. Guihua Pan, deserve a special note of acknowledgement for their commendable work. Their meticulous precision in pathological section processing and imaging, their dedication to animal breeding, and their thorough commitment to scientific inquiry have been crucial in shaping our research. Ms. Liujing Huang’s formidable prowess in bioinformatics has been a cornerstone in our project. Her deftness in script development and data processing across multi-omics data has greatly advanced our understanding of our findings. Our deep appreciation extends to Dr. Jingxing Ou and his Ph.D. student, Guanghui Jin. Their unwavering involvement, especially in the metabolic cage experiment during the manuscript revision, has added a deeper layer of richness to our study. Their professional dedication and academic proficiency have greatly elevated the depth and quality of our research. To all lab members who have shown diligent dedication in reagent preparation, cell model generation, and testing procedures, your unwavering efforts have proven to be a linchpin in the timely completion of our revisions. Our heartfelt thanks go out to our wider circle of friends and colleagues, who have consistently provided insightful discussions and constructive suggestions, enriching our research journey. Each author’s family deserves special acknowledgement for their enduring support, demonstrating the invaluable role of personal encouragement in the successful execution of a comprehensive, long-term project.

## Author Contribution

Experimentation: XP. Y., XD. M., Y.T., SJ. C., XS. D., XJ. J., ZY. L., GH. J., Z. L., ZJ. F., ML. H., LL. X., GH. P., J. S., and H. C. During the manuscript revision, JX. O. and GH. J. provided fully support of metabolic cage experiment, data interpretation and discussion. Analysis: LJ. H, and XH. P.; Manuscript preparation: XP. Y., and LW. X.; Manuscript revision: LW. X., JX. O., XD. M., H. C. Supervision: LW. X.; Funding acquisition: LW. X. All authors reviewed the manuscript and approve the submission of the revised manuscript.

## Lead Contact and Materials Availability

Further information and requests for resources and reagents should be directed to and will be fulfilled by the Lead Contact, Liwei Xie. All unique/stable reagents generated in this study are available from the Lead Contact with a completed Materials Transfer Agreement.

## Data and Code Availability

RNAseq, ChIP-seq and ATAC-seq sequencing data were deposited at National Center for Biotechnology Information with project number of PRJNA819493 (https://www.ncbi.nlm.nih.gov/bioproject/?term=PRJNA819493).

## Supplementary figure legends

**Figure S1.**
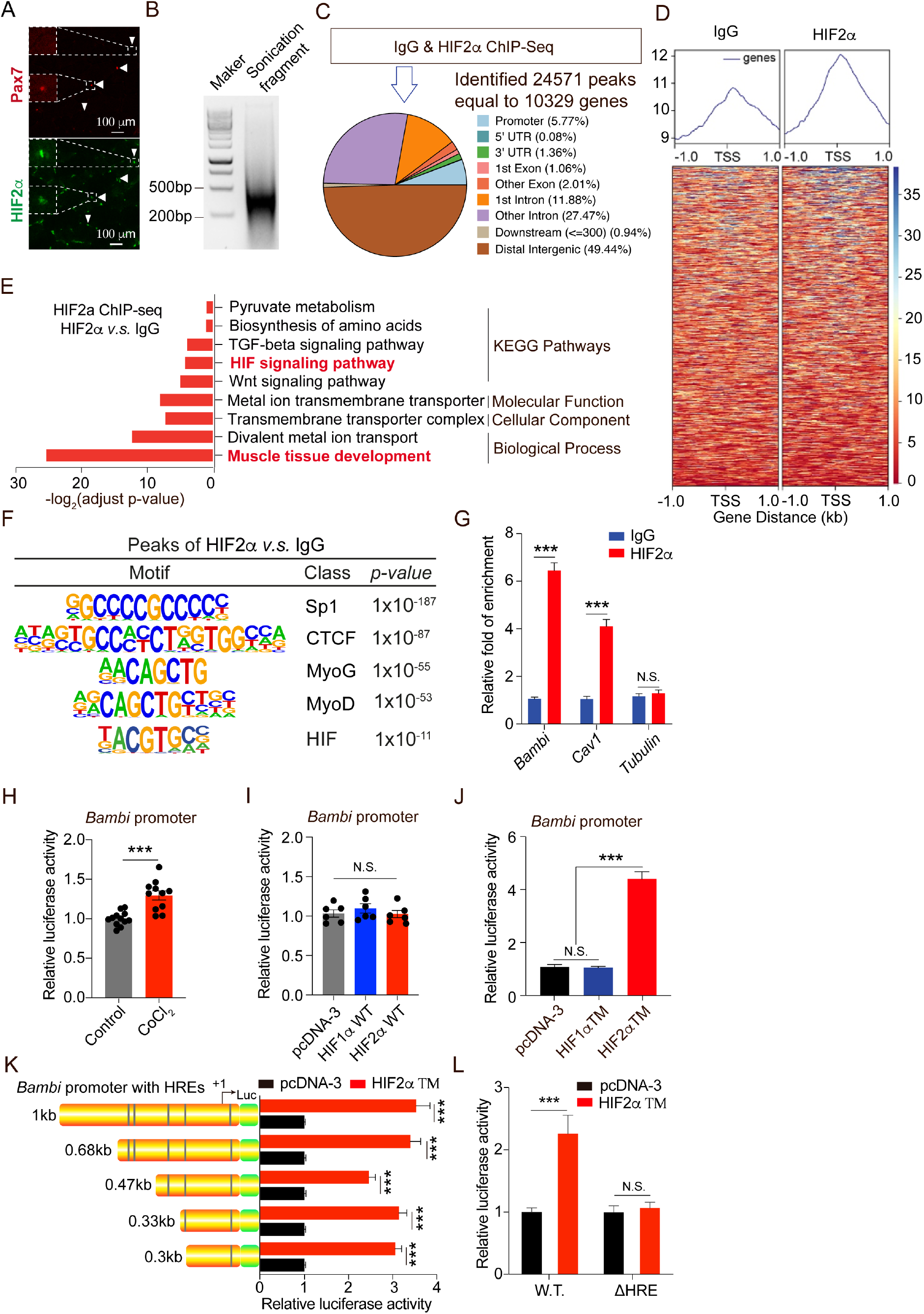
Identification of *Bambi* as a direct target of HIF2α. (A) Representative images of Pax7 (Red), HIF2α (Green) and DAPI (Blue) immunofluorescence staining in TA muscle cryosections from *C57BL/6J* mice (n=5 mice). (B) Image of the DNA agarose gel showing the size of the DNA fragment after sonication. (C) The proportion of the genomic distribution of HIF2α-bound DNA fragments in the myonuclei of skeletal muscle. (D) Heatmap of HIF2α-bound DNA fragment enrichment flanking a 1 kb region of TSS between IgG and HIF2α. (E) Gene ontology (GO) and KEGG pathway analysis of annotated genes associated with HIF2α-bound DNA fragments. (F) Motif enrichment analysis of the peaks aligned in the promoter region. (G) ChIP-qPCR indicated that HIF2α directly binds to the promoter region of *Bambi* and *Cav1* (a known HIF2α target as a positive control), with the Tubulin promoter shown as a negative control (n=3). (H) Luciferase assays of *Bambi* promoter activity under CoCl_2_ treatment (n=6). (I) Luciferase assays of *Bambi* promoter activity with co-transfection of construct expressing wild type HIF1α or HIF2α (n=6). (J) Luciferase assays of *Bambi* promoter activity with co-transfection of construct expressing mutant HIF1α (HIF1αTM) or HIF2α (HIF2αTM) (n=6). (K) Luciferase assay of truncated *Bambi* promoter activity, containing different numbers of HREs, under overexpression of HIF2αTM. (L) Luciferase assay of 300 bp *Bambi* promoter bearing wild type and deleted HRE with overexpression of HIF2αTM. The luciferase activity induced by HIF2αTM was abolished upon the deletion of HRE in the −300 bp *Bambi* promoter construct (n=6). N.S., not significant, \*\*\**p* < 0.005, by 2-sided Student’s t test. Data represent the mean ± SEM.

**Figure S2 for Figure S1.**
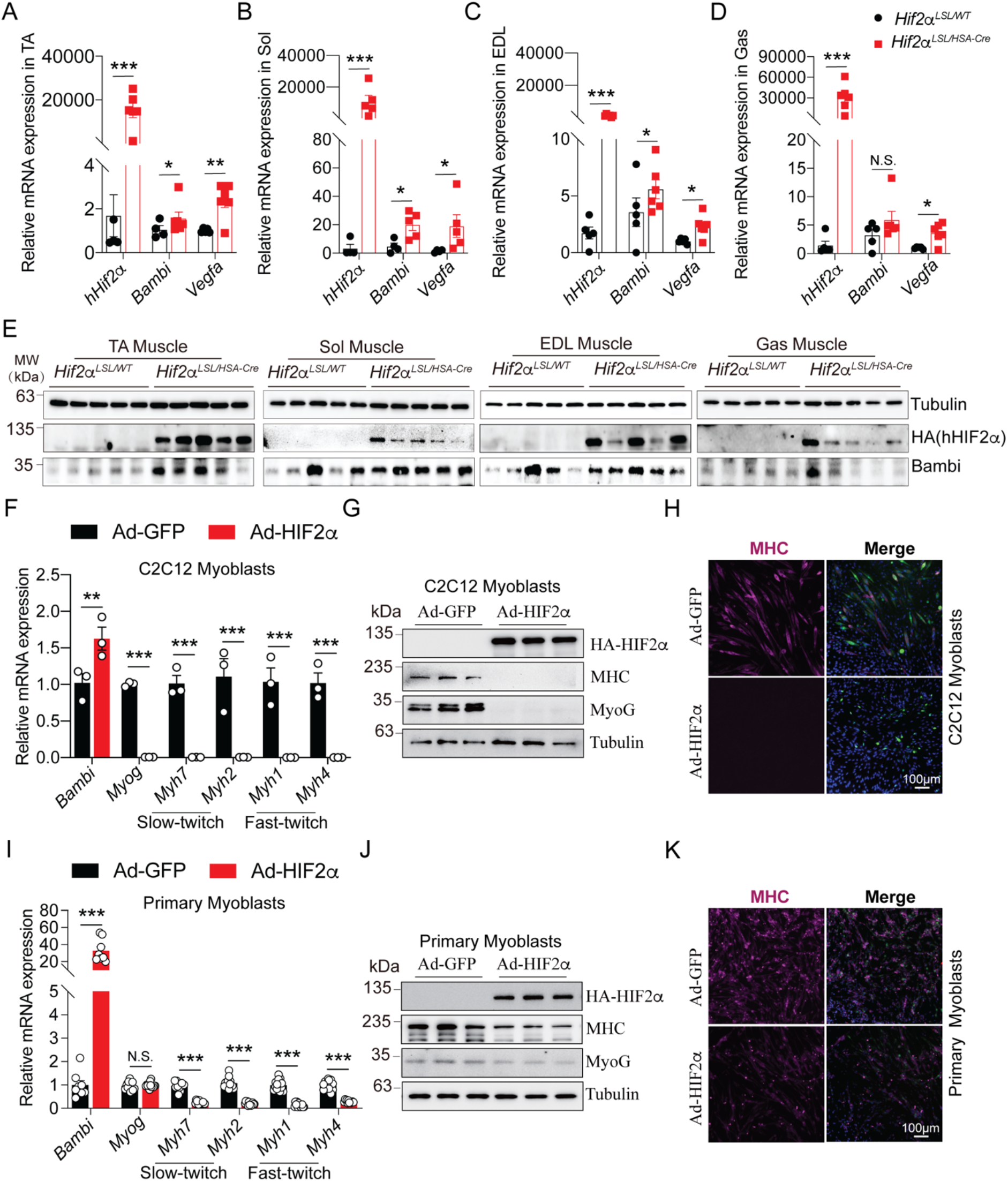
Identification of *Bambi* as a direct target of HIF2α. (A-D) qPCR analysis of *hHif2α*, *Bambi* and *Vegfa* mRNA expression in TA (A), Sol (B), EDL (C), and Gas (D) muscle of *Hif2α^LSL/WT^* and *Hif2α^LSL/HSA-Cre^* mice. (E) Representative immunoblots of hHIF2α-HA, Bambi and Tubulin in TA, Sol, EDL, and Gas muscle of *Hif2α^LSL/WT^* and *Hif2α^LSL/HSA-Cre^* mice. (F) qPCR analysis of *Bambi*, *Myog*, *Myh7*, *Myh2*, *Myh1* and *Myh4* mRNA expression in C2C12 myoblasts infected with either Ctrl (Ad-GFP) or HIF2α-expressing adenovirus infection (Ad-HIF2α). (G) Representative immunoblots of HIF2α-HA, MHC and MyoG protein expression level in C2C12 myoblasts. (H) Representative images of MHC (Purple) immunostaining of C2C12 myoblasts. (I) qPCR analysis of *Bambi*, *Myog*, *Myh7*, *Myh2*, *Myh1* and *Myh4* mRNA expression in primary myoblasts infected with either Ctrl (Ad-GFP) or HIF2α-expressing adenovirus infection (Ad-HIF2α). (J) Representative immunoblots of HIF2α-HA, MHC and MyoG protein expression level in primary myoblasts. (K) Representative images of MyoG (Red) and MHC (Purple) immunostaining of primary myoblasts. N.S., not significant, \**p* < 0.05, \*\**p* < 0.01, and \*\*\**p* < 0.001 by two-sided Student’s t test. Data represent the mean ± standard error of the mean.

**Figure S3 for Figure 1.**
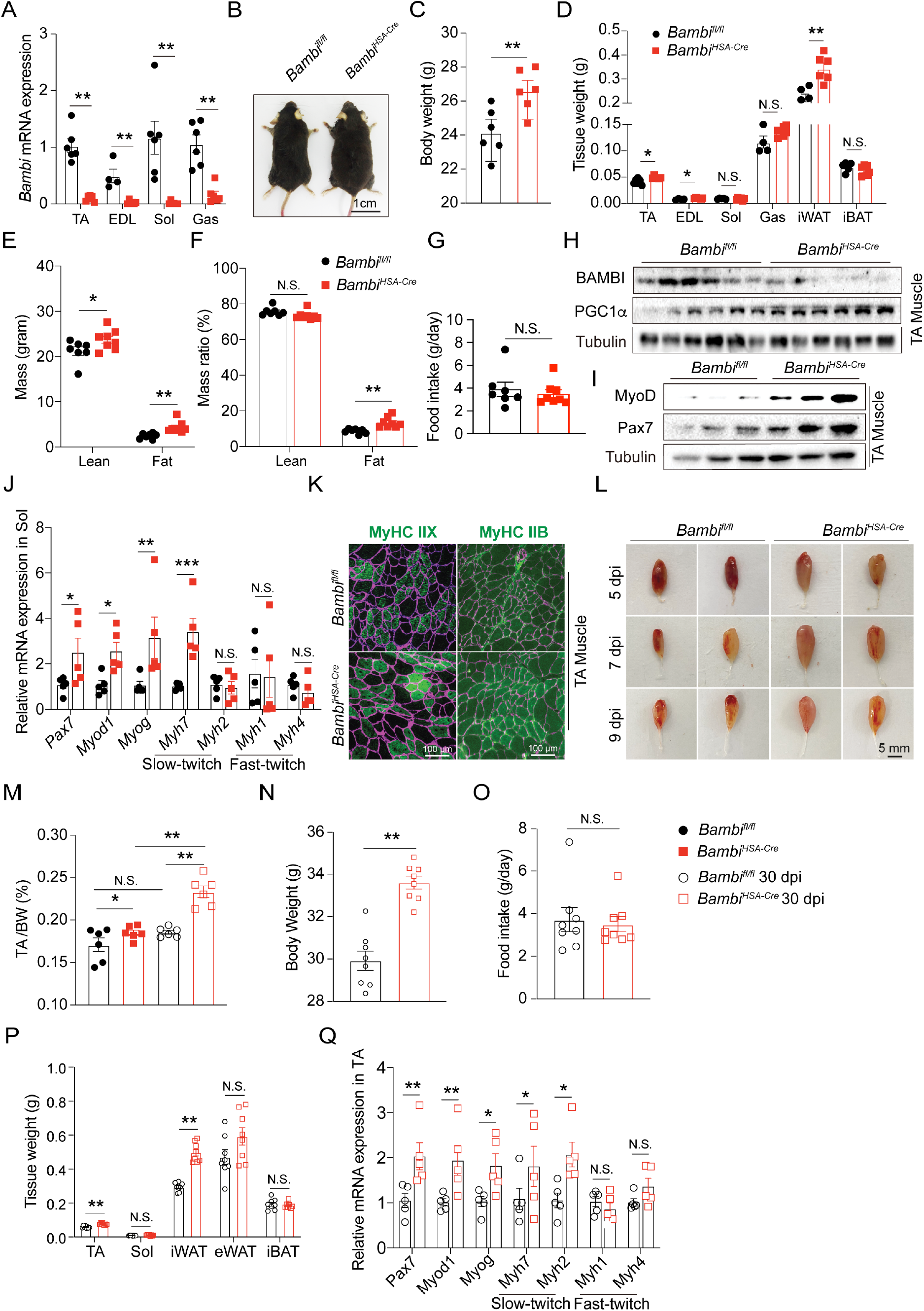
Muscle-specific deletion of *Bambi* induces muscle hypertrophy. (A) qPCR analysis of *Bambi* expression in skeletal muscles (TA, EDL, Sol and Gas) between *Bambi^fl/fl^* and *Bambi^HSA-Cre^* mice (n=6 mice/group). (B) Representative images of *Bambi^fl/fl^* and *Bambi^HSA-Cre^* mice on rodent chow diet (n=6 mice/group). (C) Body weight of adult of *Bambi^fl/fl^*and *Bambi^HSA-Cre^* mice. (D) The tissue weight of TA, EDL, Sol, Gas, iWAT and BAT (n=6 mice/group). (E-F) DEXA scanning of lean and fat mass. (G) Daily food intake. (H) Representative immunoblots of Bambi and PGC1α protein levels in the TA muscle (n=6 mice/group). (I) Representative immunoblots of MyoD, Pax7 and Tubulin in TA muscle (n=6/group). (J) qPCR analysis of gene expression in Sol muscle (n=5 mice/group). (K) Representative images of MyHC IIX/IIB (Green) and LaminB2 (Purple) immunofluorescence staining of TA muscle cryosections. (L) Representative images of TA dissected from *Bambi^fl/fl^*and *Bambi^HSA-Cre^* mice post injury (5, 7, and 9 dpi) (n=6). (M) The ratio of TA muscle/body weight between *Bambi^fl/fl^* and *Bambi^HSA-Cre^* mice before and 30 days post-CTX injury. (N) Body weight of *Bambi^fl/fl^*and *Bambi^HSA-Cre^* mice 30 dpi. (O) Daily food intake in *Bambi^fl/fl^*and *Bambi^HSA-Cre^* mice 30 dpi. (P) The tissue weight (TA, Sol, iBAT, iWAT and eWAT) of *Bambi^fl/fl^* and *Bambi^HSA-Cre^* mice 30 dpi. (Q) qPCR analysis of gene expression in TA muscle samples from *Bambi^fl/fl^* and *Bambi^HSA-Cre^* mice (n=5 mice/group) 30 dpi. N.S., not significant, \**p* < 0.05, and \*\**p* < 0.01 by two-sided Student’s t test. Data represent the mean ± standard error of the mean.

**Figure S4 for Figure 2.**
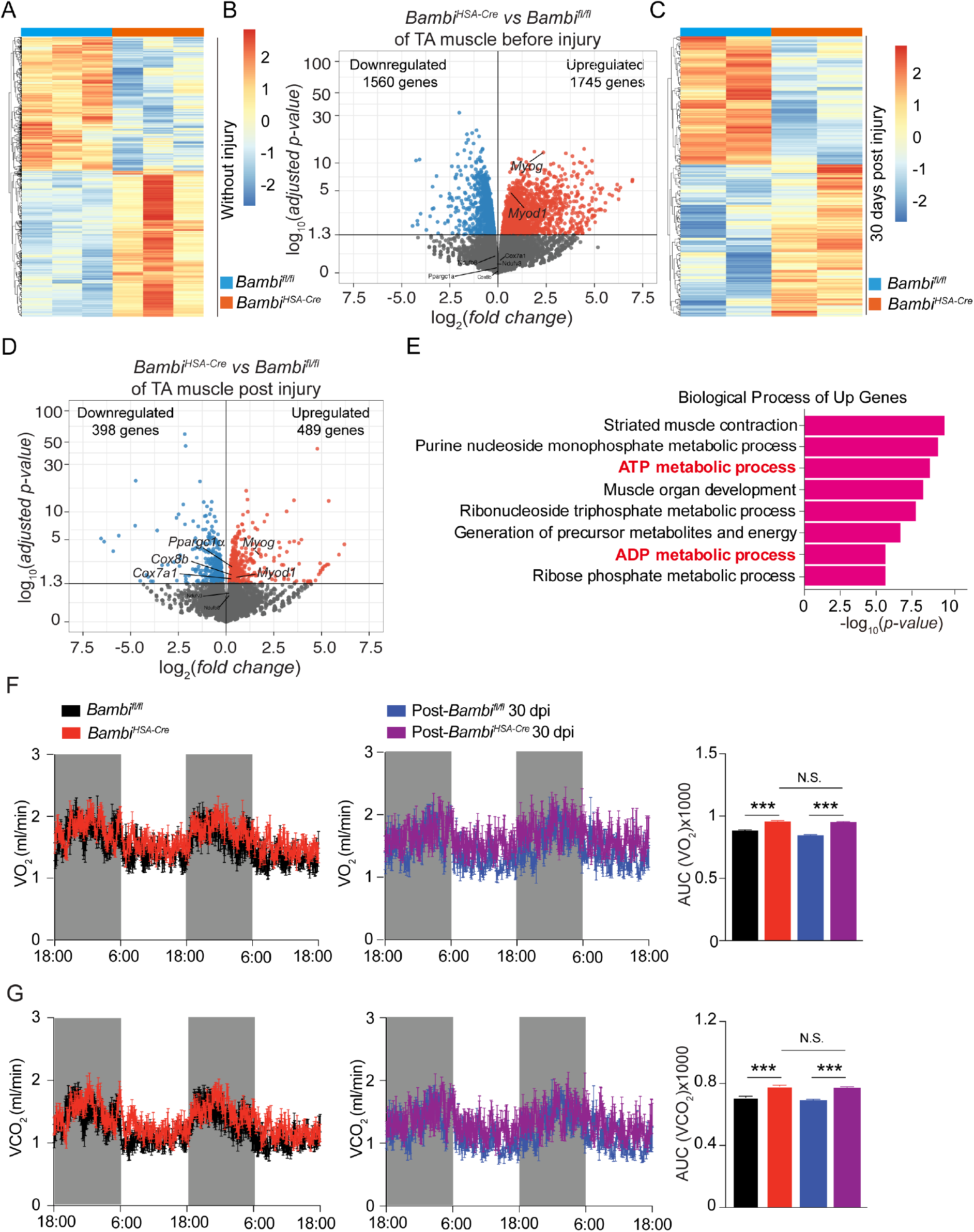
*Bambi* deletion-induced muscle hypertrophy promotes systemic energy metabolism. (A) Heatmap of the gene expression profile in TA muscle between *Bambi^fl/fl^* and *Bambi^HSA-Cre^* mice without injury (n=3 mice/group). (B) Volcano plot of DEGs in TA muscle between *Bambi^fl/fl^* and *Bambi^HSA-Cre^* mice without injury. (C) Heatmap of the gene expression profile in TA muscle between *Bambi^fl/fl^* and *Bambi^HSA-Cre^* mice 30 days post injury (n=3 mice/group). (D) Volcano plot of DEGs in TA muscle between *Bambi^fl/fl^* and *Bambi^HSA-Cre^* mice 30 days post injury. (E) GO analysis (Biological Process) of upregulated DEGs in TA muscle of *Bambi^fl/fl^*and *Bambi^HSA-Cre^* mice 30 days post injury. (F) Average VO_2_ (Oxygen consumption rate) was monitored over a 48-hr period for *Bambi^fl/fl^* and *Bambi^HSA-Cre^*mice before and 30 days post injury. (G) Average VCO_2_ was monitored over a 48-hr period for *Bambi^fl/fl^* and *Bambi^HSA-Cre^* mice before and 30 days post injury. N.S., not significant, \*\*\**p* < 0.001 by two-sided Student’s t test. Data represent the mean ± standard error of the mean.

**Figure S5 for Figure 3.**
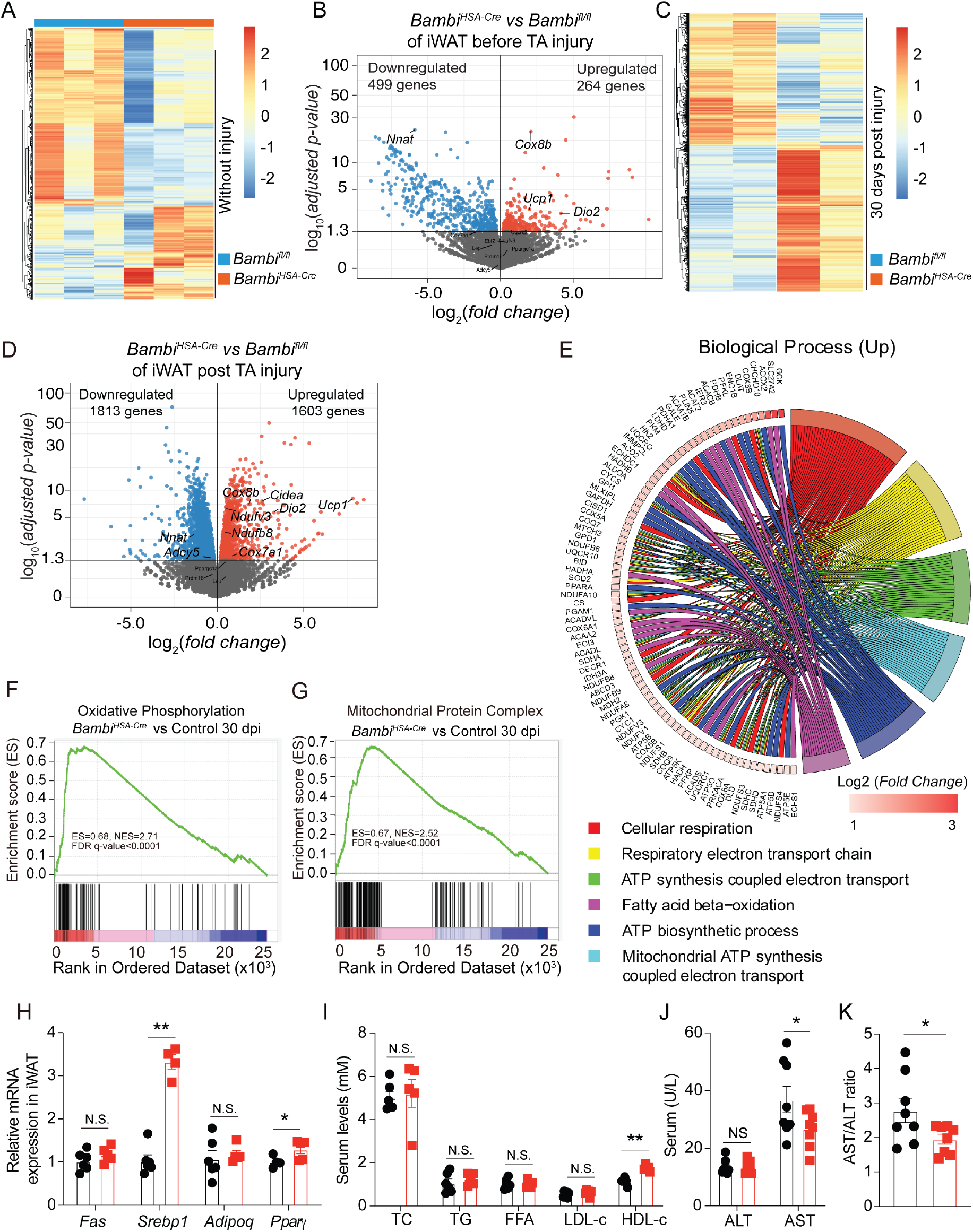
*Bambi* deletion in skeletal muscle activates the thermogenic capacity of iWAT. (A) Heatmap of the gene expression profile in iWAT between *Bambi^fl/fl^* and *Bambi^HSA-Cre^* mice without injury (n=3 mice/group). (B) Volcano plot of differentially expressed genes in iWAT between *Bambi^fl/fl^*and *Bambi^HSA-Cre^* mice without injury. (C) Heatmap of gene expression in iWAT between *Bambi^fl/fl^* and *Bambi^HSA-Cre^* mice 30 days post injury. (D) Volcano plot of differentially expressed genes in iWAT between *Bambi^fl/fl^* and *Bambi^HSA-Cre^* mice 30 days post injury. (E) GO analysis (Biological Process of upregulated genes in iWAT of *Bambi^fl/fl^* and *Bambi^HSA-Cre^*mice 30 days post injury. (F-G) GSEA of significantly upregulated pathways (oxidative phosphorylation and mitochondrial protein complex). (H) qPCR analysis of gene expression in iWAT in *Bambi^fl/fl^* and *Bambi^HSA-Cre^* mice 30 days post injury. (I) Serum levels of TC, TG, FFA, LDL-c and HDL-c in *Bambi^fl/fl^* and *Bambi^HSA-Cre^*mice 30 days post injury. (J) Serum levels of ALT and AST in *Bambi^fl/fl^* and *Bambi^HSA-Cre^* mice 30 days post injury. (K) Ratio of ALT and AST. N.S., not significant, \**p* < 0.05, and \*\**p* < 0.01 by two-sided Student’s t test. Data represent the mean ± standard error of the mean.

**Figure S6 for Figure 4.**
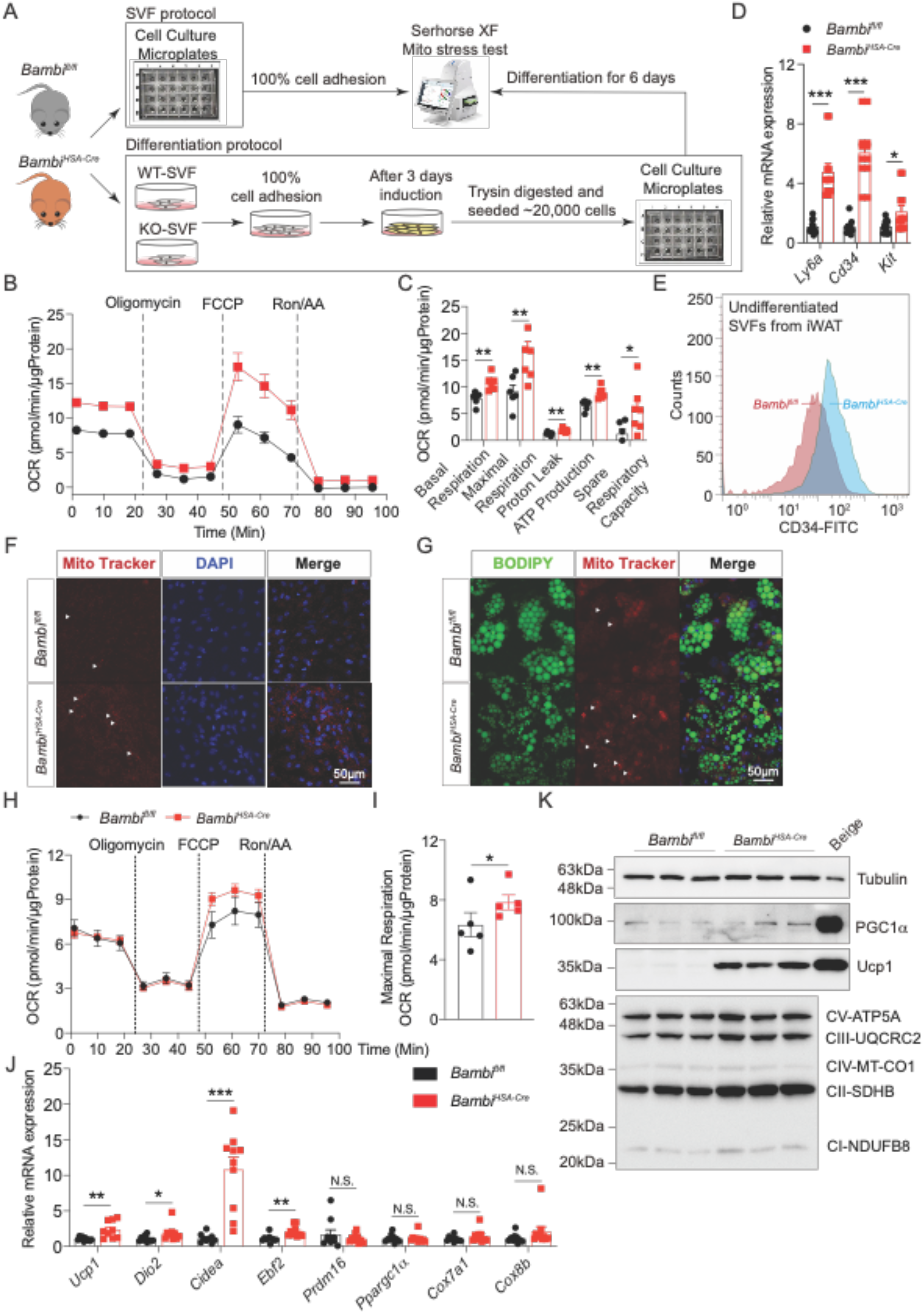
Mice with *Bambi* deletion in skeletal muscle resistant to HFD-induced metabolic disorders. (A) Diagram depicting the timeline of seahorse flux experiment on undifferentiated SVFs isolated from iWAT or differentiated primary white adipocytes of *Bambi^fl/fl^* and *Bambi^HSA-Cre^*mice. (B) Seahorse flux analysis of OCR of undifferentiated SVFs isolated from iWAT of *Bambi^fl/fl^* and *Bambi^HSA-Cre^* mice (n=3 independent repeat). (C) Statistical analysis of OCR at the stage of basal respiration, maximal respiration, proton leaky, ATP production and spare respiration capacity. (D) qPCR analysis of *Ly6a*, *Cd34*, *Kit* mRNA expression levels in undifferentiated SVFs from iWAT of *Bambi^fl/fl^*and *Bambi^HSA-Cre^* mice (n=12). (E) Flow cytometry analysis of CD34^+^ SVFs from iWAT of *Bambi^fl/fl^* and *Bambi^HSA-Cre^*mice. (F) Representative images of Mito-Tracker (Red) immunostaining of undifferentiated SVFs from iWAT of *Bambi^fl/fl^* and *Bambi^HSA-Cre^* mice. (G) Representative images of BODIPY (Green) and Mito-Tracker (Red) immunostaining of differentiated primary white adipocytes of *Bambi^fl/fl^*and *Bambi^HSA-Cre^* mice. (H) Seahorse flux analysis of OCR of differentiated primary white adipocytes of *Bambi^fl/fl^* and *Bambi^HSA-Cre^* mice (n=3 independent repeat). (I) Statistical analysis of OCR at the stage of maximal respiration. (J) qPCR analysis of thermogenesis-related gene expression in differentiated primary white adipocytes of *Bambi^fl/fl^* and *Bambi^HSA-Cre^* mice (n=8). (K) Representative immunoblots of Ucp1, PGC1α, Mito-complex and Tubulin in differentiated primary white adipocytes of *Bambi^fl/fl^* and *Bambi^HSA-Cre^*mice (n=3 independent repeat). N.S., not significant, \**p* < 0.05, \*\**p* < 0.01, and \*\*\**p* < 0.001 by two-sided Student’s t test. Data represent the mean ± standard error of the mean.

**Figure S7 for Figure 5.**
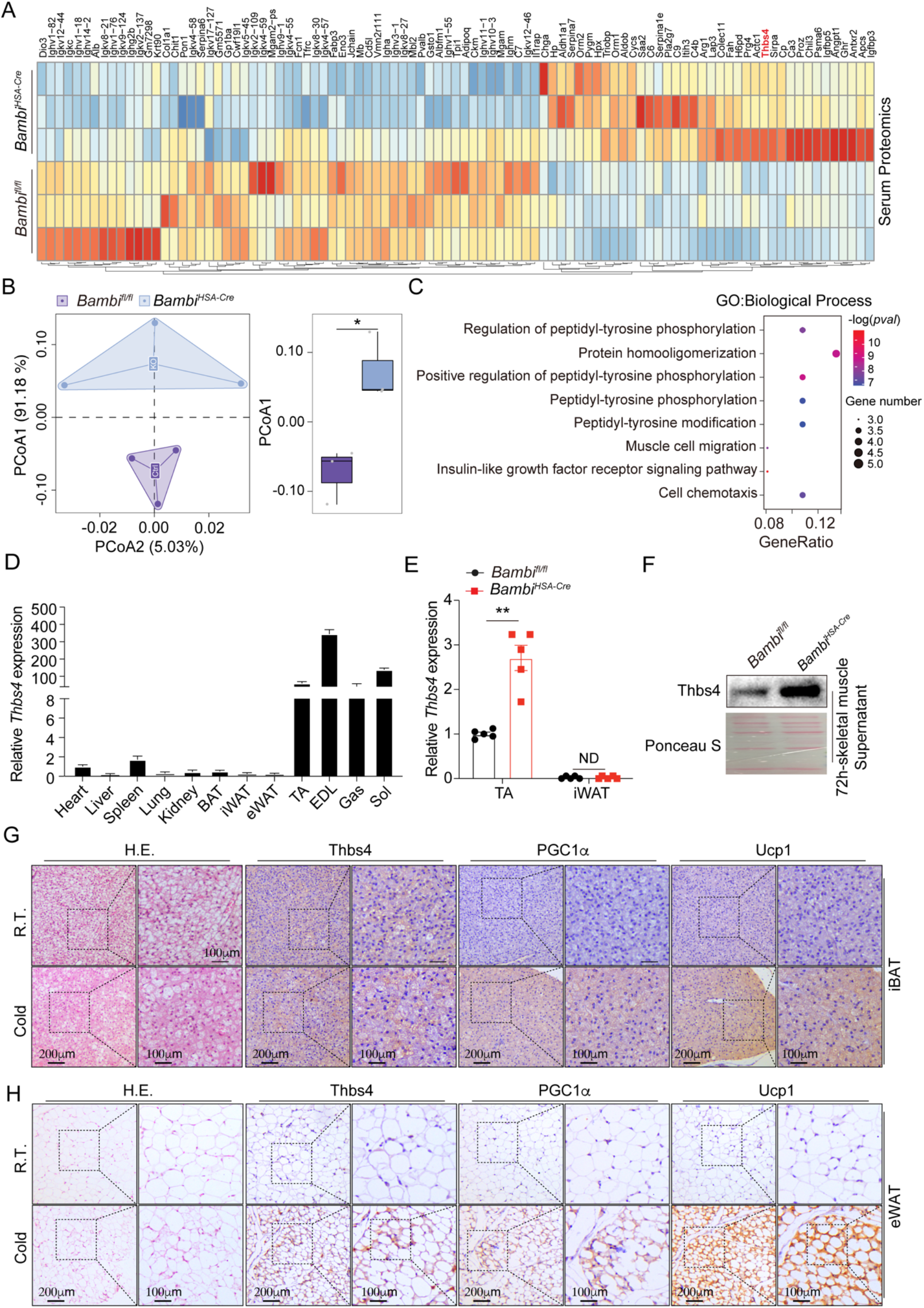
*Bambi* deletion-induced hypertrophy enhances Thbs4 expression and secretion. (A) Heatmap of serum proteomics of *Bambi^fl/fl^* and *Bambi^HSA-Cre^* mice. (B) Principal coordinate analysis (PCoA) of serum proteomics of *Bambi^fl/fl^* and *Bambi^HSA-Cre^* mice. (C) GO analysis (Biological Process) of differentially upregulated protein biomarkers of *Bambi^fl/fl^* and *Bambi^HSA-Cre^* mice. (D) qPCR analysis of *Thbs4* expression in different tissues of *C57BL/6J* mice. (E) qPCR analysis of *Thbs4* expression in TA and iWAT of *Bambi^fl/fl^*and *Bambi^HSA-Cre^* mice (n=5 mice/group). (F) Representative immunoblots of Thbs4 protein levels in the supernatant of *ex vivo* cultured TA muscle for 72-hr of *Bambi^fl/fl^* and *Bambi^HSA-Cre^*mice. (G) Representative images of H&E staining. and IHC of Thbs4, PGC1α and Ucp1 in iBAT sections from the RT and cold challenge groups. (H) Representative images of H&E staining. and IHC of Thbs4, PGC1α and Ucp1 in eWAT sections at the condition of R.T. or cold challenge for 7 days. ND, not detected, N.S., not significant, \**p* < 0.05, and \*\**p* < 0.01 by two-sided Student’s t test. Data represent the mean ± standard error of the mean.

**Figure S8 for Figure 5.**
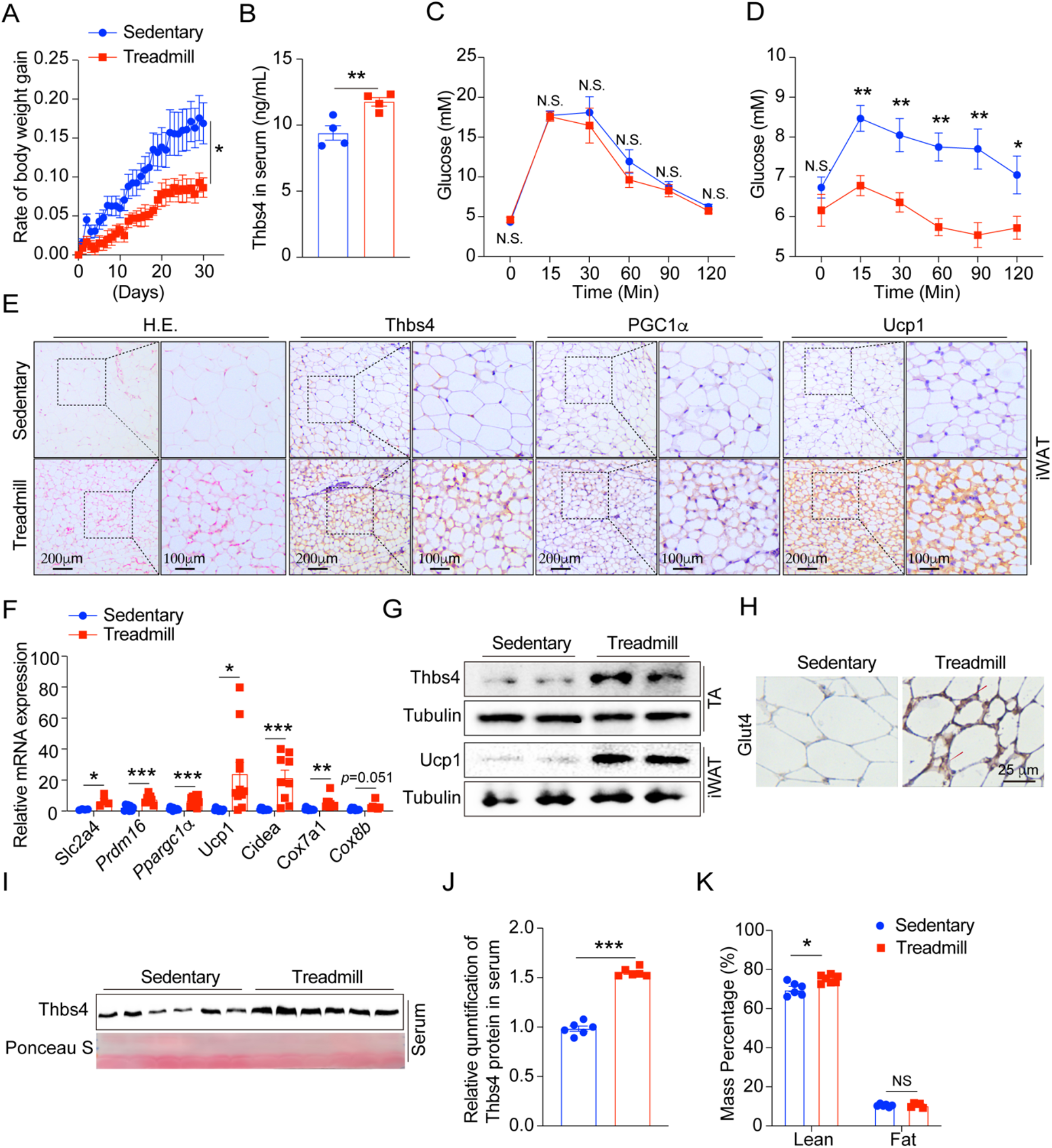
*Bambi* deletion-induced hypertrophy enhances Thbs4 expression and secretion. (A) The rate of body weight gain in the sedentary and 30-day aerobic training of treadmill running groups (n=4 mice/group of two independent repeat). (B) Serum Thbs4 levels in the sedentary and treadmill running groups. (C-D) GTT and ITT for the sedentary and treadmill running groups. (E) Representative images of H&E staining and IHC staining of Thbs4, PGC1α and Ucp1 antibodies in the iWAT of mice in the sedentary and treadmill running groups. (F) qPCR analysis of thermogenesis-related gene expression in the iWAT of mice in the sedentary and treadmill running groups. (G) Representative immunoblots of Thbs4 (TA muscle) and Ucp1 (iWAT) in the sedentary and treadmill running groups. (H) Representative images of IHC staining of Glut4 in the iWAT of mice in the sedentary and treadmill running groups. (I) Representative immunoblots of Thbs4 protein levels in serum in the sedentary and treadmill running groups. (J) Serum Thbs4 protein levels in the sedentary and treadmill running groups. (K) The percentage of lean mass and fat mass in the sedentary and treadmill running groups. N.S., not significant, \**p* < 0.05, \*\**p* < 0.01 and \*\*\**p* < 0.001, by two-sided Student’s t test. Data represent the mean ± standard error of the mean.

**Figure S9 for Figure 5.**
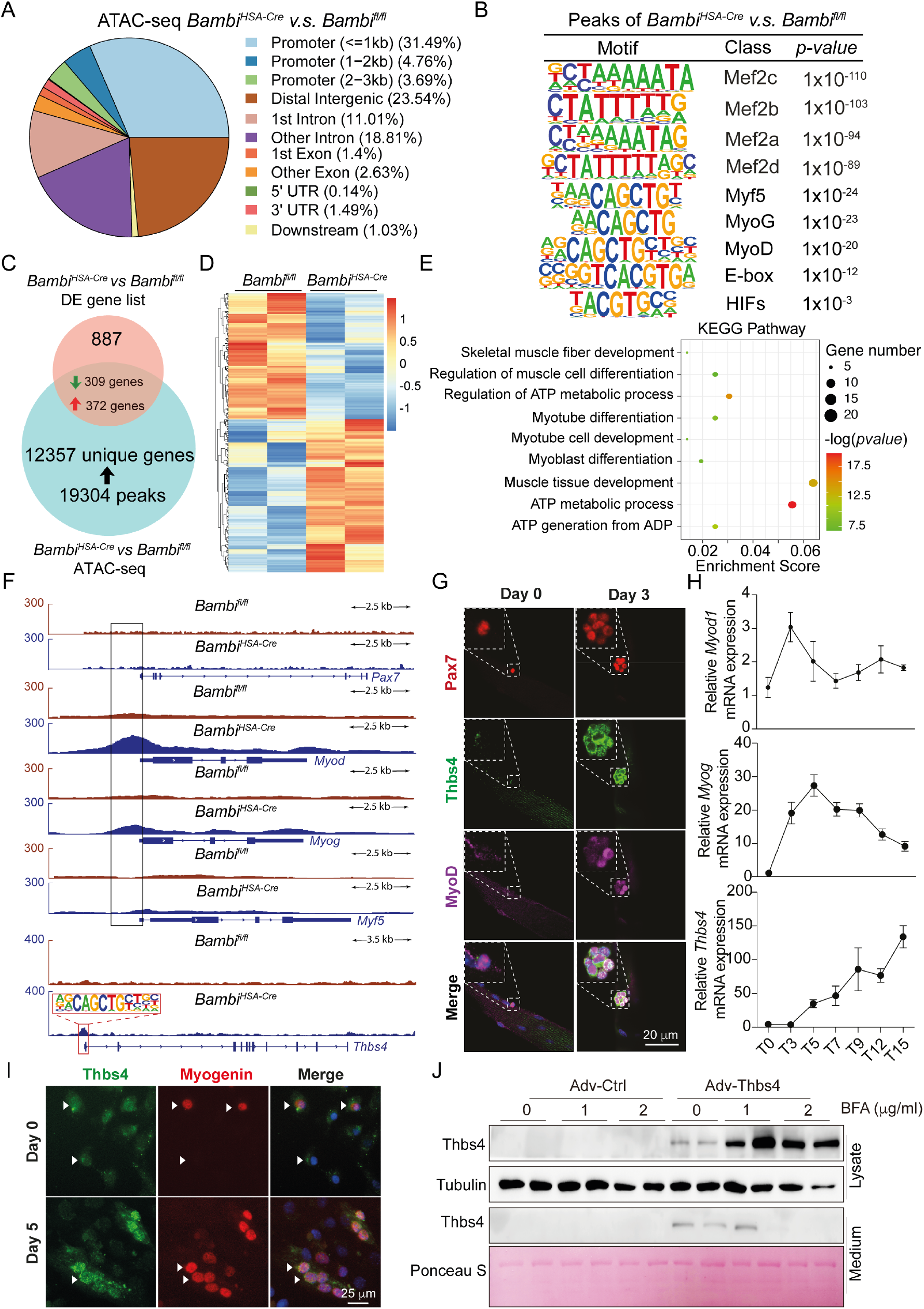
*Bambi* deletion-induced hypertrophy enhances Thbs4 expression and secretion. (A) Distribution of ATAC-seq peaks in the whole genome across promoters, introns, exon distal intergenic regions and UTRs. (B) Motif enrichment analysis of the peaks aligned in the chromatin opening region. (C) Venn diagrams of overlapping DEGs between RNA-seq (between *Bambi^fl/fl^* and *Bambi^HSA-Cre^* mice) and ATAC-seq (peaks associated unique genes). (D) Heatmap of 681 overlapping DEGs between *Bambi^fl/fl^* and *Bambi^HSA-Cre^* mice. (E) KEGG pathway enrichment analysis of upregulated genes in TA muscle from *Bambi^fl/fl^* and *Bambi^HSA-Cre^* mice 30 days post-CTX-induced injury. (F) UCSC genome browser traces of ATAC-seq peaks at the promoter regions of the *Pax7*, *Myod1*, *Myog*, *Myf5*, and *Thbs4* loci. Identification of a conserved MyoD/MyoG binding site in the proximal promoter region of Thbs4. (G) Representative images of Pax7 (Red), Thbs4 (Green), and MyoD (Purple) immunostaining of the SCs (day 0) or cultured for 72-hr (day 3) in single myofiber (n>100 clusters from n=3 mice/group). (H) qPCR analysis of *Myod1*, *Myog* and *Thbs4* expression in TA muscle post CTX-induced injury on T0, T3, T5, T7, T9, T12 and T15 (n=6 mice). (I) Representative immunostaining images of Thbs4 (Green) and myogenin (Red) in C2C12 myoblasts at the proliferation (day 0) and differentiation stages (day 5). (J) Representative immunoblots of Thbs4 protein levels in C2C12 myoblast lysate and culture medium with either Ctrl or Thbs4-expressing adenovirus infection with/without BFA treatment at 0, 1, and 2 μg/ml. Data represent the mean ± standard error of the mean.

**Figure S10 for Figure 6.**
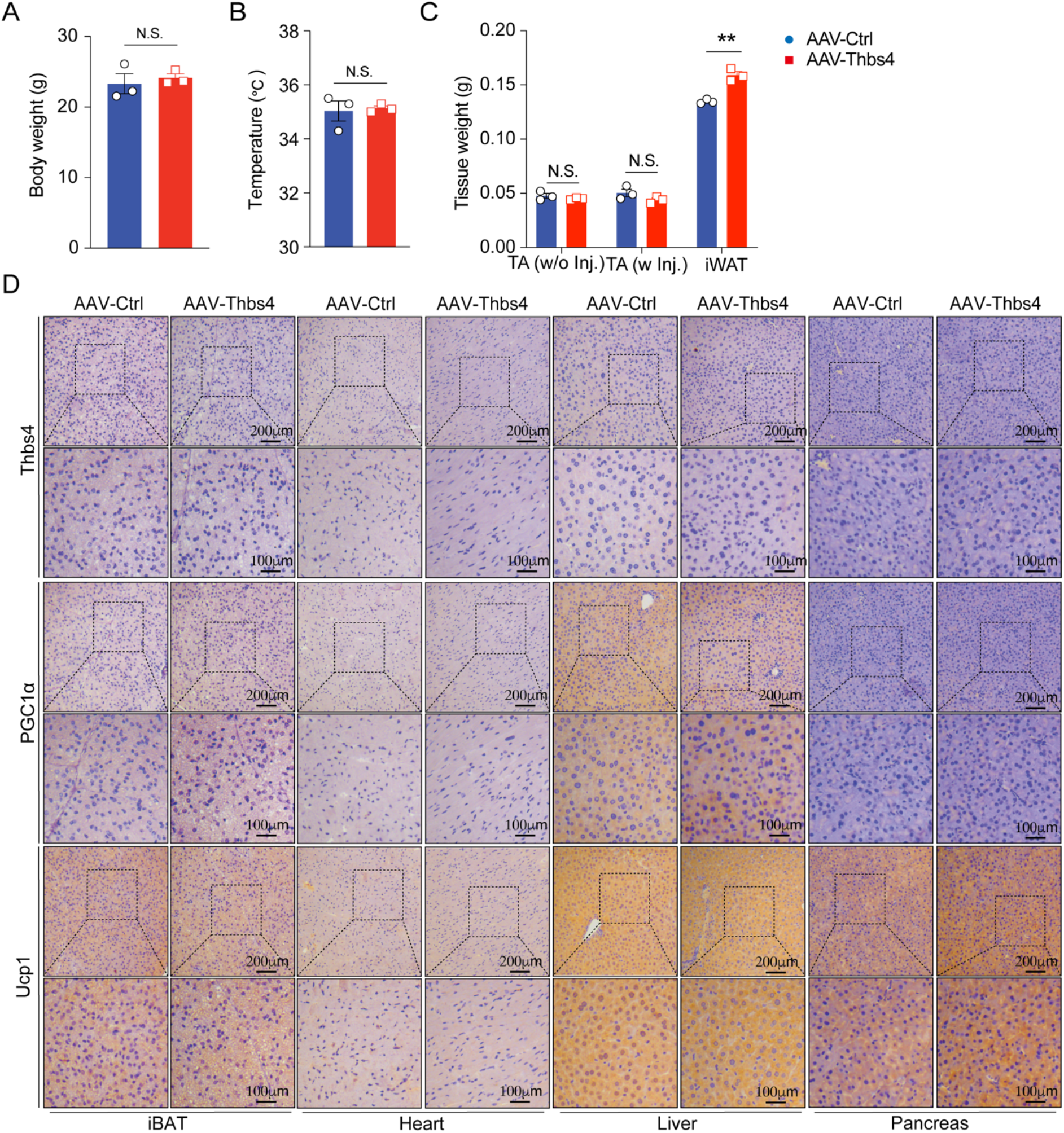
Thbs4 plays a long-term role in protecting against metabolic disorders. (A) Body weight of the AAV-Ctrl and AAV-Thbs4 groups (n=3 mice/group). (B) The rectal temperature of the AAV-Ctrl and AAV-Thbs4 groups. (C) The tissue weight of TA muscle and iWAT of the AAV-Ctrl and AAV-Thbs4 groups. (D) IHC of Thbs4 and Ucp1 in the section of iBAT, Heart, Liver and Pancreas of *C57BL6/J* mice of the AAV-Ctrl or AAV-Thbs4 groups. N.S., not significant, and \*\**p* < 0.01, by two-sided Student’s t test. Data represent the mean ± standard error of the mean.

**Figure S11 for Figure 6.**
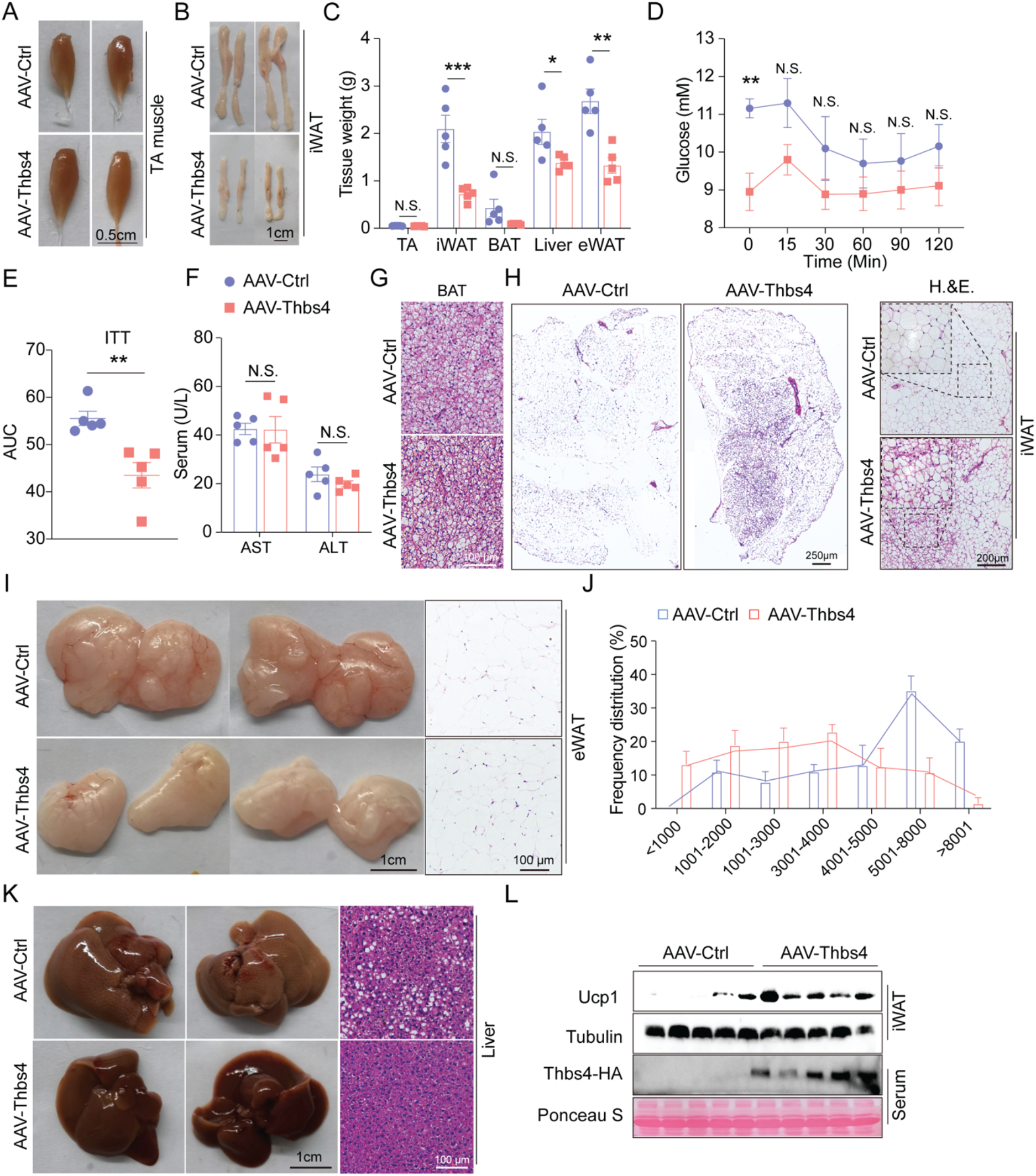
Thbs4 plays a long-term role in protecting against metabolic disorders. (A) Representative images of TA muscle of the AAV-Ctrl and AAV-Thbs4 groups under HFD feeding (n=5 mice/group). (B) Representative images of iWAT of the AAV-Ctrl and AAV-Thbs4 groups under HFD feeding. (C) Tissue weight of TA, iBAT, iWAT, eWAT and liver of the AAV-Ctrl and AAV-Thbs4 groups under HFD feeding. (D) ITT for AAV-Ctrl and AAV-Thbs4 groups under HFD feeding. (E) AUC of ITT of AAV-Ctrl and AAV-Thbs4 groups under HFD feeding. (F) Serum levels of AST and ALT of the AAV-Ctrl and AAV-Thbs4 groups under HFD feeding. (G) Representative images of H&E staining of iBAT sections of the AAV-Ctrl and AAV-Thbs4 groups under HFD feeding. (H) Representative images of H&E staining of iWAT sections, whole section images (left) and snap images (right) of the AAV-Ctrl and AAV-Thbs4 groups under HFD feeding. (I) Representative images of eWAT (left) and H&E staining (right) of the AAV-Ctrl and AAV-Thbs4 groups under HFD feeding. (J) Frequency distribution of the size of lipid droplets in eWAT of the AAV-Ctrl and AAV-Thbs4 groups under HFD feeding. (K) Representative images of liver (left) and H&E staining (right) of AAV-Ctrl and AAV-Thbs4 groups under HFD feeding. (L) Representative of immunoblots of Ucp1 and Tubulin of iWAT and Thbs4-HA of serum of the AAV-Ctrl and AAV-Thbs4 groups under HFD feeding. N.S., not significant, \**p* < 0.05, \*\**p* < 0.01 and \*\*\**p* < 0.001, by two-sided Student’s t test. Data represent the mean ± standard error of the mean.

**Figure S12 for Figure 7.**
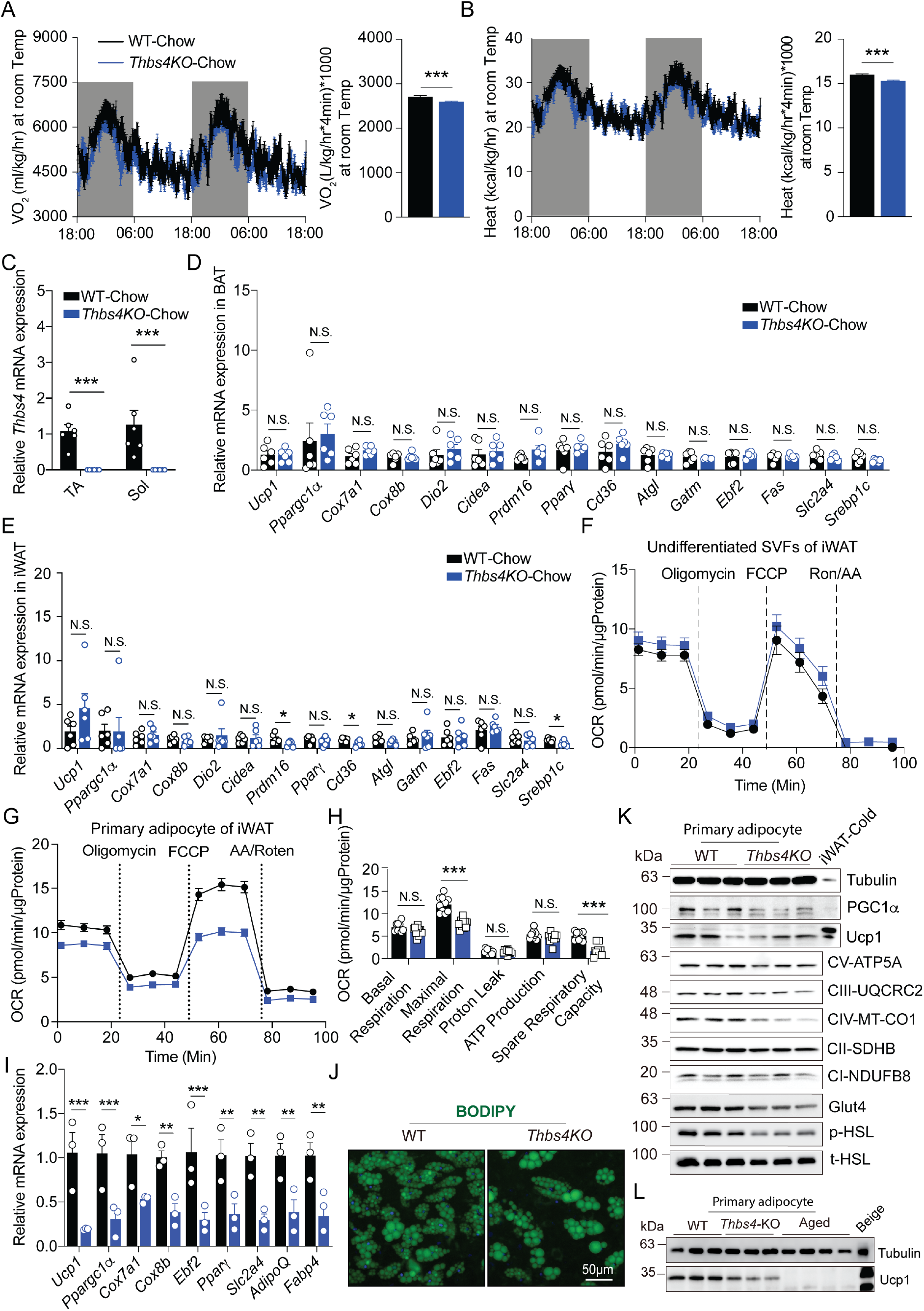
*Thbs4*-KO exacerbates HFD-induced metabolic syndromes. (A) Average VO_2_ (left) was monitored over 48-hr period and the AUC of VO_2_ (right) curve for WT and *Thbs4-*KO mice at R.T. (n=4 mice/group of two independent repeats). (B) Average VCO_2_ (left) was monitored over a 48-hr period and the AUC of VCO_2_ (right) curve for WT and *Thbs4-*KO mice at R.T. (n=4 mice/group of two independent repeats). (C) qPCR analysis of Thbs4 expression in TA and Sol muscle of WT and *Thbs4-*KO mice (n=6 mice/group). (D) qPCR analysis of thermogenesis-related gene expression in iBAT of WT and *Thbs4-*KO mice (n=6 mice/group). (E) qPCR analysis of thermogenesis-related gene expression in iWAT of WT and *Thbs4-*KO mice (n=6 mice/group). (F) Seahorse flux analysis of OCR of undifferentiated SVFs isolated from iWAT of WT and *Thbs4-*KO mice. (G) Seahorse flux analysis of OCR of differentiated primary white adipocyte of WT and *Thbs4-*KO mice. (H) Statistical analysis of OCR at the stage of basal respiration, maximal respiration, proton leaky, ATP production and spare respiration capacity. (I) qPCR analysis of thermogenesis-related gene expression of differentiated primary white adipocyte of WT and *Thbs4-*KO mice. (J) Representative image of BODIPY (Green) staining of differentiated primary white adipocyte. (K) Representative immunoblots of Ucp1, PGC1α, Glut4, p-HSL (Ser565), t-HSL and Mito-complex. (L) Representative immunoblots of Ucp1 and Tubulin of differentiated primary adipocyte from WT, *Thbs4*-KO and aged *C57BL6/J* mice. N.S., not significant, \**p* < 0.05, \*\**p* < 0.01, and \*\*\**p* < 0.001, by two-sided Student’s t test. Data represent the mean ± standard error of the mean.

**Figure S13 for Figure 7.**
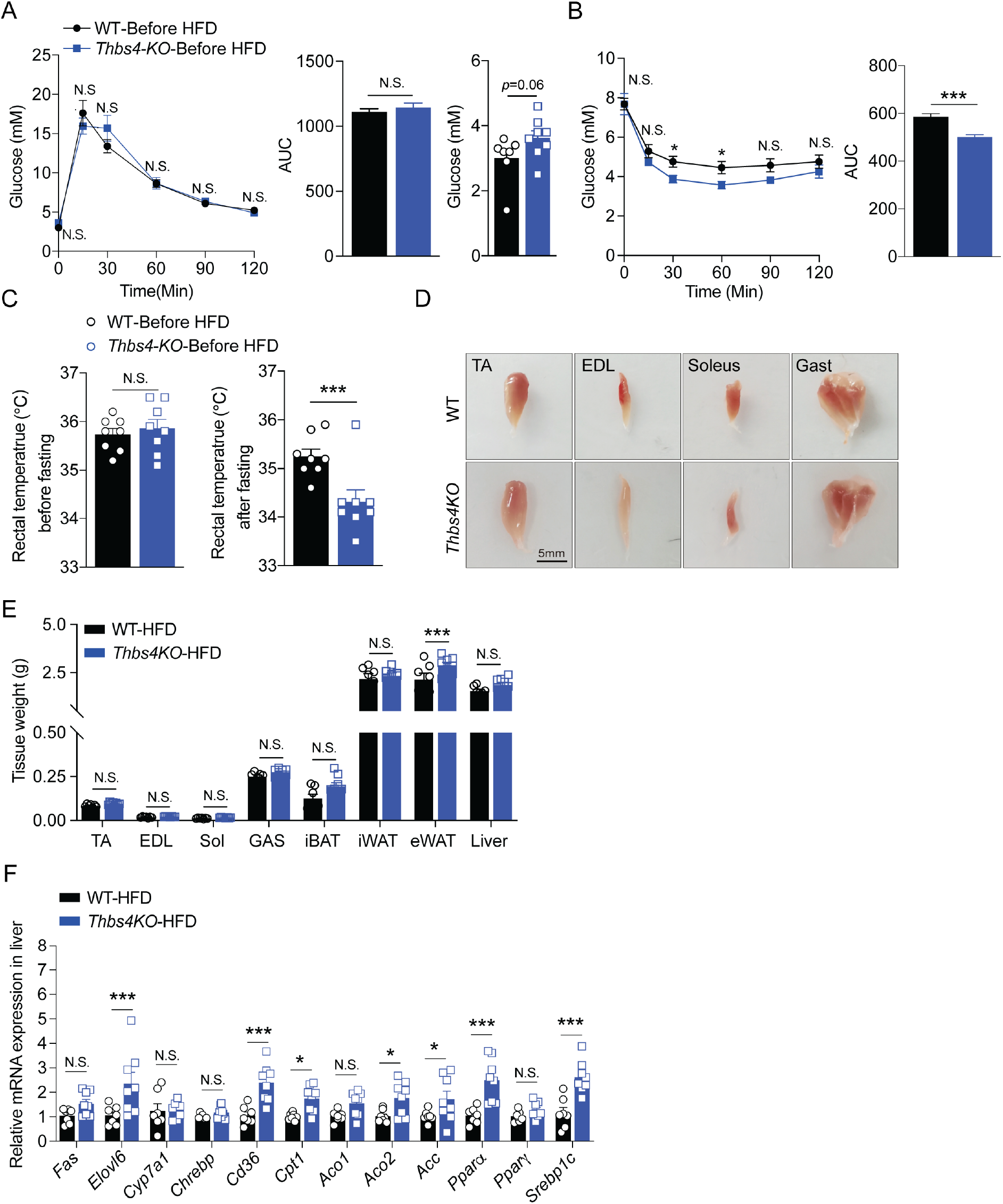
*Thbs4*-KO exacerbates HFD-induced metabolic syndromes. (A) GTT and initial fasting glucose level of WT and *Thbs4*-KO mice under HFD feeding (n=7-8 mice/group). (B) ITT of WT and *Thbs4*-KO mice under HFD feeding (n=7-8 mice/group). (C) The rectal temperature of WT and *Thbs4*-KO mice under HFD feeding before and after 16-hr fasting (n=7-8 mice/group). (D) Representative images of tissues (TA, Sol, EDL and Gas muscle) of WT and *Thbs4*-KO mice under HFD feeding (n=7-8 mice/group). (E) The tissue weight of TA, EDL, Sol, Gas, iBAT, iWAT, eWAT and liver of WT and *Thbs4*-KO mice under HFD feeding (n=7-8 mice/group). (F) qPCR analysis of lipid metabolism-related gene expression in liver of WT and *Thbs4*-KO mice under HFD feeding (n=7-8 mice/group). N.S., not significant, \**p* < 0.05, and \*\*\**p* < 0.001, by two-sided Student’s t test. Data represent the mean ± standard error of the mean.

**Figure S14.**
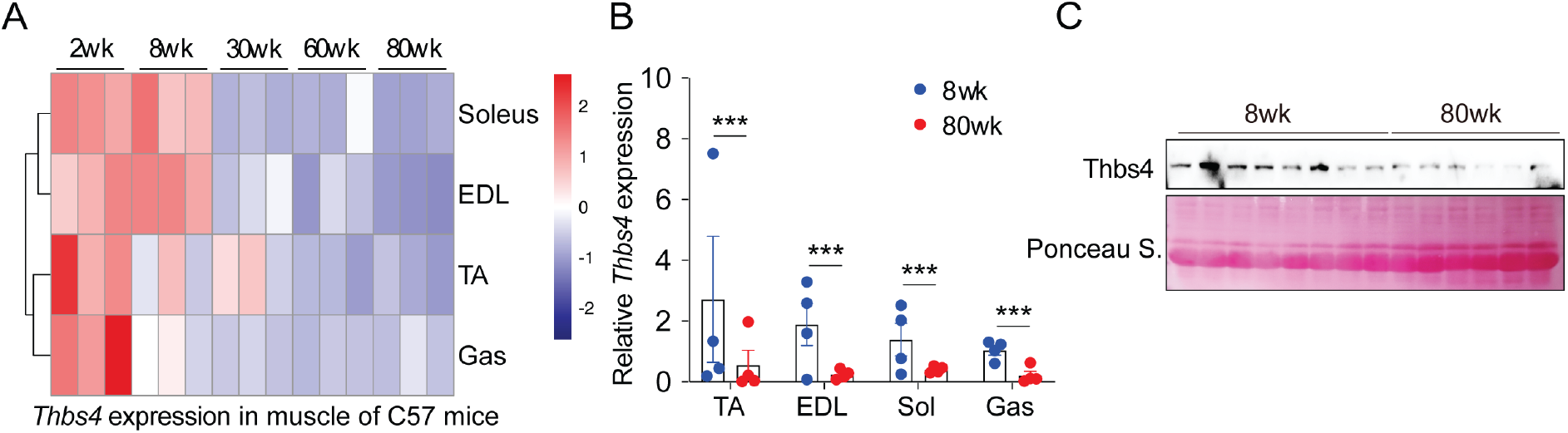
Thbs4 expression is decreased along with the progression of ageing. (A) Heatmap of the *Thbs4* expression profile in TA, EDL, Soleus and Gas muscle of *C57BL6/J* mice at different ages (2wk, 8wk, 30wk, 60wk and 80wk) (n=3 mice/group). (B) qPCR analysis of *Thbs4* expression in TA, EDL, Sol and Gas muscle of *C57BL6/J* mice between adult (8wk) and aged (80wk) (n=4). (C) Representative immunoblots of Thbs4 protein level in TA muscle of *C57BL6/J* mice between adult (8wk) and aged (80wk) (n=8). \*\*\**p* < 0.001, by two-sided Student’s t test. Data represent the mean ± standard error of the mean.

---

**Supplementary Table 1:**
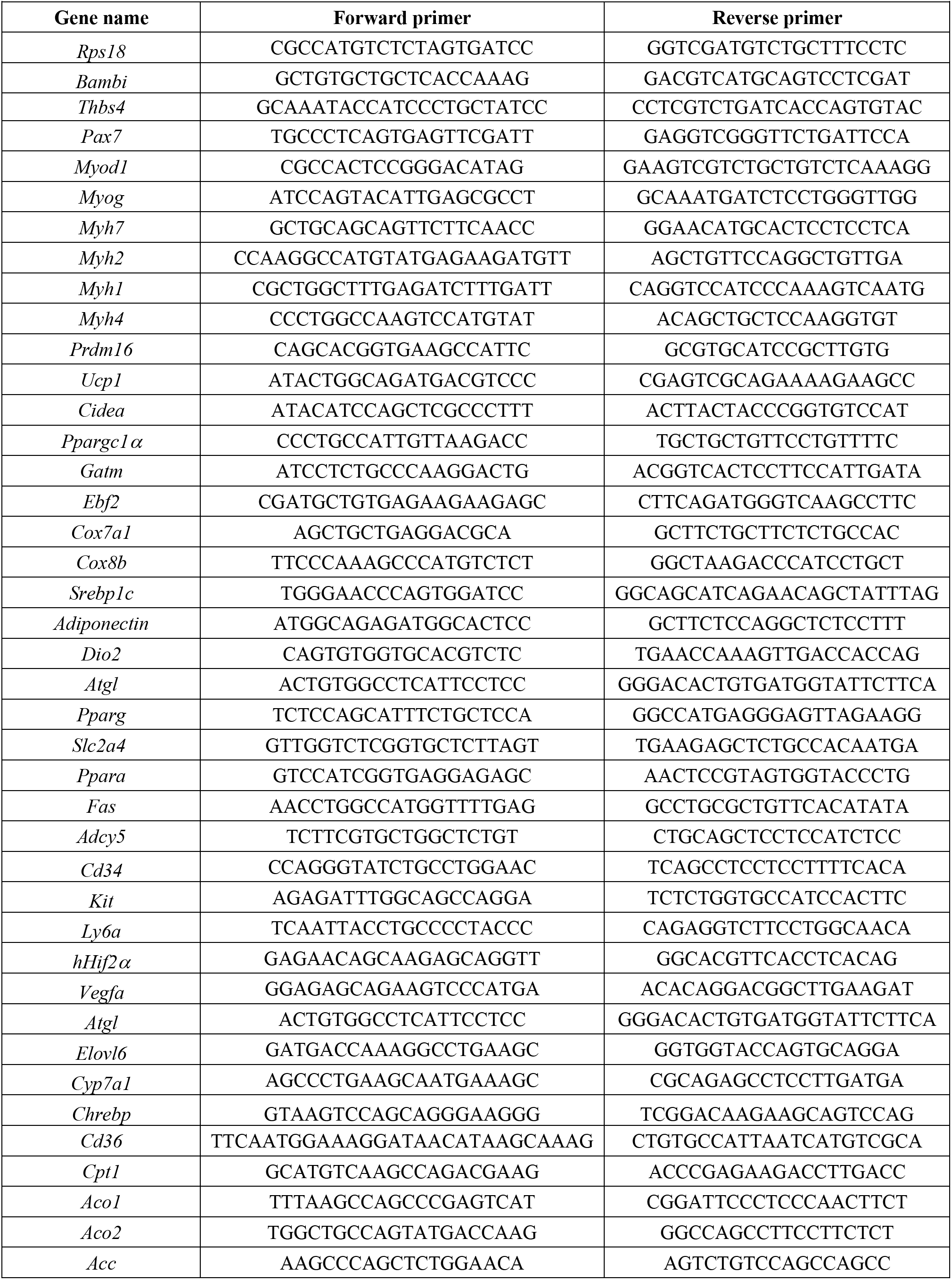
qPCR Primer list.

**Supplementary Table 2.**
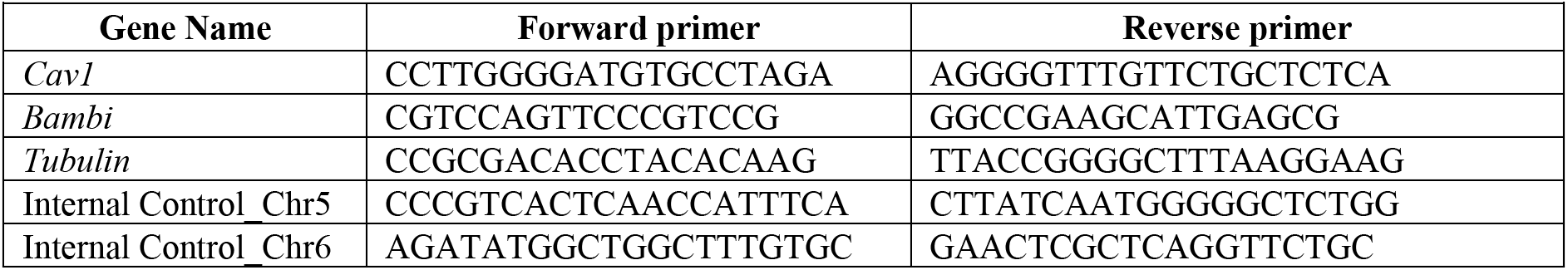
ChIP-qPCR Primer List.

**Supplementary Table 3.**
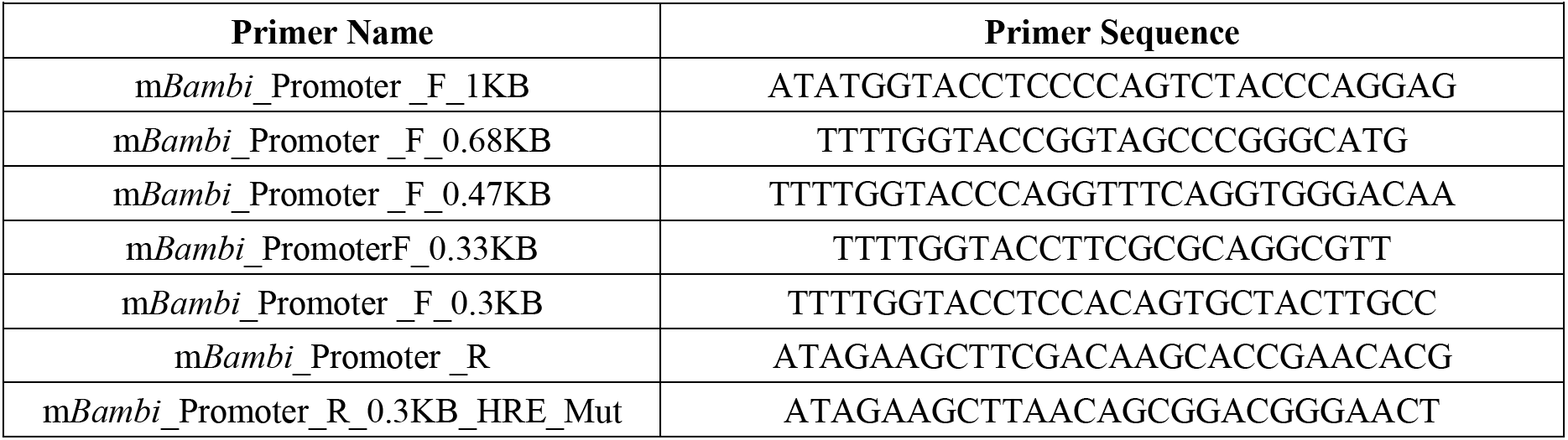
Cloning Primer List.

